# Deep-learning-assisted simulation of a cortical circuit: integrating anatomy, physiology and function

**DOI:** 10.64898/2026.03.13.711751

**Authors:** Shinya Ito, Darrell Haufler, Javier Galván Fraile, Kael Dai, Joseph Aman, Guozhang Chen, Claudio Mirasso, Wolfgang Maass, Anton Arkhipov

**Affiliations:** Allen Institute, Seattle WA, USA; IFISC, Universitat de les Illes Balears, Palma de Mallorca, Spain; School of Computer Science, Peking University, Beijing, China; Graz University of Technology, Graz, Austria

## Abstract

Mechanistic understanding of the brain requires models constrained by anatomy, physiology, and functional activity. We present a differentiable simulator and a ~67,000-neuron model of mouse primary visual cortex that integrates multimodal data, including electron-microscopy connectomics, multipatch synaptic physiology, cell-type-resolved intrinsic electrophysiology, and large-scale Neuropixels recordings from diverse cell types. End-to-end training completes on a single GPU in ~6.5 hours while preserving biological constraints. Networks trained only on brief drifting-grating responses reproduce cell-type-specific benchmarks and generalize to new contrasts and natural scenes. We uncover heterogeneous cell-type- and tuning-dependent synaptic organization and show that training preferentially sculpts inhibitory connectivity into distinct cohorts that exert outsized control over network activity. Targeted ablations show that removing biological priors on synaptic weight distributions can preserve functional activity yet disrupt emergent wiring rules. The freely shared models and code facilitate differentiable simulations as a computationally practical framework for studying brain circuit function and mechanisms under biological constraints.

## Main

Understanding brain mechanisms, computations, and diseases requires models that bridge synapse-level organization and cellular-resolution neural dynamics. Biorealistic circuit models pursue this by systematically integrating multimodal data on circuit composition, connectivity, and activity^1–7^. However, constructing such models remains challenging due to limitations in the availability of the necessary data. Moreover, they remain difficult to constrain at scale. Even when a model reproduces selected neural activity statistics, multiple parameterizations can yield similar outputs^8–10^, limiting identifiability and mechanistic interpretability. Reducing this degeneracy requires combining complementary constraints that restrict admissible parameter regimes and neural dynamics within a given circuit architecture.

Recent datasets are beginning to provide such constraints with cellular resolution across modalities and in multiple species and brain areas. In particular, extensive characterization of neuronal intrinsic electrophysiology, synaptic connectivity, and *in vivo* neural activity with unprecedented cell-type resolution in the mouse visual cortex creates unique opportunities for biorealistic modeling^7^. Systematic multimodal characterization of cortical cell types—through transcriptomics, electrophysiology, and morphological reconstruction—has yielded detailed taxonomies that serve as essential building blocks for constructing biorealistic models of mouse cortical circuits^11–17^. Large-scale synaptic physiology surveys have quantified cell-type-specific synaptic strength and temporal dynamics^18,19^. Surveys of *in vivo* activity across cortical layers and cell types provide robust targets for model evaluation and fitting^20,21^. More recently, electron microscopy-based functional connectomics has begun to link synapse-level anatomy to functional tuning within the same tissue, enabling direct tests of predicted structure–function relationships^22^.

Data-driven cortical models have started integrating subsets of these constraints. For example, a model of the mouse primary visual cortex (V1) by Billeh et al.^23^ combined diverse—though rather incomplete—anatomical and physiological measurements available at the time to build a large recurrent spiking circuit that reproduced multiple response benchmarks and yielded testable hypotheses about circuit organization^24–27^. A remaining key bottleneck is optimization: highly recurrent spiking networks can be unstable and historically required extremely time-consuming manual tuning to maintain realistic activity regimes^23^. Recently, differentiable simulation frameworks that enable gradient-based optimization of biological spiking networks against empirical objectives have started emerging^28–30^, shifting functional datasets from *post hoc* validation to active constraints on model development. Nevertheless, efficient automatic optimization of large-scale, heterogeneous, biorealistic brain circuit models remains challenging, both in terms of the computational resources required and in terms of creating suitable optimization objectives or loss functions.

Here we present a differentiable, GPU-accelerated simulation framework and a point-neuron recurrent spiking model of mouse V1 integrating electron microscopy (EM)-derived connectivity (MICrONS^22^ and V1 DeepDive^31^), multipatch synaptic physiology^19^, and large-scale *in vivo* recordings^20,21^ across all cortical layers and 19 cell types. Training the ~ 67,000-neuron network end-to-end takes ~ 6.5 h on a single GPU, and the resulting network simulates faster than real time. Models trained on a narrow set of drifting grating responses generalize to natural images and unseen contrast regimes, reproducing cell-type-specific firing rates, selectivity, and gain modulation. Analysis of the trained circuits reveals rich synaptic organization: excitatory connections follow like-to-like wiring rules consistent with connectomic data^32–34^, while inhibitory connections exhibit diverse pathway-specific structures. Training preferentially sculpts inhibitory synaptic organization into outgoing-weight cohorts that exert substantial control over network activity and selectivity despite their small size. Targeted constraint ablations show that biological priors on synaptic weight distributions, although not required to reproduce aggregate firing statistics, are essential for shaping circuit-level wiring motifs. All model code, trained parameters, and the simulation framework are publicly available.

### Model overview and data integration

The V1 model integrates multiple categories of data, including EM connectomics^22,31^, intrinsic neuronal electrophysiology^13,14^, multipatch synaptic physiology^19^, and large-scale *in vivo* recordings from the Neuropixels Visual Coding dataset^21^ (Fig. 1a). These data inform network initialization— covering neuron properties, connectivity, and synaptic parameters—producing a fully specified network in the SONATA format^35^ that preserves biological heterogeneity, including broad distributions of synaptic strengths and total input synapse counts across and within cell types (see Methods for details). Neuropixels data additionally provide cell-type-resolved firing-rate and selectivity targets for gradient-based optimization.

**Figure 1:**
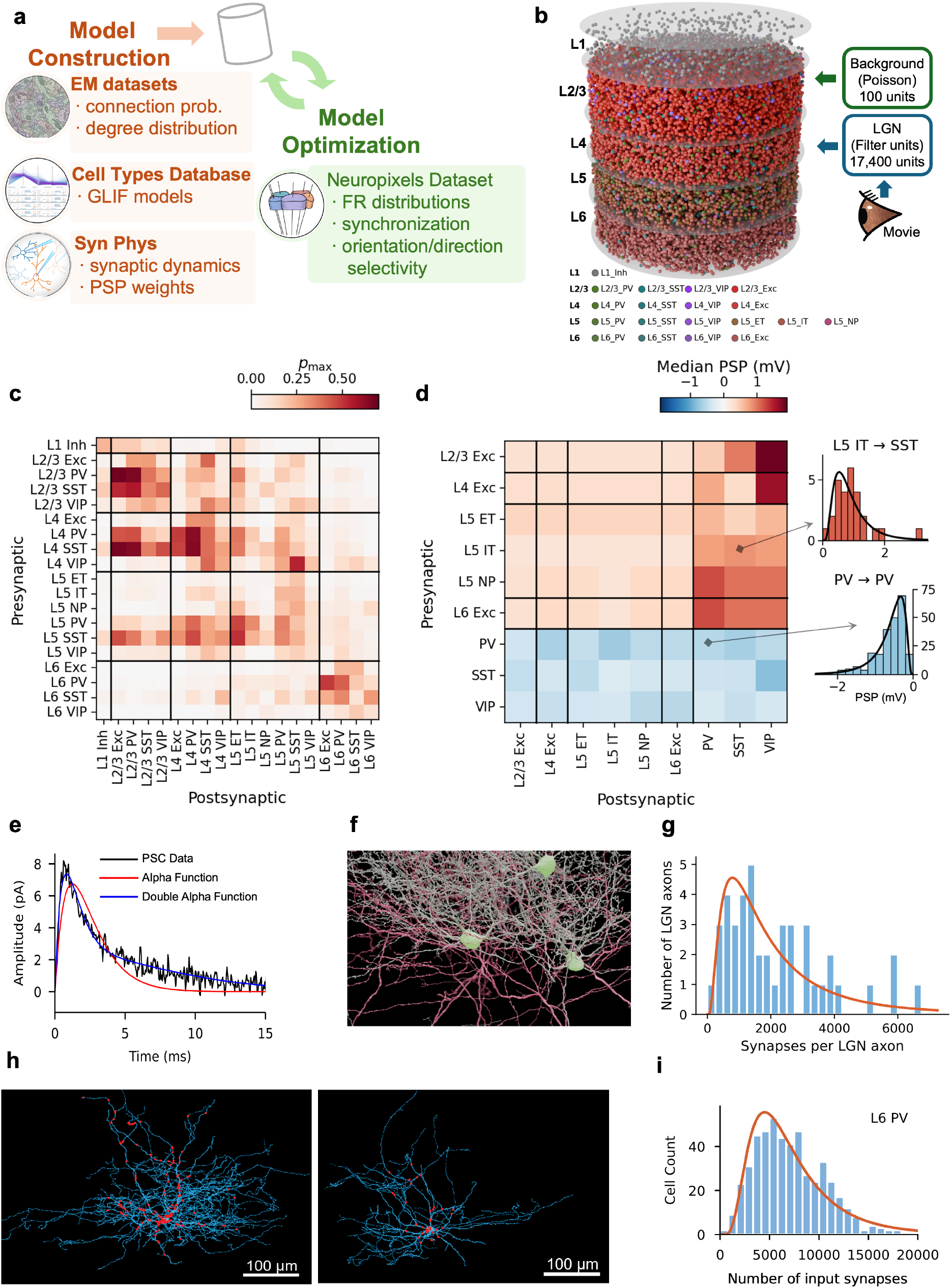
Model overview and data integration. Multimodal experimental data are integrated to parameterize a ~ 67,000-neuron spiking network model of mouse V1. **a**, Schematic of the major data sources integrated into the model. EM connectomics, intrinsic neuronal electrophysiology, and multipatch synaptic physiology inform model construction; large-scale Neuropixels recordings provide targets for model optimization. **b**, Three-dimensional rendering of the model. Individual neurons are colored by cell type; LGN (thalamocortical) and background inputs are indicated. **c**, Connection matrix across cell-type pairs showing the peak connection probability (*p*_max_). **d**, Postsynaptic potential (PSP) amplitudes across cell-type pairs derived from the Synaptic Physiology dataset^19^. Left, median PSP amplitudes; right, example PSP distributions and their maximum-likelihood log-normal fits. Top: L5 IT to SST (*N* = 24 connected pairs); bottom: PV → PV (*N* = 240). **e**, An example postsynaptic current (PSC) recorded in voltage clamp from a PV → L5 IT connection. Black, data; red, single-alpha function fit; blue, double-alpha function fit, which captures both fast and slow decay components. Connection-type-specific kinetic parameters were derived from median fits across quality-filtered recordings for each cell-type pair (see Methods). **f**, EM reconstruction showing three L4 basket (putatively PV) interneurons (green) contacted by two heavily branching LGN axons (red). **g**, Distribution of total synapse counts across 43 reconstructed LGN axons in EM data^22^. Bin width: 250 synapses. The orange line is a log-normal fit. **h**, EM reconstructions of two L6 basket (putative PV) interneurons (dendrites, blue) with synapses from other L6 basket cells (red). Left, a neuron with extensive dendritic arborization and numerous inputs from other L6 basket cells; right, a neuron with more compact morphology and fewer inputs. **i**, Distribution of total input synapse counts for L6 PV (basket) interneurons in EM data. Orange curve, maximum-likelihood log-normal fit.

We constructed 10 instances of recurrent spiking networks comprising 66,670 ± 240 neurons (mean ± s.d.) drawn from 19 cell types, including excitatory populations spanning layers 2/3 through 6 (with distinct L5 extratelencephalic (ET), intratelencephalic (IT), and near-projecting (NP) subtypes) and inhibitory populations (parvalbumin (PV)-, somatostatin (SST)-, and vasoactive intestinal peptide (VIP)-expressing neurons per layer, plus L1 inhibitory), represented by 201 point-neuron models fit to experimental data^13,23^. Neurons were placed within a cylindrical column (400 µm radius) at layer- and type-appropriate densities (Fig. 1b).

Connection probabilities, spatial spread parameters, and per-neuron target in-degrees were derived from EM data^22,31^, and synaptic transmission properties—postsynaptic potential (PSP) amplitude distributions and synaptic current kinetics—from multipatch recordings^19^ (Fig. 1c–e; see Methods). Synaptic current kinetics—fast and slow time constants and their amplitude ratio—were extracted by fitting double-alpha functions to voltage-clamp recordings for each connection type (see Methods). Recurrent connections were assigned using distance-dependent, orientation-dependent^36^, and like-to-like connectivity rules, with each neuron’s total input scaled to match log-normal in-degree distributions from EM data. The resulting peak connection probabilities and median PSP amplitudes are summarized in Fig. 1c, d, and the total number of connections is ~24.3 ± 0.2 million.

The MICrONS EM data^22^ also provided 43 reconstructed lateral geniculate nucleus (LGN) axons, revealing a broad, heavy-tailed distribution of per-axon synapse counts (Fig. 1f,g) as well as the distribution of per-connection synapse counts (Extended Data Fig. 1). In the model, a population of 17,400 LGN units generates spike trains through linear–nonlinear receptive-field filters^37,38^, with stronger thalamocortical connections assigned preferentially to spatially aligned sources (see Methods).

To assign synaptic weights, we accounted for within-cell-type variability in dendritic extent and total synaptic input (Fig. 1h) following the dendritic constancy rule^39^: neurons with larger in-degrees receive more, individually weaker connections, recovering the experimentally measured log-normal PSP distribution^40,41^ across the population (Fig. 1i; see Methods).

### Biologically constrained, scalable training of the cortical circuit model

After initialization, a brain circuit model requires optimization that ideally (i) preserves biological structure, (ii) fits multiple experimentally measured activity statistics simultaneously, and (iii) remains computationally practical (Fig. 2a). Typically, large recurrent spiking network models are either biologically detailed but difficult to optimize^30,42–44^ or trainable^45–48^ but omit substantial biological realism. To address this, we designed a GPU-based differentiable implementation and a multi-objective optimization framework, briefly described below (see Methods for details).

**Figure 2:**
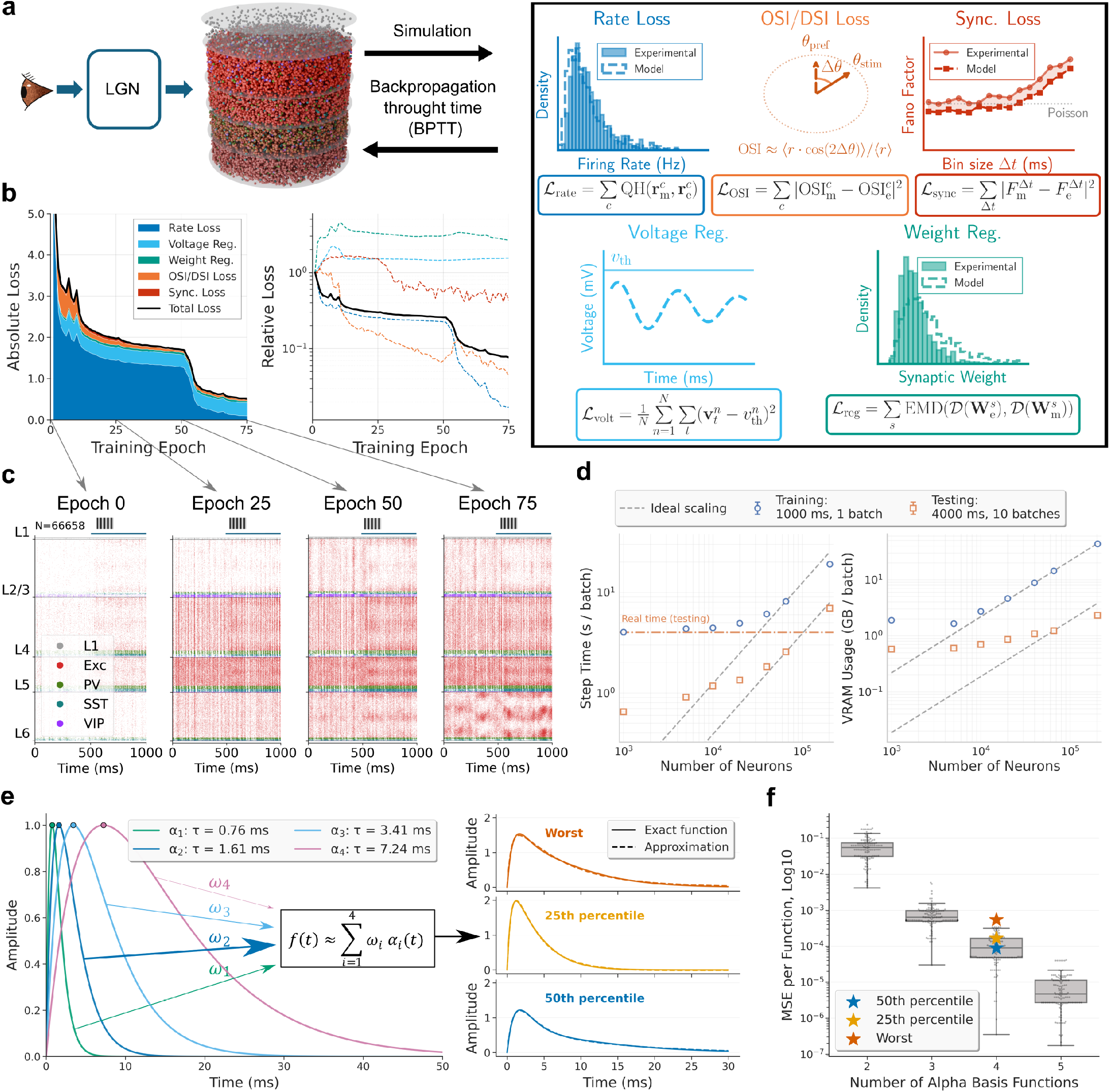
Biologically constrained, scalable training of the cortical circuit model. **a**, End-to-end training schematic. Visual input is processed by the LGN model and drives the recurrent V1 network. Network activity is compared with experimentally derived constraints through a multi-objective loss, including firing-rate distributions (evoked and spontaneous), orientation/direction selectivity, synchronization/variability statistics, voltage regularization, and synaptic-weight regularization. Gradients are propagated through time (BPTT) using surrogate-gradient approximations. See Methods for details. **b**, Training trajectories. Left: absolute loss components and total loss across epochs. Right: corresponding relative loss scales (log axis), showing coordinated optimization across objectives. **c**, Representative population rasters at initialization and during training (epochs 0, 25, 50, 75), organized by cortical layer and cell type, with neurons of each cell type vertically ordered by tuning angle. **d**, Computational scaling with network size under training and testing settings. The gray dashed lines show an ideal linear-scaling reference, anchored at *N* = 40, 000 neurons. Real-time performance for inference mode (“testing”; 4 real-world seconds for a 4-s-long simulation) is highlighted with a dashed orange line. **e**, Multi-alpha postsynaptic current (PSC) kernel approximation. Left: four shared basis functions (τ = {0.76 *ms*, 1.61 *ms*, 3.41 *ms*, 7.24 *ms*}). Right: reconstructed double-alpha kernels for representative 50th-percentile, 25th-percentile, and worst-case fits. **f**, Basis size versus approximation error. Box plots show MSE distributions across fitted double-alpha kernels. Error decreases sharply with basis size; with four basis functions, all fits satisfy MSE < 10^−3^. Stars indicate the 50th-percentile, 25th-percentile, and worst-case examples shown in panel (e).

Our simulator preserves sparse delayed connectivity, detailed synaptic dynamics, and fixed excitatory/inhibitory sign structure (Dale’s law^49^). For gradient propagation through non-differentiable spike generation we use BPTT with surrogate gradients^50,51^. We introduced an *Exponentiated Adam* optimizer that combines Adam-style adaptive moments^52^ with sign-preserving multiplicative updates^53–56^, naturally respecting Dale’s law and the heavy-tailed synaptic weight statistics of the cortex. GPU-level optimizations include sparse event-driven computation, mixed precision^57^, selective just-in-time (JIT) compilation, gradient checkpointing^58^, and parallel execution.

What target should one use for training? One approach is to assume a specific function for the trained network, such as motion detection^59^; however, it is unlikely that a single computation captures the full role of mouse V1^60–66^. We therefore chose to match neural responses experimentally recorded *in vivo*. In the Neuropixels data we use^21^, responses to drifting gratings (DGs) are well represented. Though an artificial stimulus, DGs provide well-established metrics such as orientation and direction selectivity indices (OSI/DSI), whereas responses to natural images or movies may require more sophisticated analyses. We therefore used DG-evoked responses, along with spontaneous activity (gray-screen condition), as training targets. This choice, also made to control computational cost, is justified *post hoc* by good generalization of the trained networks across stimulus classes (see below).

At each optimization step, visual stimuli were converted to LGN spikes and propagated through the recurrent V1 circuit, and the resulting activity was compared to Neuropixels-derived benchmarks^21^ (Fig. 2a). Optimization adjusted recurrent (V1→V1) and background synaptic weights, while thalamocortical (LGN → V1) weights were held fixed (see Methods). Each step used two 500-ms stimulus conditions, gray screen and DGs, with batch size 5 for each condition. Agreement with *in vivo* data was measured using per-cell-type firing-rate distributions, OSI/DSI, and excitatory population multi-scale synchrony (via Fano factor). To make OSI/DSI tractable over the short 500-ms BPTT chunks used for training, we developed a *crowd-surrogate OSI/DSI loss* (see Methods). Synaptic weight distributions per pre–post cell-type pair were regularized to preserve biologically observed distributions^19^ while permitting individual synaptic changes to achieve the training targets.

The training trajectories in Fig. 2b show coordinated reduction of all loss components across epochs, with no evidence of runaway instability (Extended Data Fig. 2). Recurrent spiking models are often susceptible to pathological regimes (runaway excitation or near-complete suppression) during gradient-based training^67,68^, but we found that the initializations of our models from biological data were already stable, and training progressively retuned the circuit toward experimentally matched operating points. Representative raster plots (Fig. 2c) illustrate stable activity that shifts during training toward realistic firing levels and selectivity across layers and cell types.

Computational scaling (Fig. 2d) shows approximately linear growth with network size; inference remained faster than real time for networks below 100,000 neurons, and a batch-size sweep confirms near-linear VRAM growth (Extended Data Fig. 3). On a single NVIDIA RTX PRO 6000 GPU, each 1 s training step required ~ 11.8 s wall-clock time and full optimization took ~ 6.5 h. To control BPTT memory cost without discarding PSC diversity, we approximated the fitted double-alpha kernels using a shared 4-function alpha basis (Fig. 2e,f), achieving MSE < 10^−3^ for all connection types and reducing PSC state storage by 5.5× (from 44 to 8 states per neuron; see Methods).

Together, these developments enabled end-to-end training of biorealistic networks with hundreds of thousands of neurons and tens of millions of synapses on a single GPU (see Methods).

### Trained models reproduce physiological responses across cell types

We first evaluated trained networks using DGs—the same stimulus type used for training, but with longer presentations (2 s vs. 0.5 s in training) across 8 canonical drift directions (Fig. 3a,b). This tests robustness to longer stimuli, matching the duration of experimental recordings^21^, and covers the standard visual response properties used in the field.

**Figure 3:**
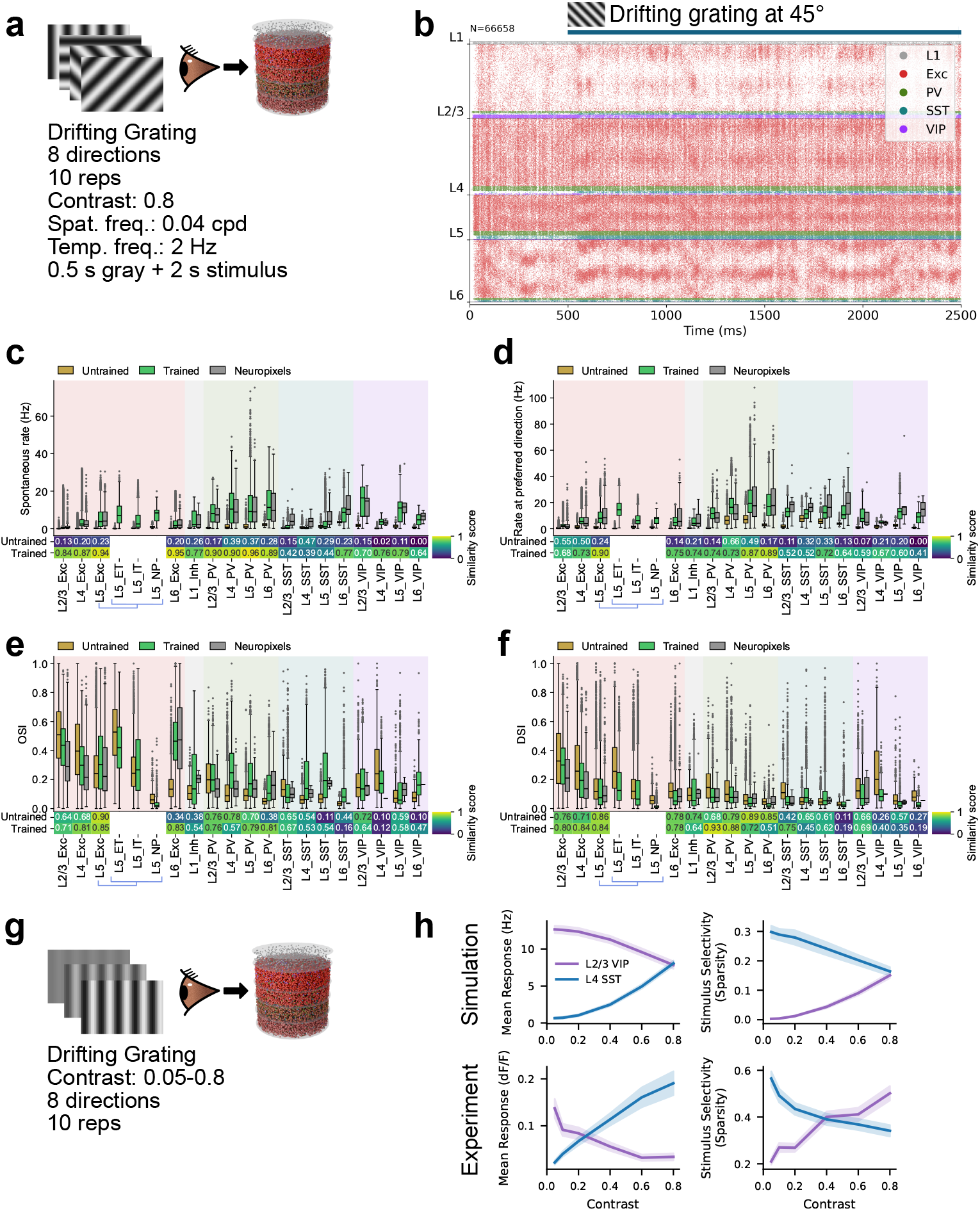
Trained models reproduce physiological responses across cell types. **a**, Schematic of the simulations with the DG stimuli. **b**, Raster plot of the activity of a trained network (sorted by cell type, then by the tuning angle). **c-f**, Quantification of response properties, computed for each cell and grouped by cell type. Trained and untrained models (*N* = 10 individually trained models each) are compared with experimental Neuropixels recordings *in vivo*. Similarity scores between the models and experiments (1 − *D*_KS_, where *D*_KS_ is the Kolmogorov-Smirnov statistic) are shown below the plots. Note that both aggregated and segregated results for L5 excitatory types are shown, but only aggregated results are available in the Neuropixels data. **c**, Spontaneous rates (i.e., gray screen). **d**, Firing rates to preferred direction. **e**, Orientation selectivity index (OSI). **f**, Direction selectivity index (DSI). **g**, Contrast-response simulations. **h**, Mean response and stimulus selectivity as a function of contrast for L2/3 VIP and L4 SST.

As described above, the biological structure priors yielded stable network dynamics already at initialization, and stability is maintained throughout training (Fig. 3b; Extended Data Fig. 2). Notably, this robust operating regime generalizes to the longer 2 s stimuli used here for evaluation, compared to the 0.5 s trials used during training.

Fig. 3c,d shows that training greatly improved the match with experimental data for both spontaneous and evoked firing rates across cell types, with median similarity scores increasing from 0.21 to 0.80 for spontaneous and from 0.21 to 0.70 for evoked firing rates. OSI distributions (Fig. 3e) were also better matched after training (median score: 0.56 to 0.64). DSI (Fig. 3f) was already close to experimental distributions before training, but at much lower firing rates; training brought firing rates to physiological levels while preserving selectivity.

Training used only high-contrast DGs, yet the trained models generalized well to lower contrasts (Extended Data Fig. 4). A contrast-dependent dichotomy between SST and VIP populations in the upper layers emerged naturally—consistent with experimental data^69^—as did a contrast-dependent decrease in selectivity for SST neurons (Fig. 3g,h).

### Trained models generalize to natural images

We next tested whether the models trained on the artificial DG stimuli generalized to the more ethologically relevant and complex natural-image inputs. We first used the natural scenes from the “Brain Observatory” (BO) set (Fig. 4a). While relatively small (118 images), this set is important in that it was used for extensive Neuropixels recordings in mouse V1^21^, offering an excellent quantitative benchmark. The BO natural images evoked neural responses that were consistent in their magnitude between the trained models and Neuropixels data across cell types, showing an improvement over the untrained models (median similarity score: 0.24 for untrained vs. 0.65 for trained; Fig. 4b). Selectivity of neurons to image presentations was consistent as well (median similarity score: 0.32 for untrained vs. 0.58 for trained; Fig. 4c).

**Figure 4:**
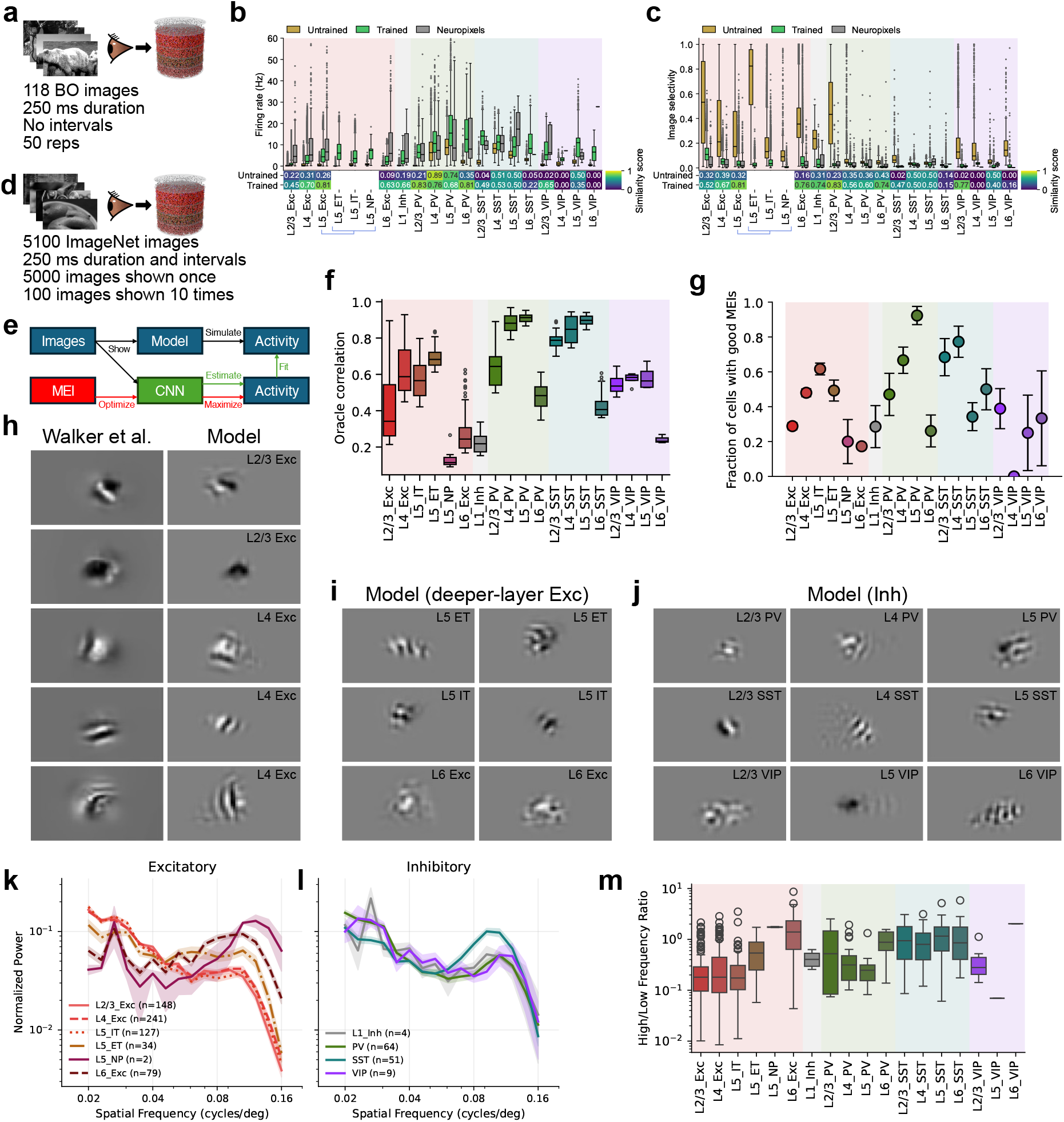
Trained models generalize to natural images. **a**, Brain Observatory (BO) natural scenes stimulus, consisting of 118 images. **b–c**, Firing rates (b) and image selectivity (c) for the BO stimulus, compared between trained/untrained models and experiment. **d**, ImageNet stimulus. Images were presented for 250 ms and separated by 250 ms of gray screen. A total of 5000 images were shown once, and 100 images were shown 10 times to calculate response reliability. **e**, Most exciting input (MEI) optimization schematic. **f**, Oracle correlation (a measure of response reliability) for each cell type. **g**, Fraction of cells that produced good MEIs. **h–j**, Example MEIs for upper-layer excitatory neurons (including comparisons with MEIs obtained using experimental data^70^ available for these populations), deeper-layer excitatory, and inhibitory neurons. **k–l**, Normalized spatial power spectrum of the MEIs for excitatory (k) and inhibitory (l) cell types. **m**, Distributions of the ratio between the high frequency power and the low frequency power of the MEI spatial power spectrum for each cell type.

To probe the features that drive individual model neurons, we presented ImageNet images (5000 once, 100 repeated; Fig. 4d) and generated most exciting inputs (MEIs)^70^ by fitting a CNN to each neuron’s responses and optimizing stimuli to maximize predicted firing (Fig. 4e; see Methods). Response reliability was quantified via oracle correlation (Fig. 4f), and the fraction of cells yielding stable MEIs was computed per cell type (Fig. 4g). Oracle correlations and MEI quality differed systematically across populations, reflecting cell-type-specific differences in circuit organization.

In particular, L5 NP cells had the lowest oracle score, indicating limited image-dependent response reliability, and yielded fewer spatially confined MEIs (Fig. 4f,g). This differs from L5 ET and IT, despite shared DG training targets for all L5 excitatory populations, suggesting that differences arise from cell-type-specific wiring. We confirmed this model prediction by analyzing cell-type-labeled neurons with visual physiology data in the MICrONS EM dataset, where L5 NP cells also showed lower oracle scores than L5 ET and IT cells (Extended Data Fig. 5a). Furthermore, the EM data show circuit differences that may explain these functional distinctions: the in-degree distribution of NP cells is markedly distinct from those of IT and ET cells (log-normal shape parameters: IT 0.331, ET 0.424, NP 1.376; scale parameters: IT 2552, ET 7407, NP 680), indicating that NP cells receive far fewer but more variable synaptic inputs; NP cells also receive a smaller fraction of their excitatory input from L2/3 and L4, and proportionally more from L5 ET and L6 (Extended Data Fig. 5b), suggesting less responsiveness to feedforward visual inputs.

We found that MEIs for upper-layer excitatory neurons exhibited localized structure that often included smooth stripe patterns, with many examples closely resembling MEIs previously reported for experimentally recorded neurons from the same populations in mouse V1 (Fig. 4h)^70^. In deeper excitatory populations, MEIs from our network models showed more complex spatial patterns (Fig. 4i) along with the stripes; inhibitory neurons exhibited similarly rich structure across all layers (Fig. 4j). To quantify these differences, we computed the spatial power spectra of the MEIs for excitatory (Fig. 4k) and inhibitory (Fig. 4l) classes and summarized each neuron’s spectral profile by a high (0.08–0.16 cycles per degree (cpd)) vs. low (0.02–0.04 cpd) spatial-frequency ratio (Fig. 4m). This analysis revealed a systematic shift toward higher-frequency content in deeper-layer excitatory neurons, whereas inhibitory neurons exhibited similarly rich structure across layers, consistent with high spatial complexity in the MEIs for these populations.

Thus, the statistics of trained models’ responses to natural images matched experimental data, and their MEI-derived response characteristics were consistent with experiments where data were available (i.e., upper-layer excitatory neurons). Beyond these validated areas, the MEIs revealed layer- and cell-type-dependent differences in preferred features, which serve as predictions for future experimental testing.

### Training yields tuning-dependent synaptic structure and reveals inhibitory cohorts with distinct functional roles

Having established that the trained models exhibit realistic response statistics across stimulus families, we next asked what synaptic patterns and circuit-level mechanisms emerge in these models. We characterized synaptic weight structure using two complementary functional axes: similarity of stimulus-evoked responses and similarity of the preferred direction of motion in a connected pair of neurons. Note that presence or absence of connections is not modified in our training method (see Methods), meaning that the effects described below are in addition to the existing connectivity structure.

#### Relation of synaptic weights and response correlations

Like-to-like connectivity has been studied extensively for excitatory neurons^32–34,71^, but organization principles for inhibitory connections and across layers remain debated^72–74^. Our models offer an opportunity to study these functional wiring rules for all cell types at scale.

We computed response correlations (see Methods) under the BO natural scenes (Fig. 5a) and observed a substantial diversity of weight-vs.-correlation relations across source–target type pairs (Fig. 5b,c). While prior studies focused primarily on connection probability rather than weights^32,34,71,75^, we found that synaptic strengths in our models also exhibited clear functional logic. Generally, excitatory connections are dominated by like-to-like relations, where pairs with similar stimulus responses have stronger weights (e.g., L2/3 Exc → L6 Exc; Fig. 5b). Inhibitory connections are notably more diverse: some pathways follow anti-like-to-like relations (stronger weights for negatively correlated pairs, such as L5 PV → L5 IT and SST → VIP) or more complex non-monotonic patterns (e.g., L5 PV → L5 ET and VIP → PV) (Fig. 5b; see full matrix in Extended Data Fig. 6).

**Figure 5:**
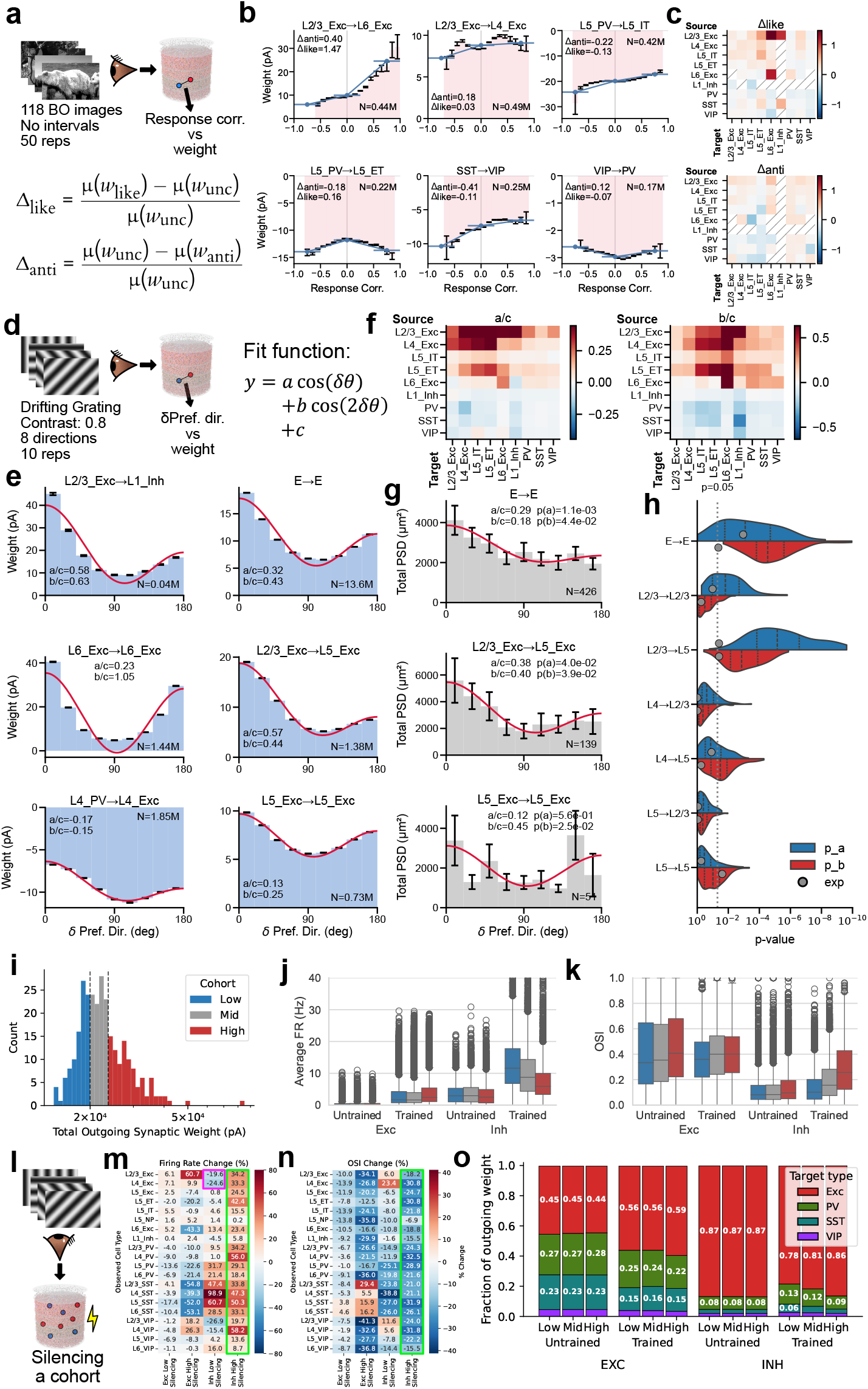
Training yields tuning-dependent synaptic structure and reveals inhibitory cohorts with distinct functional roles. Analyses shown aggregate data across *N* = 10 individually trained models. **a**, Schematics of the stimulus and analysis of weight as a function of response correlation. **b**, Examples of diverse relations between synaptic weight and response correlation of connected neuron pairs in the trained network models. Number (N) of pairs analyzed is indicated. **c**, Heatmaps of Δ_like_ and Δ_anti_ for a subset of cell-type pairs; matrix elements involving PV, SST, and VIP types exclude inter-layer connections. See Extended Data Fig. 6 for the full matrix including intra- and inter-layer connections. For both heatmaps, positive values correspond to like-to-like and negative to anti-like-to-like relations. When neuron pairs with sufficient correlation (< − 0.5 or > 0.5) were not found, these metrics were not calculated and a diagonal line is placed in the heatmap element. **d**, Schematics of the stimulus and analysis for weight changes as a function of preferred-direction difference. **e**, Examples of relations between synaptic weight and preferred-direction difference. **f**, Heatmaps of cosine-fit effect sizes (a/c: direction like-to-like component, b/c: orientation like-to-like component) for a subset of cell-type pairs; matrix elements involving PV, SST, and VIP types exclude inter-layer connections. See Extended Data Figs. 7 and 8 for the full matrices including intra- and inter-layer connections. **g**, Electron microscopy experimental data (V1DD) showing the relation between preferred-direction difference and post-synaptic density (PSD). **h**, Statistical comparison between V1DD data (gray) and Monte Carlo sampling from the trained models when the same number of connections as available in V1DD data were chosen (red and blue). **i**, Example total output synaptic weight distribution (L4 PV), split into low, mid and high outgoing-weight cohorts. **j**, Firing rates (pre- vs. post-training) for each cohort. **k**, OSI (pre- vs. post-training) for each cohort. **l**, Schematic for silencing simulations. **m**, Change in firing rates during silencing. The purple rectangle highlights paradoxical inhibition of the L2/3 and L4 excitatory populations when low-outgoing-weight inhibitory neurons are silenced. The green rectangle highlights a broad increase in firing rate when high-outgoing-weight inhibitory cohorts are silenced. **n**, Change in OSI during silencing. The green rectangle highlights a broad suppression of the selectivity when high-outgoing-weight inhibitory neurons are silenced. **o**, Fraction of outgoing synaptic weights to different cell types for each cohort.

To summarize these effects, we computed metrics (Δ_like_ and Δ_anti_) that capture how strongly synaptic weight increases on the positively and negatively correlated sides, respectively (Fig. 5a,c; see Methods). This analysis revealed a general trend for like-to-like connectivity: excitatory neurons tended to form like-to-like connections regardless of the recipient cell type, and this pattern held for both positively and negatively correlated pairs. This tendency was more pronounced in projections from superficial to deeper layers. In contrast, inhibitory connections were more diverse, exhibiting both like-to-like and anti-like-to-like motifs, and frequently switching signs between positive and negative correlation regimes. These results demonstrate that the mapping from correlation to inhibitory synaptic strength is complex, non-monotonic, and pathway-dependent, reflecting the intricate interplay between interneuron types.

#### Relation of synaptic weights and preferred-direction differences

Previous experimental studies suggest that preferred-direction similarity predicts connection probability in V1^32–34,71,75^ but not strength for L2/3 E-to-E connections^33^. We measured the dependence of synaptic weight on preferred-direction difference for all cell types and quantified effects using a cosine-series fit (Fig. 5d–f; see Extended Data Fig. 7). This revealed widespread like-to-like organization for excitatory connections and anti-like-to-like for inhibitory connections (prime examples in Fig. 5e), with increasing like-to-like tendencies from superficial to deeper layers for excitatory pathways.

To directly compare model predictions to anatomy, we used the V1DD EM data^31^ linking functional tuning differences to post-synaptic density (PSD) as a proxy for synaptic strength. Like-to-like relations between synaptic weight and preferred direction reached statistical significance for excitatory cells in aggregate, L2/3 Exc→L5 Exc, and L5 Exc→L5 Exc connections (Fig. 5g). Matching sample sizes between model and experiment via Monte Carlo resampling (Fig. 5h; see Methods) showed that the available EM dataset afforded sufficient power for only a subset of pathways; for those, model predictions were confirmed. L2/3 E-to-E connections are a notable exception where the like-to-like effect is expected to be weak even in models, consistent with previous work^33^. Overall, the models show specific structure not only for correlation but also for direction similarity, which is generally like-to-like for excitatory and anti-like-to-like for inhibitory connections. Experimental data support this where sampling is sufficient, and future experiments should resolve this further.

#### Synaptic weight heterogeneity and its functional correlates

Cortical circuits exhibit heavy-tailed synaptic weight distributions^41,76^ that may support hub-like dynamics^77–79^. Our trained models showed approximately log-normal outgoing weight distributions (Fig. 5i). Within each of the 19 cell types, we split neurons in the 200 µm analysis core into low-, mid-, and high-outgoing-weight tertiles and compared firing rates, selectivity, downstream targeting, and network-wide impact under cohort-specific silencing (Fig. 5j–o; see Methods).

We did not observe strong cohort-specific differences in firing rates or OSI in the excitatory populations, suggesting that the functional roles of the excitatory cells are not segregated by the amount of their output even after training. On the other hand, the inhibitory populations showed prominent cohort-specific differences after training. In the trained networks, firing rates differed systematically across inhibitory cohorts (Fig. 5j), and the same cohorts showed structured differences in OSI (Fig. 5k), consistent with the idea that subsets of inhibitory neurons can acquire distinct functional profiles. These cohort differences were attenuated or absent in the untrained network, suggesting that high-outgoing-weight inhibitory phenotypes emerged as part of the learned circuit solution rather than being a trivial consequence of initialization. Such training-dependent changes in functional properties were not observed for cohorts separated by the incoming connections (Extended Data Fig. 9). Therefore, we focused on characterizing the relation between functional properties and outgoing weights.

#### Cohort-specific silencing reveals network-wide impacts

We performed cohort-specific silencing simulations (Fig. 5l, see Methods) and quantified the resulting changes in firing rate and OSI across the remainder of the network. Silencing inhibitory cohorts produced strong network-wide effects (Fig. 5m–n) despite the cohorts’ small size (~1% of all neurons: ~ 12% inhibitory × 1/3 tertile × 1/4 core-area fraction; see Methods). In particular, silencing high-outgoing-weight inhibitory neurons led to a widespread increase in firing rates and suppression of OSI across populations (green rectangles), whereas silencing low-outgoing-weight inhibitory neurons produced paradoxical suppression^80,81^ of firing in L2/3 and L4 excitatory populations (purple rectangle). These results suggest that inhibitory heterogeneity, when organized along outgoing-weight axes, can create distinct regimes of circuit control.

Inhibitory cohorts also differed in targeting (Fig. 5o): high-outgoing-weight neurons preferentially targeted excitatory populations, while low-outgoing-weight neurons preferentially targeted inhibitory ones (a disinhibitory pattern), providing a possible mechanism for the paradoxical suppression observed when silencing the low-weight cohort.

Cohort effects were consistent across inhibitory subtypes (Extended Data Fig. 10): PV cohorts had the strongest impact, SST cohorts showed laminar specificity, and VIP cohorts produced weaker effects. Together, these simulation results (consistent with recent findings in piriform cortex^82^) suggest that cohort identity—defined by outgoing weight rather than transcriptomic type—captures a continuous axis of inhibitory heterogeneity. This prediction offers a potentially fruitful direction for future experiments in V1.

### Effect of biological weight constraints

We next asked how strongly circuit organization depends on the biological constraint on synaptic weight distributions from multipatch physiology^19^—a non-trivial test, since unconstrained models may still satisfy population-level activity targets. We trained *N* = 10 models with the weight-distribution regularization term removed and compared the resulting circuit organization to constrained models discussed above.

The unconstrained models remained stable and reproduced firing-rate and selectivity patterns from Neuropixels data (Fig. 6a–c; Extended Data Fig. 11), with similarity scores consistent with constrained models (median peak firing rate: 0.70 vs. 0.73; OSI: 0.64 vs. 0.68). This indicates that functional objectives alone can guide the network to a satisfactory operating regime; however, it also implies that multiple synaptic configurations satisfy the same activity targets, making the constrained–unconstrained comparison informative about which aspects of organization require biological priors.

**Figure 6:**
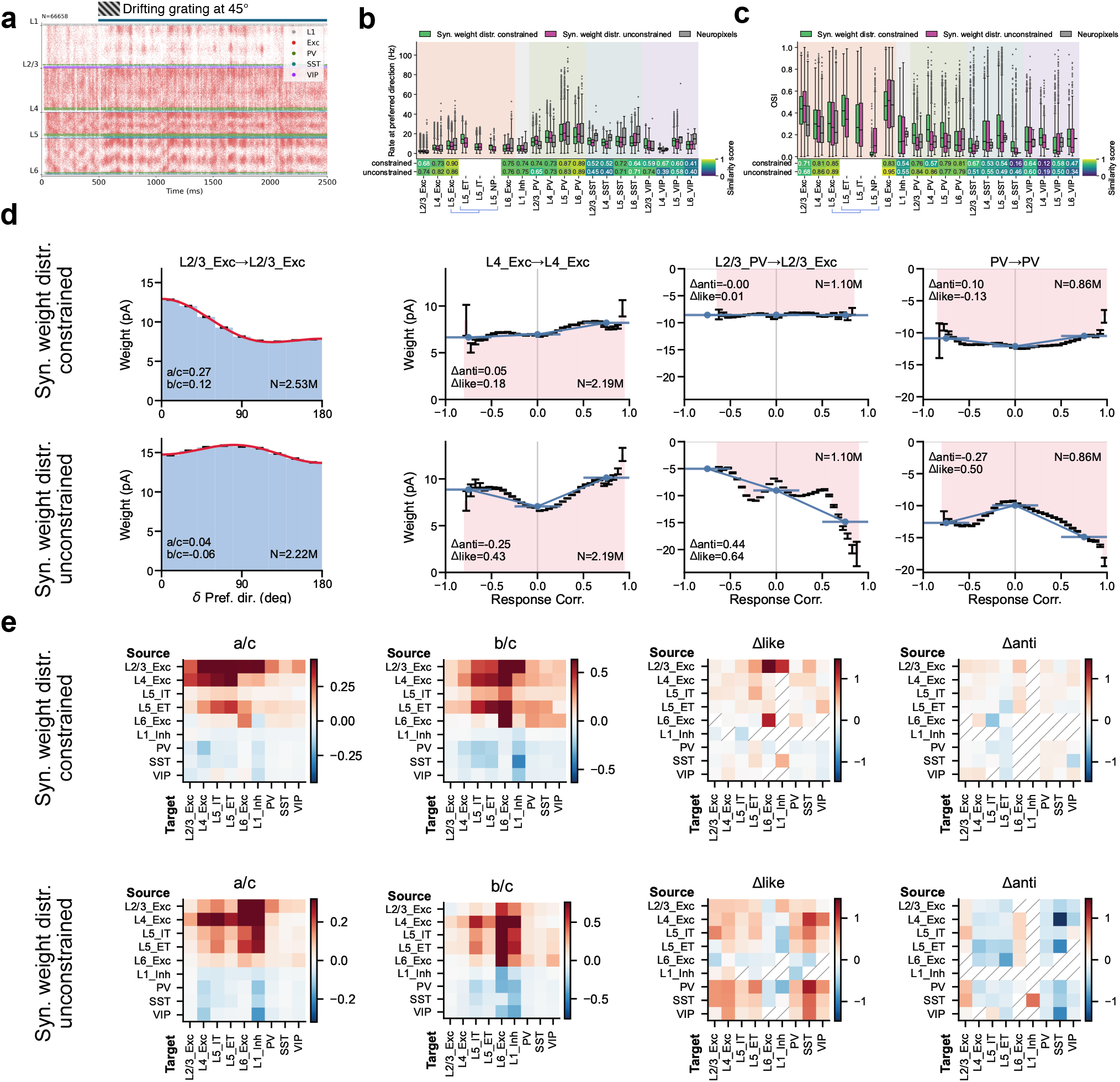
Effect of biological weight constraints. Analyses shown aggregate across *N* = 10 individually trained models per condition. **a**, Raster plot of a network trained without constraints on the synaptic weight distributions. **b, c**, Firing rates to preferred direction and OSI for networks trained with and without synaptic weight distributions constrained to the experimental data. **d**, Examples of synaptic weight vs. preferred-direction difference or response correlation for networks trained with and without synaptic weight distribution constraints. **e**, Heatmaps summarizing effect sizes and correlation-derived Δ metrics across cell-type pairs for networks trained with and without synaptic weight distribution constraints.

Remarkably, removing weight-distribution constraints altered the detailed structure of the emergent synaptic rules despite matching activity statistics (Fig. 6d,e). For example, the L2/3 Exc → L2/3 Exc pathway shifted from direction like-to-like to orientation anti-like-to-like, and the relative magnitudes of like-to-like effects were substantially redistributed across cell-type pairs. For response correlations, the general trend of like-to-like excitatory and anti-like-to-like inhibitory connections was disrupted: in unconstrained models, both strongly correlated and anti-correlated neuron pairs acquired strong weights (Fig. 6e).

These results emphasize that biological constraints can be valuable even when they are not necessary to fit functional response statistics. By limiting the space of allowable parameter distributions, constraints can guide training toward solutions that better approximate real brain circuit mechanisms. This suggests a practical strategy for future model development: use functional objectives to ensure that neural activity is realistic, incorporate the available biological constraints to facilitate realistic circuit mechanisms, and use targeted constraint ablations to identify which constraints support specific wiring motifs and cell-type-level organization.

### Discussion

This work presents models of mouse V1 that integrate multimodal data of unprecedented breadth— EM connectomics and multipatch synaptic physiology^19,22^, intrinsic electrophysiology of 19 cell types^13,14^, and extensive *in vivo* Neuropixels recordings^21^—together with a differentiable, GPU-accelerated training framework enabling end-to-end optimization of ~67,000-neuron circuits on a single GPU in ~6.5 h. Models trained on a limited set of drifting grating responses generalize well to diverse stimuli, providing cell-type-resolved predictions about synaptic organization, inhibitory function, and the role of biological priors.

While full-brain connectomic reconstructions have been achieved in *C. elegans*^83,84^ and *Drosophila*^85,86^, electrophysiological characterization of cell types and synaptic connections in these species is currently less well resolved than in the mouse cortex, and *in vivo* recordings rely mostly on slower optical imaging rather than high temporal-resolution electrophysiology^87^. The broad range of data for mouse V1 integrated here constitutes some of the most complete characterizations of a brain circuit available^7^.

These technical capabilities translate into a set of scientific findings about cortical circuit organization. The generalization noted above is remarkably broad: from a narrow training set, the models reproduce cell-type-specific firing rates, selectivity, and gain modulation across 2-s-long DGs at multiple contrasts, 118 Brain Observatory natural scenes, and thousands of ImageNet images. Non-trivial phenomena that were not part of the training objective, such as the contrast-dependent dichotomy between SST and VIP populations^69^, emerge naturally. This suggests that biological architecture, combined with cell-type-resolved activity constraints, captures key principles of cortical processing rather than fitting stimulus-specific patterns—complementing earlier large-scale models that relied on forward simulation for evaluation^23,42,88^.

Analysis of the trained synaptic weights revealed functional wiring rules that extended well beyond a uniform like-to-like motif. We found that excitatory connections generally follow like-to-like organization^32–34^, but inhibitory pathways exhibit a far more diverse repertoire—anti-like-to-like and non-monotonic relationships that vary by cell type and layer^73,74^. These predictions span the full set of cell types we used, including deep-layer populations (e.g., L6) not yet probed experimentally. Where comparison with connectomic data was possible, the predictions were confirmed: we found significant like-to-like structure between synaptic weight and preferred-direction similarity for pathways where previous studies, limited by sample sizes, concluded that no such relationship existed^22,33^. The emergence of functional connectomics^22,34^ now provides a route that may soon help evaluate such model-derived predictions directly at synapse resolution across the full set of cell types.

Perhaps the most unexpected finding concerns inhibitory interneurons. Training preferentially reshaped inhibitory—but not excitatory—synaptic structure, giving rise to functionally distinct cohorts defined by outgoing synaptic weight that differed in firing rate, selectivity, and downstream targeting. These differences emerged through training rather than from initialization, indicating that they reflected functional requirements of the circuit. Cohort-specific silencing showed that these small subpopulations—each roughly 1% of the network—exerted outsized network control: silencing high-outgoing-weight inhibitory neurons produced widespread disinhibition and selectivity loss, whereas silencing low-outgoing-weight neurons triggered paradoxical suppression^80,81,89^ via disinhibition of other inhibitory neurons. These results point to an axis of inhibitory heterogeneity, defined by connectivity strength rather than transcriptomic identity^90,91^, that may constitute a fundamental organizing principle for cortical computation.

A persistent challenge in mechanistic circuit modeling is identifiability: multiple parameterizations can satisfy the same activity constraints, limiting interpretability^8–10,92^. Constraint ablations offer a practical handle on this degeneracy. We found that removing biological priors on synaptic weight distributions did not prevent the network from matching aggregate firing-rate and selectivity targets, yet it altered the emergent synaptic rules in cell-type-specific ways. Thus, constraints from synaptic physiology act not merely as regularizers but as selectors among functionally equivalent solutions, guiding the circuit toward configurations that preserve cell-type-level synaptic organization.

More broadly, this work aligns with a growing effort to construct integrative circuit models that are both biologically grounded and quantitatively testable^5,23,44,59,93–97^. As multimodal datasets expand—with richer perturbation protocols, improved interneuron coverage, and synapse-resolution connectomics^22^—differentiable circuit models^28,30^ should enable ever tighter linking of biological priors, learned parameters, and experimentally falsifiable predictions. While large networks can now be simulated at brain scale^98–100^, mechanistic insight requires models whose synapses can be optimized against functional data, making scalable training essential for linking anatomy, dynamics, and computation within a unified framework.

All models, code, and trained parameters from this work are freely shared to support such efforts.

## Acknowledgements

We thank the founder of the Allen Institute, Paul G. Allen, for his vision, encouragement, and support.

## Data availability

All experimental datasets used in this study are publicly available. The Allen Cell Types Database, Allen Institute Synaptic Physiology dataset, and Allen Visual Coding Neuropixels dataset are available through the Allen Brain Map portal (https://portal.brain-map.org). The MICrONS dataset^22^ is available at https://www.microns-explorer.org. The V1DD dataset^31^ is available at https://github.com/AllenInstitute/v1dd_physiology. Trained model weights, simulation outputs, and network configuration files generated in this study are available at https://github.com/AllenInstitute/biorealistic-v1-model.

## Code availability

All custom code used in this study is available at https://github.com/AllenInstitute/biorealistic-v1-model. The repository includes instructions for installing dependencies and reproducing the results reported in this manuscript.

## Author contributions

S.I. contributed to Conceptualization, Data curation, Formal analysis, Methodology, Software, Writing—Original Draft, and Writing—Review & Editing. D.H. contributed to Conceptualization, Data curation, Formal analysis, Methodology, Software, Writing—Original Draft, and Writing— Review & Editing. J.G.F. contributed to Conceptualization, Data curation, Formal analysis, Methodology, Software, Writing—Original Draft, and Writing—Review & Editing. K.D. contributed to Data curation, Methodology, Software, and Writing—Review & Editing. J.A. contributed to Data curation, Formal analysis, Methodology, Software, Writing—Original Draft, and Writing—Review & Editing. G.C. contributed to Conceptualization, Methodology, Software, and Writing—Review & Editing. C.M. contributed to Conceptualization, Funding acquisition, Methodology, Software, Supervision, and Writing—Review & Editing. W.M. contributed to Conceptualization, Funding acquisition, Methodology, Software, Supervision, and Writing—Review & Editing. A.A. contributed to Conceptualization, Data curation, Funding acquisition, Methodology, Software, Supervision, Writing—Original Draft, and Writing—Review & Editing. D.H. and J.G.F. contributed equally. A.A. is the corresponding author.

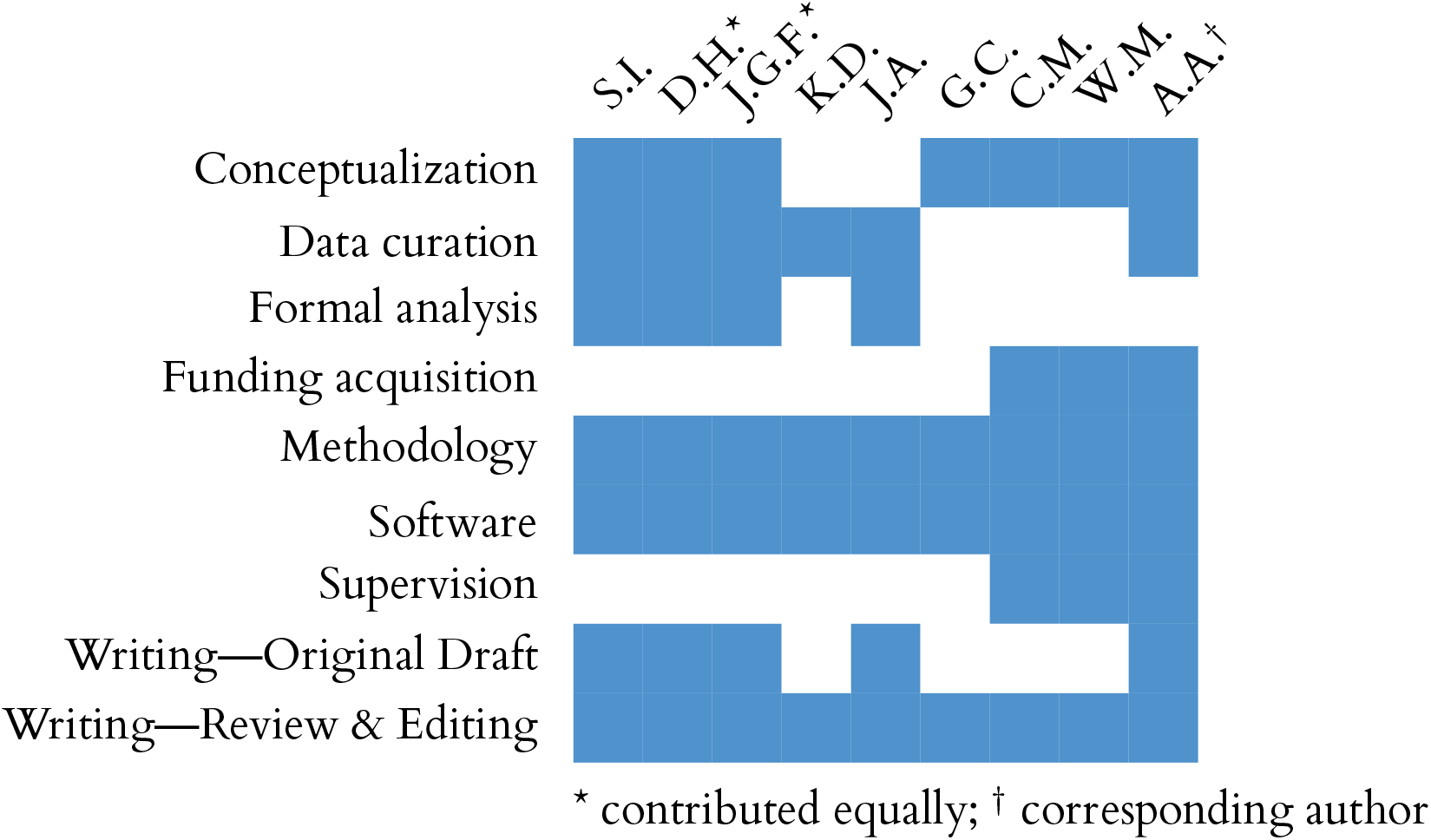

## Methods

### V1 network construction

#### Data sources

The V1 model integrates data from several Allen Institute datasets. Synaptic physiology data—paired whole-cell recordings providing PSP/PSC amplitudes and kinetics across cell-type combinations—were obtained from the Allen Institute Synaptic Physiology dataset^19^. Cell types were identified using Cre-driver lines targeting PV, SST, and VIP interneurons as well as excitatory populations, following the earlier modeling approach^23^. Electron microscopy connectomics data were drawn from the MICrONS dataset^22^, an EM volume spanning multiple visual cortical areas with matched 2-photon calcium imaging measurements of neural activity from the same neurons, and from the V1 Deep Dive (V1DD) dataset^31^, which provides similar calcium imaging and EM data centered in V1. Due to its sole focus on V1, the V1DD dataset was used to derive connection probabilities and spatial spread parameters. Interneurons in V1DD were identified by innervation pattern—proximal-targeting, distal-targeting, and inhibitory-cell-targeting—and assigned to model PV, SST, and VIP types, respectively; interneurons with somata in L1 were assigned to the L1 Inh type. MICrONS data were used to characterize LGN axon synapse statistics, which are not yet available in V1DD, as well as in-degree distributions by cell class and per-connection synapse count distributions. For the latter two, we used MICrONS because it currently has more extensive proof-reading than V1DD, allowing better statistical estimation—under the assumption that in-degree and synapse count distributions do not differ substantially between V1 and nearby higher visual areas (whereas connection probabilities within these different areas may be reasonably expected to differ). Intrinsic electrophysiology and cell-type characterization data from the Allen Cell Types Database^13,14^ provided the GLIF neuron models, and *in vivo* recordings from the Neuropixels Visual Coding^21^ dataset provided firing-rate and selectivity targets for model evaluation and training (see Multi-objective training loss).

#### Network composition

The network contains 19 cell types organized by cortical layer and transcriptomic subclass, with the limitation that little data are available to differentiate certain subclasses in terms of their connectivity and other properties. This resulted in using six excitatory populations (L2/3 excitatory, L4 excitatory, L5 ET excitatory, L5 IT excitatory, L5 NP excitatory, and L6 excitatory) and thirteen inhibitory populations (PV, SST, and VIP in each of L2/3, L4, L5, and L6, plus L1 Inh). Neurons are represented by GLIF Type-3 point models (GLIF_3_; LIF with after-spike currents) fit to individual mouse V1 neurons from the Allen Cell Types Database^13^ and parameterized by experimentally measured membrane capacitance, input resistance, resting potential, spike threshold, after-spike current amplitudes and time constants, and refractory period. Only models achieving an explained variance ratio ≥ 0.7 on a held-out fixed-noise stimulus were retained, resulting in 201 distinct cell models that we used to construct the V1 networks. Multiple GLIF_3_ models per cell type capture within-type electrophysiological diversity, and, consequently, most cell types in our V1 models are represented by multiple GLIF_3_ models.

The densities of the excitatory and inhibitory neurons were estimated for each layer from the MICrONS EM data. Within inhibitory neurons, relative populations of PV, SST, and VIP neurons were set for each layer based on the data from Lee et al.^101^. For a cylindrical cortical column with 400 µm radius, the total number of neurons is ~67, 000. At construction, each V1 neuron was assigned a random position within the cylindrical column, with depth drawn from layer-appropriate ranges. Each neuron was also assigned a tuning angle, which was used to guide both network construction (e.g., LGN inputs, orientation-dependent connectivity rules) and model-training objectives. Finally, each neuron was assigned a target in-degree 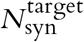, representing its total dendritic input synapse count, drawn from a cell-type-specific log-normal distribution. These distributions were fit by maximum likelihood to total input synapse counts quantified for each cell class in the MICrONS volume^22^, using only neurons with proofread dendritic reconstructions and excluding those with fewer than 700 synapses as likely incompletely reconstructed. Cortical layer assignments were determined from soma positions^22^. The target in-degree serves as a proxy for dendritic extent and is used during both connectivity assignment (Target-size modulation) and synaptic weight construction (Synaptic weight distributions).

#### V1 recurrent connectivity

The baseline connectivity between V1 cell types is modeled as a distance-dependent Gaussian: for each pre–post cell-type pair *a* → *b*, the connection probability decays as a Gaussian function of lateral distance with peak probability 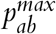 (at zero separation) and spatial spread σ_*ab*_. These two parameters were fit for each cell-type pair by maximum likelihood estimation (MLE) applied to connection probability data from the V1DD dataset^31^, supplemented by synaptic physiology measurements^19^ and literature values where V1DD data were insufficient. This distance-dependent Gaussian profile serves as the foundation for recurrent connectivity, which is then refined by three additional rules: an orientation-dependent spatial modulation, a like-to-like modulation of excitatory connections, and a target-size modulation based on EM-derived in-degree distributions. Axonal transmission delays for recurrent connections vary across cell-type pairs, ranging from 0.88 to 2.92 ms, and were assigned based on the presynaptic–postsynaptic cell-type combination.

#### Spatial connectivity rule

The baseline Gaussian connectivity profile is extended to incorporate orientation-dependent spatial structure^23,36^: excitatory presynaptic ensembles are elongated—in terms of the arrangement of the receptive fields of the presynaptic neurons—along the postsynaptic neuron’s preferred orientation, while inhibitory ensembles are displaced ahead of the preferred direction. We implement this as an anisotropic, orientation-dependent bivariate normal connectivity profile:

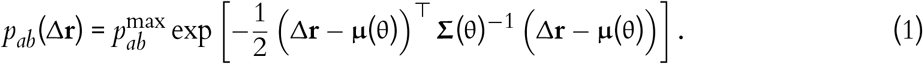

where Δ**r** = **r**_pre_ − **r**_post_ is the lateral displacement vector (excluding cortical depth), 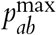 is the peak connection probability, θ is the tuning angle of the target neuron, and **µ**(θ) and **Σ**(θ) are the mean and covariance of the Gaussian profile, respectively. Note that Δ**r** and **µ**(θ) are 2D vectors, and **Σ**(θ) is a 2 × 2 matrix. For **inhibitory** presynaptic neurons, the centroid is displaced by *d*_*R*_ = 50 µm along the postsynaptic neuron’s preferred orientation, with isotropic spread: **µ**_*I*_ = *d*_*R*_[cos θ, sin θ]^*T*^, 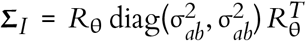, where *R*_θ_ is a rotation matrix. For **excitatory** presynaptic neurons, the centroid is displaced by *d*_*R*_ in the opposite direction, with anisotropic covariance (κ = 1.5): **µ**_*E*_ = −*d*_*R*_[cos θ, sin θ]^*T*^, 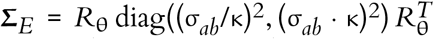. The displacement is applied only within 1.5 times the core radius (default 200 µm) from the network center in order to prevent E/I connectivity imbalances in edge neurons; outside this region, it is set to zero, and the profile reduces to a centered Gaussian. Throughout, the *core* refers to this central 200 µm-radius cylinder and the *periphery* to the surrounding region extending to the full network boundary (400 µm radius); the core provides the high-fidelity functional readout, and the periphery provides contextual support to reduce boundary artifacts.

#### Orientation-dependent (like-to-like) modulation

Experimental evidence shows that excitatory neurons with similar orientation preferences are connected at higher rates than expected from spatial proximity alone—a phenomenon termed like-to-like connectivity^32,71^. For excitatory-to-excitatory connection types, connection probability is additionally modulated by the orientation difference between pre- and postsynaptic neurons:

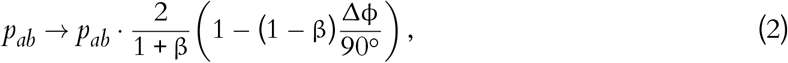

where Δϕ = |ϕ_pre_ − ϕ_post_| (folded to 0–90°) and β controls the strength of like-to-like bias. When β < 1, the rule elevates probability for similarly tuned neurons and reduces it for orthogonally tuned neurons. The modulation averages to unity over all orientation differences, preserving 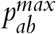.

#### Target-size modulation

As described above (Network composition), each V1 neuron is assigned a target in-degree 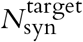 representing its total dendritic input synapse count. Connection probability is scaled by the ratio of the neuron’s target in-degree to its population mean:

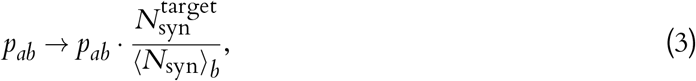

so that neurons with larger dendritic arbors receive proportionally more connections, preserving the log-normal variability in total input observed in EM data. The same size parameter is reused during synaptic weight assignment (see Synaptic weight distributions).

#### Synaptic kinetics

Synaptic currents are modeled with a double-alpha function,

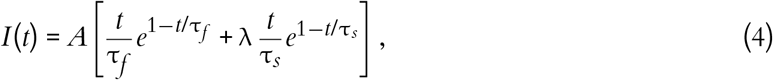

where *A* is the overall amplitude, τ_*f*_ and τ_*s*_ are the fast and slow decay time constants, and λ is the relative amplitude of the slow component, to capture distinct fast and slow components observed in voltage-clamp recordings from the Synaptic Physiology dataset^19^. For each synapse, averaged voltage-clamp response traces were sign-corrected by synapse type (excitatory or inhibitory), aligned to response onset, and baseline-subtracted. Fits were done using nonlinear least squares; the double-alpha function substantially improved fits compared to a single-alpha function across connection types (e.g., *R*^2^ increased from 0.79 to 0.92 for PV to L5 IT; Fig. 1e). Fits were retained only if *R*^2^ > 0.3, the fast time constant 0.2 ms < τ_*f*_ < 15 ms, the slow time constant τ_*f*_ < τ_*s*_ < 15 ms, and both amplitude components were positive. For each pre–post cell-type pair with ≥ 5 quality-filtered recordings, the median τ_*f*_, τ_*s*_, and amplitude ratio λ = *A*_*s*_/*A*_*f*_ were computed. When fewer than 5 recordings were available for a specific pair, broader groupings were substituted: all excitatory types were pooled as a single pyramidal class, and all inhibitory types as a single interneuron class. Because synaptic physiology recordings were not available for L5 NP neurons, synaptic kinetic parameters for this population were approximated using L6 excitatory values, motivated by transcriptomic evidence that L5 NP neurons are more closely related to L6 corticothalamic neurons than to L5 ET or IT populations^12,102^.

#### PSP amplitude distributions

Synaptic strengths were drawn from log-normal distributions^40,41^ fitted to experimental PSP amplitudes. For each connected pair in the Synaptic Physiology dataset^19^, the 90th-percentile pulse response amplitude was used as a robust estimate of synaptic strength under elevated release probability. Log-normal distributions were fit by maximum likelihood for each connection type with *N* ≥ 5 pairs.

Because individual GLIF_3_ models differ in intrinsic properties, a per-model calibration step was used to convert desired PSP amplitudes (in mV) to the simulator’s dimensionless weight scale. For each of the 201 GLIF_3_ models and each connection type, simulations were carried out where a single presynaptic spike was delivered at unit weight using the calibrated double-alpha kernel, and the resulting peak membrane potential deflection was recorded as the unitary PSP amplitude. The final simulator weight was then obtained by dividing the desired PSP amplitude by this unitary value.

#### Synaptic weight distributions

Because neurons differ in their total number of input connections (in-degree), per-connection synaptic weights must be adjusted so that neurons with more inputs receive individually weaker connections, preserving the experimentally observed distribution of PSP amplitudes across the population^39^. This is achieved by exploiting the closure of the log-normal family of distributions under quotients: dividing one log-normal variable by an independent log-normal variable yields a log-normal result whose log-variance equals the sum of the two individual log-variances. Because the final simulator weight is obtained by dividing the sampled weight by the target in-degree, up to the population-mean normalization factor introduced below, the sampled distribution for *W* must be chosen so that the quotient 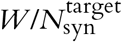 reproduces the experimentally fitted PSP distribution.

Here, LogNormal(µ, σ) denotes a log-normal distribution such that ln *X* ~ 𝒩 (µ, σ^2^); equivalently, the median is exp(µ). For each connection type *a* → *b*, the experimentally fitted PSP distribution LogNormal(µ_PSP_, σ_PSP_) and the target in-degree distribution LogNormal(µ_*N*_, σ_*N*_) together determine a per-connection weight distribution LogNormal(µ_*W*_, σ_*W*_) with

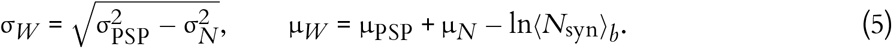

A weight *W* is drawn independently for each connection, and the final weight is scaled by the ratio of the population-mean in-degree to the target neuron’s in-degree:

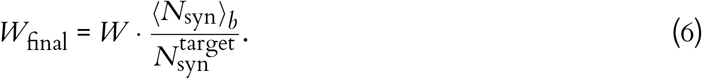

Because *W* and 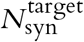 are independent by construction, the quotient 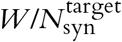 is log-normal with variance 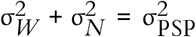 and median exp(µ_PSP_), exactly recovering the experimental PSP distribution. When σ_PSP_ < σ_*N*_, σ_*W*_ is set to a small positive value, effectively removing weight variability for that connection type.

#### LGN inputs

Visual input is provided through an LGN model comprising 17,400 units, implemented with Brain Modeling ToolKit (BMTK)’s FilterNet module^37^, which generates spike trains by passing stimuli through spatiotemporally separable linear–nonlinear filters approximating LGN receptive field properties. The LGN visual field subtends 120 × 80 degrees of visual angle (azimuth × elevation), and input images are 120 × 80 pixels with a resolution of one pixel per degree. The LGN model comprises sustained-ON, sustained-OFF, and transient-OFF functional classes of LGN cells^23^, characterized by Durand et al.^38^. Direction selectivity in V1 arises from convergent input of spatially offset sustained and transient afferents, following the mechanism of Lien & Scanziani^103^, as described in previous studies^23,104^.

Thalamocortical connection statistics were constrained using anatomically identified LGN axons from the MICrONS volume^22^. Forty-three LGN axons were reconstructed and their synapses onto V1 neurons quantified (Fig. 1g). Based on the number of synapses from LGN axons to each cell type, the relative probability each V1 neuron received from LGN was calculated (relative to L4 Exc cells; Extended Data Fig. 1). Per-connection synapse counts were modeled with the Yule–Simon distribution^105^,

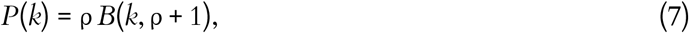

where ρ > 0 is the shape parameter and *B*(·, ·) the beta function. The Yule–Simon distribution arises from preferential attachment^106^ and produces a power-law tail with exponent ρ+ 1 without requiring a minimum-degree threshold. The shape parameter was fit by maximum likelihood to the MICrONS data for each of 6 excitatory types (layer segregated) and 3 inhibitory types (layer aggregated).

For each V1 target neuron, the total LGN synapse count was estimated from the neuron’s indegree (with LGN synapses comprising a fixed fraction of the total, e.g., 20% for L4 excitatory neurons; Extended Data Fig. 1a), and the number of distinct connections was derived as *N*_conn_ = *N*_syn_ ·(ρ−1)/ρ. Source LGN neurons were selected probabilistically based on receptive-field proximity and functional subtype following the procedure outlined in previous work^23^, with more realistic connection probability derived from the MICrONS data. Connections with higher synapse counts were preferentially assigned to spatially closer sources (Gaussian kernel with 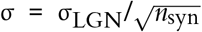, σ_LGN_ = 5°), so that stronger connections arose from more precisely aligned receptive fields. All LGN-to-V1 connections use a fixed axonal delay of 1.7 ms.

#### Background inputs

In addition to LGN-driven visual input, each V1 neuron receives excitatory background drive representing possible inputs from the rest of the brain. The background population consists of 100 virtual Poisson spike-generator units with a frequency of 250 Hz. Each V1 neuron is connected to 4 randomly selected background units. Background synaptic weights were determined per cell model by adjusting them—in isolation from recurrent connections—until each cell model’s spontaneous firing rate matched its biological target, given spontaneous LGN activity held fixed throughout this process. All background connections use a delay of 1.0 ms.

### Implementation of the TensorFlow-based simulator

While the model can be run on the NEST simulator using CPUs, gradient-based fitting requires a differentiable implementation that supports backpropagation through time (BPTT). We therefore implemented a TensorFlow-based simulator following the general approach of Chen et al.^28^, and substantially optimized it for the present study to improve GPU memory efficiency and throughput.

Single-neuron dynamics (GLIF_3_). All neurons were modeled with the GLIF_3_ formalism^13^, which extends the leaky integrate-and-fire model with after-spike currents (ASCs) that capture refractory/adaptive effects from ionic channel dynamics. Simulations used forward Euler integration with time step Δ*t* = 1 ms.

For neuron *j*, let *V*_*j*_[*n*] denote membrane potential at discrete step *n*, and let 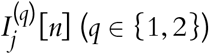 denote the two ASC states. Defining

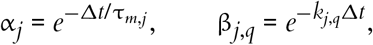

where τ_*m,j*_ is the membrane time constant and *k*_*j,q*_ the decay rates of the ASCs, the dynamics are

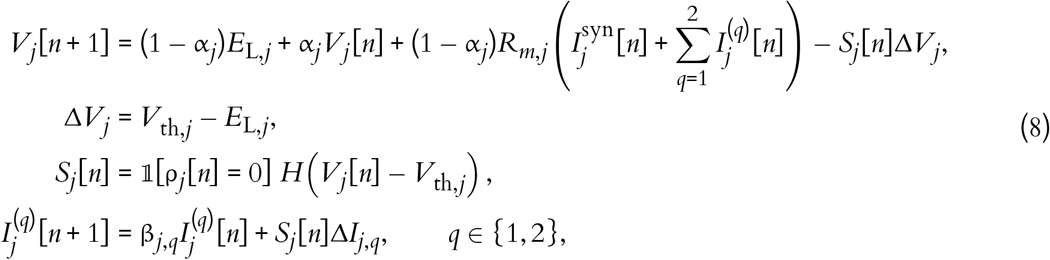

where *R*_*m,j*_ is membrane resistance, 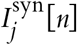 is the total synaptic current (recurrent V1 + LGN + BKG), *E*_*L,j*_ the resting membrane potential, *V*_th,*j*_ the firing threshold, *S*_*j*_[*n*] ∈ {0, 1} is the spike indicator, *H* is the Heaviside function, and ρ_*j*_[*n*] is an absolute-refractory countdown state (in ms steps). The term *S*_*j*_[*n*]Δ*I*_*j,q*_ implements the spike-triggered increment of the *q*-th ASC. We used a *soft reset* (subtracting Δ*V*_*j*_ upon spiking) rather than a *hard reset*, following prior surrogate-gradient RSNN work and Chen et al.^28,107^. This preserves forward spiking behavior while improving gradient propagation.

The refractory countdown is updated as

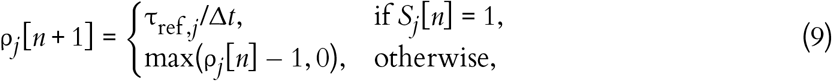

so that spike emission is disabled during the cell-model-specific absolute refractory period τ_ref,*j*_ (2–8 ms in the fitted GLIF_3_ parameter set^13^).

Neuron electrophysiological parameters were taken from the fitted GLIF_3_ models in the Allen Cell Types Database^13^ (201 selected neurons).

#### Double-alpha synaptic current model

Synaptic interactions were modeled with a current-based double-alpha formulation, capturing both fast and slow PSC components. Each synapse belongs to a receptor family *r* — a distinct synaptic input channel defined by a unique set of kinetic parameters (τ_*r,f*_, τ_*r,s*_, λ_*r*_) fit from paired-recording physiology data^19^; different pre- and postsynaptic cell-type combinations give rise to distinct receptor families. For clarity, we write the dynamics for one postsynaptic neuron *j* and one receptor family *r* (the total current is the sum over families and presynaptic sources). The receptor-family contribution is

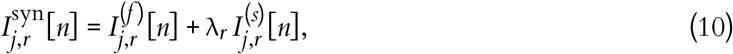

where λ_*r*_ is the receptor-family-specific ratio of slow to fast component amplitude (corresponding to *A*_*s*_/*A*_*f*_ in the continuous parametrization), and the full synaptic current is

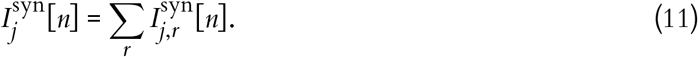

Rather than explicitly convolving each presynaptic spike train with a kernel (which requires storing spike histories), we use an equivalent linear-state implementation^108,109^. Let 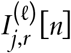 and 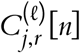 denote the two state variables of alpha component *𝓁* ∈ {*f, s*} (fast/slow), with decay time constants τ_*r,𝓁*_. For each component *𝓁*,

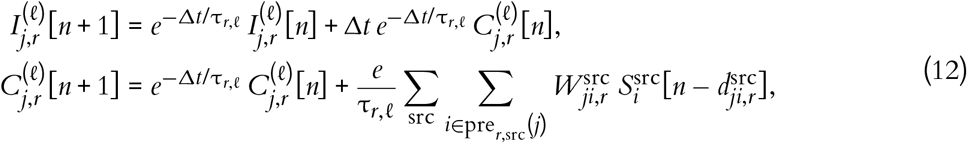

where 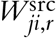 is the signed synaptic weight, 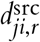 is the axonal delay (in simulation steps), and pre_*r*,src_(*j*) is the presynaptic set for neuron *j*, receptor family *r*, and source src ∈ {rec, lgn, bkg}. Because the simulation uses a discrete time step of Δ*t* = 1 ms, all axonal delays are rounded to the nearest integer millisecond; sub-millisecond precision in the biologically specified delay values (e.g., the 0.88–2.92 ms range for recurrent connections) is therefore approximated to 1 ms resolution. This formulation yields constant-cost updates per time step and is well-suited to parallel implementation. The model contains 19 cell types; because inhibitory neurons were aggregated across cortical layers in the physiology fitting due to limited experimental statistics, this yields 11 effective cell types (including general excitatory type used for the LGN and background connections) and therefore 11 × 11 = 121 distinct directed presynaptic–postsynaptic cell-type pair families.

The current implementation did not include synaptic plasticity or creation/elimination of connections due to a substantially higher computational expense associated with such functionality and a lack of extensive data to parameterize diverse connection types. This choice is appropriate for simulations of short-timescale (e.g., seconds) network dynamics studied here, while leaving the framework compatible with future extensions that may include plasticity mechanisms^104,110^.

#### State normalization and numerical stability

To improve numerical stability in large-scale mixed-precision simulation^111^, we normalize voltages and synaptic weights using the neuron-specific threshold distance Δ*V*_*j*_ = *V*_th,*j*_ − *E*_L,*j*_. We define

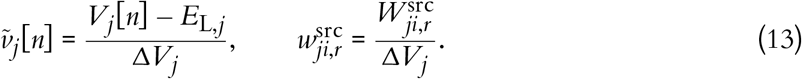

Under this transformation, reset is at 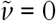 and threshold is at 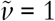.

Using normalized currents (same scaling by Δ*V*_*j*_) and suppressing receptor/source indices for readability, the membrane update becomes

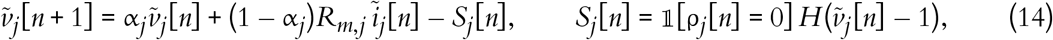

with 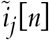 the normalized total current (ASC + synaptic contributions). The corresponding ASC and PSC state updates retain the same linear form as Eqs. (8) and (12), with amplitudes/weights expressed in normalized units. This scaling keeps principal state variables in comparable numerical ranges while preserving the underlying biophysical dynamics.

### Gradient-based learning and biologically constrained optimization in RSNNs

Training the recurrent V1 circuit requires solving temporal credit assignment under non-differentiable spike generation while preserving structural synaptic constraints (e.g., fixed sign by cell class). In the Results (Fig. 2a,b), this corresponds to the differentiable fitting stage that maps a biologically grounded initialization to a constrained physiological operating regime.

#### BPTT factorization

Backpropagation through time (BPTT)^112^ unfolds the recurrent network across time and applies gradient descent on the resulting acyclic computational graph. Let *n* = 1, …, *T* index simulation steps within a training chunk (here *T* = 500 for a 500 ms chunk with Δ*t* = 1 ms). We denote by **h**_*n*_ the full simulator hidden state at time step *n*, which comprises all dynamical variables of the network, including membrane potentials, ASC states, synaptic states, refractory counters, and delayed-spike buffer states. Inputs **x**_*n*_ include stimulus-dependent LGN drive and background spikes. The vector **z**_*n*_ ∈ {0, 1}^*N*^ denotes the binary spike output of the network at step *n*, where (**z**_*n*_)_*j*_ = 1 indicates that neuron *j* emits a spike and 0 otherwise. We use bold notation for vectors and matrices throughout. The recurrent dynamics can be written as

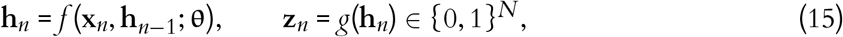

where **h**_0_ denotes the initial network state and **θ** the collection of trainable synaptic parameters (recurrent and background weights in normalized coordinates), and *g* implements spike generation applied to the hidden state.

For a scalar loss ℒ, the exact BPTT gradient is

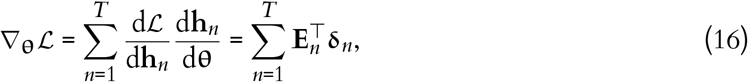

with backward learning signals **δ**_*n*_ and forward eligibility traces **E**_*n*_^109,112,113^:

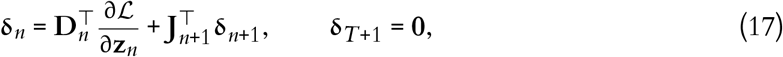

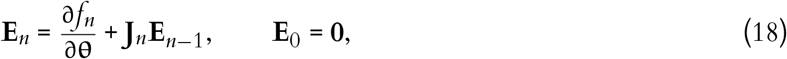

where **J**_*n*_ = ∂**h**_*n*_/∂**h**_*n*−1_ and **D**_*n*_ = ∂**z**_*n*_/∂**h**_*n*_. In practice, these derivatives are computed automatically by TensorFlow^114^.

#### Surrogate gradient for spike generation

Because *g* contains a Heaviside threshold, ∂**z**_*n*_/∂**h**_*n*_ is zero almost everywhere under the exact derivative. We therefore use BPTT with surrogate gradients^50,51,115^, preserving exact spiking in the forward pass while replacing the threshold derivative during backpropagation.

Let 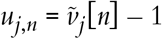 denote normalized distance to threshold. We use a triangular surrogate

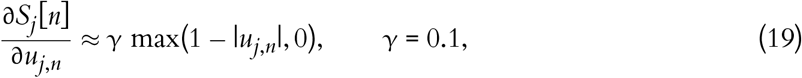

which has support in a unit-width window around the threshold and a damped slope γ^47^. In our setting, γ was chosen to stabilize long-horizon BPTT in the large recurrent network (reducing the risk of gradient explosion) while maintaining useful gradient flow.

#### Biologically constrained optimization

Temporal credit assignment alone does not intrinsically incorporate a biologically consistent geometry of synaptic change. Standard gradient descent optimizers such as Adam^52^ operate in a Euclidean geometry with additive parameter updates. Although computationally effective^28,47,48^, they are biologically agnostic: they do not intrinsically preserve the synaptic sign (which requires explicit constraints to enforce Dale’s law) and do not respect the approximately lognormal distribution of cortical synaptic strengths^41,55,56^.

In contrast, exponentiated gradient descent (EGD), a mirror-descent method that performs additive updates in log-space, is intrinsically sign-preserving and is more compatible with the heavy-tailed, lognormal-like statistics observed in cortex^53–56^. Indeed, EGD not only retains the standard gradient descent optimization power while embedding biologically meaningful constraints, but also excels at biologically relevant tasks with sparse signals and confers robustness to synaptic pruning^56^.

For each trainable synapse *k*, we decompose the normalized weight *w*_*k*_ into a fixed sign mask *s*_*k*_ and a positive magnitude *a*_*k*_:

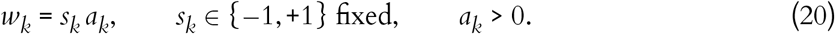

The sign *s*_*k*_ is determined by presynaptic cell class (consistent with Dale’s law) and remains fixed during training. Optimization is performed on the positive magnitudes **a** (equivalently, in log-magnitude space). To make the multiplicative update unambiguous, we write it componentwise as

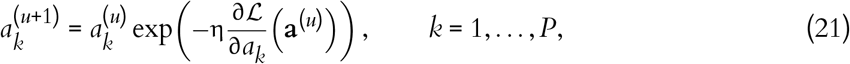

where *P* is the number of trainable synapses, *u* indexes optimizer updates, and η > 0 is the learning rate. This update preserves positivity of every *a*_*k*_ by construction and therefore preserves the synaptic signs when mapped back to *w*_*k*_.

#### Exponentiated Adam optimizer

RSNN training exhibits heterogeneous gradient magnitudes across time, synapse types, and loss terms. To improve optimization stability while retaining multiplicative geometry, we use an *Exponentiated Adam* optimizer: Adam-style adaptive moments^52^ combined with the multiplicative, sign-preserving structure of EGD^55,56^.

For each trainable synapse *k* = 1, …, *P*, let

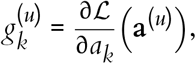

where *u* indexes optimizer updates. Following Adam, we maintain two exponential moving averages of the gradient: a first-moment estimate 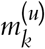, which tracks the mean gradient, and a second-moment estimate 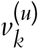, which tracks the mean squared gradient. These quantities are updated componentwise as

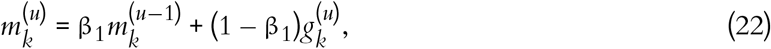

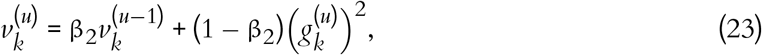

with bias-corrected forms

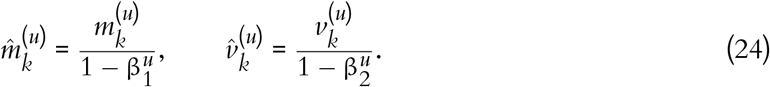

The exponentiated update is then

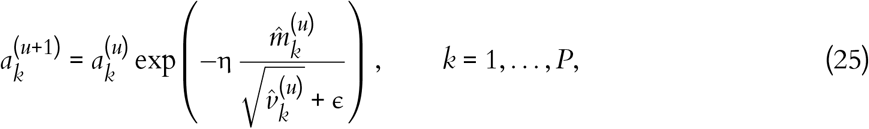

and the signed synaptic weights are recovered as

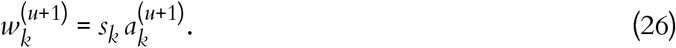

Equivalently, in vector form,

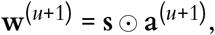

where ⊙ denotes elementwise multiplication. This adaptive multiplicative rule preserves positivity of each *a*_*k*_ by construction and therefore preserves the synaptic sign pattern throughout training. Because updates are applied in log-magnitude space, the optimizer is more compatible with multiplicative changes in synaptic strength than standard additive updates and is consistent with the broad, heavy-tailed synaptic weight distributions reported in cortex^54–56,76^. In practice, it provided stable optimization in our large recurrent spiking model while retaining the efficiency of Adam-style adaptive moment estimation. To our knowledge, this is the first application of exponentiated adaptive optimization to a large-scale, biophysically grounded cortical spiking model. Unless stated otherwise, we used η = 5 × 10^−3^, β_1_ = 0.9, β_2_ = 0.999, and ϵ = 10^−11^. A detailed derivation is provided by Galván Fraile^109^.

### Multi-objective training loss

Choosing the training objective is as important as choosing the circuit model itself: if the loss only matches one statistic (for example, firing-rate histograms), the optimizer can satisfy that target through biologically implausible mechanisms, such as excessive population synchrony or unrealistic redistribution of synaptic strength. To avoid such undesirable solutions, the objective must jointly constrain multiple dimensions of activity while remaining numerically well-behaved for gradient-based optimization (positive, smooth, and balanced across terms). The total loss therefore combines multiple activity-matching terms (derived from Neuropixels recordings^21^) with physiological regularizers on membrane voltage and recurrent-weight distributions.

#### Neuropixels target curation

Targets were derived from the Allen Institute’s Visual Coding Neuropixels resource^21^ after stimulus and unit curation, as described below. We restricted training targets to a subset of the “Brain Observatory 1.1” conditions in the Neuropixels dataset: gray-screen spontaneous activity and full-field drifting gratings at 8 directions (0° : 315° : 45°), 80% contrast, 2 Hz temporal frequency, and 0.04 cycles/degree. OSI/DSI targets were derived from the 2 Hz drifting-grating condition.

Cell-type labels in the Neuropixels dataset were aligned to model populations with layer-aware pooling. Excitatory and PV neurons were identified by the waveform (broad spikes: Exc, narrow spikes: PV); their response statistics were taken from only V1. SST and VIP cells were identified in the dataset using optotagging. For SST and VIP, response statistics were pooled across V1 and the other five visual cortical areas (LM, AL, RL, AM, and PM) in the dataset, since V1 counts alone were sparse. L5 excitatory types (ET, IT, and NP) were merged to match Neuropixels granularity. Units with receptive fields larger than 100° in azimuth or elevation were excluded. OSI/DSI target computation included only neurons with preferred-condition firing rate ≥ 0.5 Hz, to avoid spuriously high selectivity estimates from ultra-sparse firing. For firing-rate distributions, an explicit 0 Hz sample was appended per population to preserve the silent-neuron fraction in short windows. The curated target set comprised 3,156 neurons from 48 mice.

#### Training chunks and global loss structure

Each optimizer update uses two simulated chunks of equal duration and batch size:

- spontaneous chunk (gray screen): *T*_ch_ = 500 ms, *B*_spt_ = 5 trials,
- evoked chunk (drifting gratings): *T*_ch_ = 500 ms, *B*_evk_ = 5 trials.

The evoked chunk uses full-field drifting gratings (80% contrast, 0.04 cycles/degree, 2 Hz) with random orientation and phase across trials. The two chunks are simulated within the same training step.

The total objective is

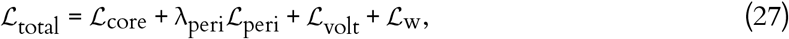

with

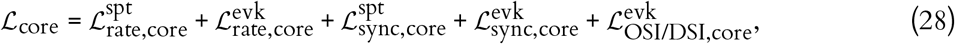

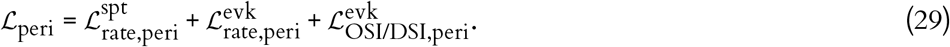

Here, *core* refers to the central analysis region of the model, which provides the primary functional readout, whereas *peri* refers to the surrounding peripheral region, which is included to provide contextual support and reduce edge effects. Accordingly, ℒ_core_ contains the main activity-matching terms evaluated in the core: 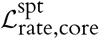 and 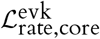 penalize mismatches in spontaneous and evoked firing rate distributions, 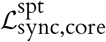 and 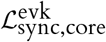 penalize mismatches in spontaneous and evoked synchrony/variability, and 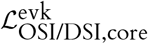 penalizes mismatches in evoked orientation and direction selectivity. The peripheral term ℒ_peri_ contains analogous but down-weighted constraints in the surrounding region: 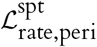 and 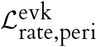 constrain spontaneous and evoked firing rate distributions, and 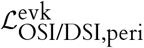 constrains evoked orientation/direction selectivity. Synchrony terms are applied only in the core. The factor λ_peri_ therefore controls how strongly the peripheral contextual region contributes relative to the primary core readout.

Rate and OSI/DSI terms use the full 500-ms window of each chunk. Synchrony is computed on the 200–500 ms window. These loss blocks correspond directly to the schematic in Fig. 2a.

Firing-rate distribution loss. For each chunk ch ∈ {spt, evk}, we match model firing-rate distributions to empirical Neuropixels distributions using a quantile-Huber loss^116^, as in prior RSNN training studies^28,109,117^. Let 𝒞 denote the set of cell types, and let *N*_*c*_ denote the number of model neurons of cell type *c*. For each cell type, we first sample a fixed target rate vector of length *N*_*c*_ from the corresponding empirical Neuropixels distribution by inverse-CDF sampling, sort it, and keep it fixed throughout training:

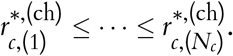

Chunk-specific model firing rates are computed from the 500 ms window:

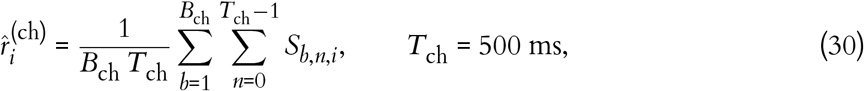

where *B*_ch_ is the number of trials in chunk ch. Within each cell type *c*, model rates are sorted to obtain

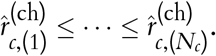

Using *q* = 1, …, *N*_*c*_ to index sorted rank positions within cell type *c*, define the residual 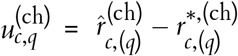. The firing-rate loss is

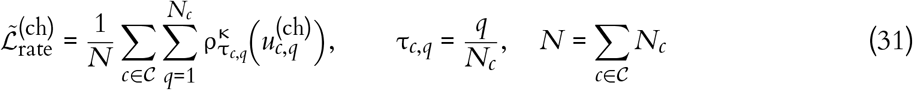

with quantile-Huber penalty

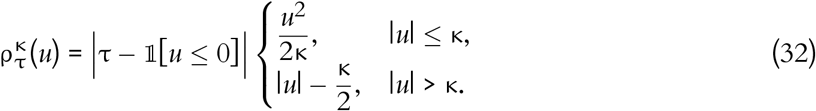

Finally, the chunk-specific firing-rate loss is

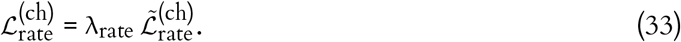

#### Crowd-surrogate OSI/DSI loss

A core difficulty in gradient-training recurrent spiking networks on selectivity targets is that classical single-neuron OSI/DSI definitions require averaging each neuron’s response across many repeated stimulus directions; short 500 ms BPTT chunks with sparse direction sampling therefore yield noisy, non-smooth estimates that destabilize the gradient signal.

We resolve this by developing a *crowd-surrogate OSI/DSI loss* (computed separately for each cell type) that replaces the classical average-over-stimuli with an average-over-neurons within a single trial, yielding a low-variance differentiable estimator of cell-type-level selectivity. The surrogate exploits a structural property of the network: within each cell type, preferred directions are assigned uniformly by construction, so for any fixed stimulus direction, the neurons tile the space of offsets from the preferred direction. Averaging normalized responses across neurons in one trial therefore approximates averaging the population mean normalized tuning curve across stimulus directions. Because this substitution targets the selectivity of the cell-type-averaged aligned tuning curve rather than of an individual neuron, it is well-suited as a cell-type-level training signal but does not replace classical per-neuron OSI/DSI at evaluation time.

Let 𝒞 denote the set of cell types, and let *N*_*c*_ be the number of model neurons of cell type *c*. For evoked trial *b*, let φ_*b*_ denote the presented grating direction, and let 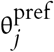 be the assigned preferred direction of neuron *j*. We define the preferred-direction offset

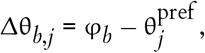

and the trial-averaged evoked firing rate

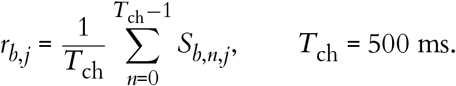

To reduce the influence of cell-to-cell differences in overall firing rate, responses are normalized by an evoked exponential moving average (EMA) with floor *R*_min_ = 0.5 Hz:

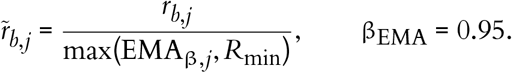

Formally, the population-averaging approximation is

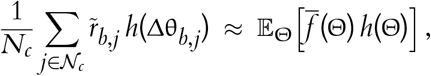

where 𝔼_Θ_[·] denotes expectation over directions Θ within cell type 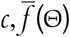 is the cell-type mean aligned normalized tuning curve, and *h* is the angular kernel of interest. In particular, we use *h*(δ) = cos(2δ) for orientation selectivity and *h*(δ) = cos(δ) for direction selectivity. A detailed derivation is provided by Galván Fraile^109^, and an illustration is shown in Extended Data Fig. 12.

Using these definitions, the crowd-surrogate OSI/DSI estimates for trial *b* and cell type *c* are

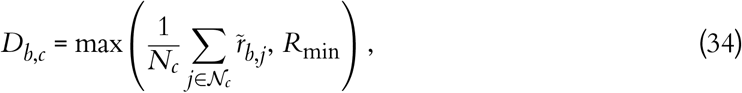

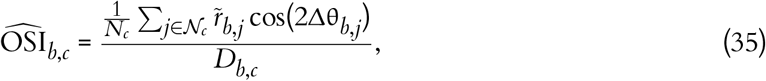

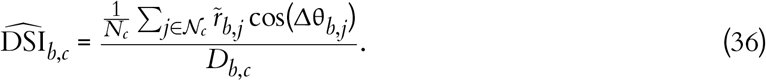

We then average these trial-level quantities over the evoked batch:

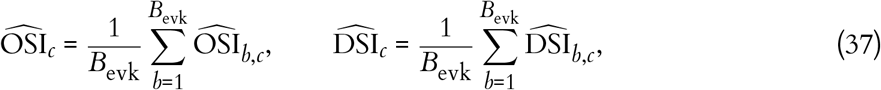

where *B*_evk_ is the number of evoked trials in the batch.

Finally, we match the resulting cell-type surrogates to the Neuropixels targets with a population-weighted mean-squared error (MSE):

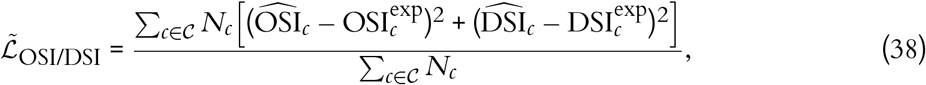

and

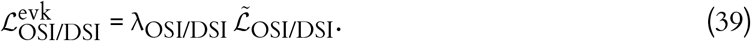

#### Synchronization (multi-scale Fano factor) loss

Firing-rate distributions alone do not constrain shared variability: a network can reproduce correct rates while exhibiting unrealistic population synchronization. We therefore include a synchrony term based on multi-scale Fano factors (variance-to-mean spike-count ratio) computed from excitatory populations in the core. Neuropixels references were precomputed from recordings separated by behavioral state (running versus stationary; we only used the running state for target metrics) and, on average across animals, display the expected cortical signatures across timescales: supra-Poisson variability, and stronger fluctuations in evoked drifting-grating epochs than in spontaneous gray-screen periods (Extended Data Fig. 13)^21^.

Each loss evaluation samples *N*_pool_ random excitatory pools (default *N*_pool_ = 500); pool sizes are drawn from a Gaussian *n*_pool_ ~ 𝒩 (70, 30^2^), clipped with a minimum of 15, to match experimental sample sizes. Spikes pooled across neurons are binned at 14 log-spaced widths 𝒲_bin_ ⊂ [1, 113] ms (drawn from 20 equally spaced bins in log-space spanning 1–1000 ms; widths ≥ half the analysis-window duration are discarded to ensure at least two bins) over the 200–500 ms post-stimulus window. For sampled pool *p* and bin width *w* ∈ 𝒲_bin_,

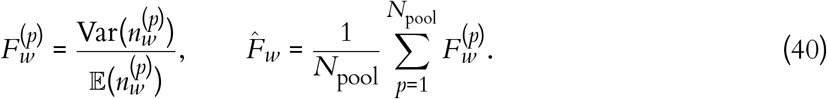

Here 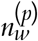 is the total spike count of pool *p* summed over all neurons within a single time bin of width *w*; Var(·) denotes the variance and 𝔼 (·) the expected value (mean), both computed over all non-overlapping bins in the 200–500 ms analysis window. 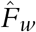 is the pool-averaged Fano factor at bin width *w*. At each training step, excitatory neurons are randomly permuted and pools are drawn as successive non-overlapping slices of that permutation, ensuring each neuron is sampled at most once per pass before the list is re-shuffled. The chunk-specific synchrony loss is

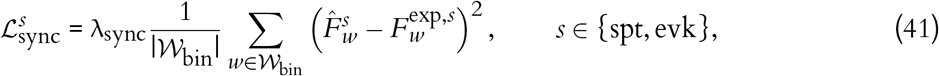

where 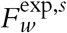 is the experimentally measured Fano factor at bin width *w* for chunk *s*, precomputed from Neuropixels recordings.

#### Voltage regularization

To discourage large voltage excursions during BPTT, we regularize membrane voltage in normalized coordinates (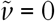 reset, 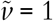 threshold):

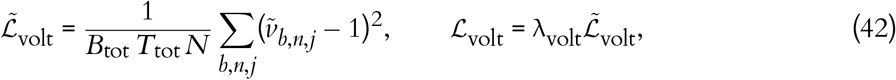

where *B*_tot_ = 10 is the total number of trials across the two chunks in one update and *T*_tot_ is the corresponding number of simulated steps included in this regularizer. This term serves as a weak regularizer on voltage trajectories; functional losses remain responsible for matching activity statistics^28,109,117^.

#### Recurrent-weight regularization (EMD / Wasserstein-1)

A major goal of our work was to train the circuit models while preserving experimentally observed values of synaptic weights. These are not known for individual connections, but the Synaptic Physiology dataset^19^ provides distributions per connection type (defined based on the pre- and postsynaptic cell type). Our implementation of this constraint took advantage of the fact that we initialized network weights at model construction from these experimentally determined distributions. Therefore, we constrain cell-type-wise distributional drift during optimization using Earth Mover’s distance (Wasserstein-1) between current and initial recurrent weights. For recurrent connection class *m* (out of *M* classes), let *n*_*m*_ be the number of synapses and let 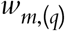 and 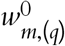 denote the *q*-th smallest current and initial normalized weights, respectively (same sign class):

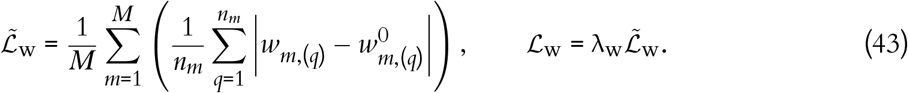

This regularizer penalizes full distributional drift and therefore preserves the experimentally shaped heavy-tailed synaptic landscape while still allowing functionally useful adaptation.

#### Loss coefficients used for reported training runs

Unless stated otherwise, the loss weights were

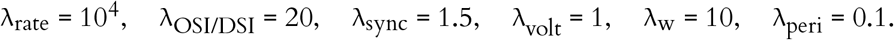

Additional defaults were: quantile-Huber threshold κ = 0.002, evoked-rate EMA decay β_EMA_ = 0.95, and *N*_pool_ = 500 sampled excitatory pools for the synchrony loss.

### TensorFlow implementation for efficient training

The simulator is differentiable by construction, but practical BPTT at the scale of our V1 models is limited by memory traffic and sparse recurrent computation. At each 1 ms step, the simulator updates membrane voltages, ASC, PSC, and delayed-spike states for a large recurrent network while preserving the trajectories needed for gradient propagation. In a naive implementation, this leads to 𝒪 (*T* · *N*) activation storage (where *T* is the number of simulation time steps and *N* the number of neurons) and repeated large recurrent products, which rapidly becomes prohibitive in VRAM and wall-clock time. The implementation therefore combines sparsity-aware kernels, compact state representations, and graph-level optimizations to make the biology-constrained fitting stage in Fig. 2 computationally practical.

#### Exploiting structural and activity sparsity

The recurrent pathway is sparse in two complementary senses: (i) *structural sparsity* in connectivity (about 2.35 × 10^7^ recurrent connections across ~6.7 × 10^4^ neurons, that is, ~0.5% occupancy of all possible directed pairs), and (ii) *activity sparsity* in time

##### Algorithm 1

Event-driven recurrent synaptic current and custom gradient calculation

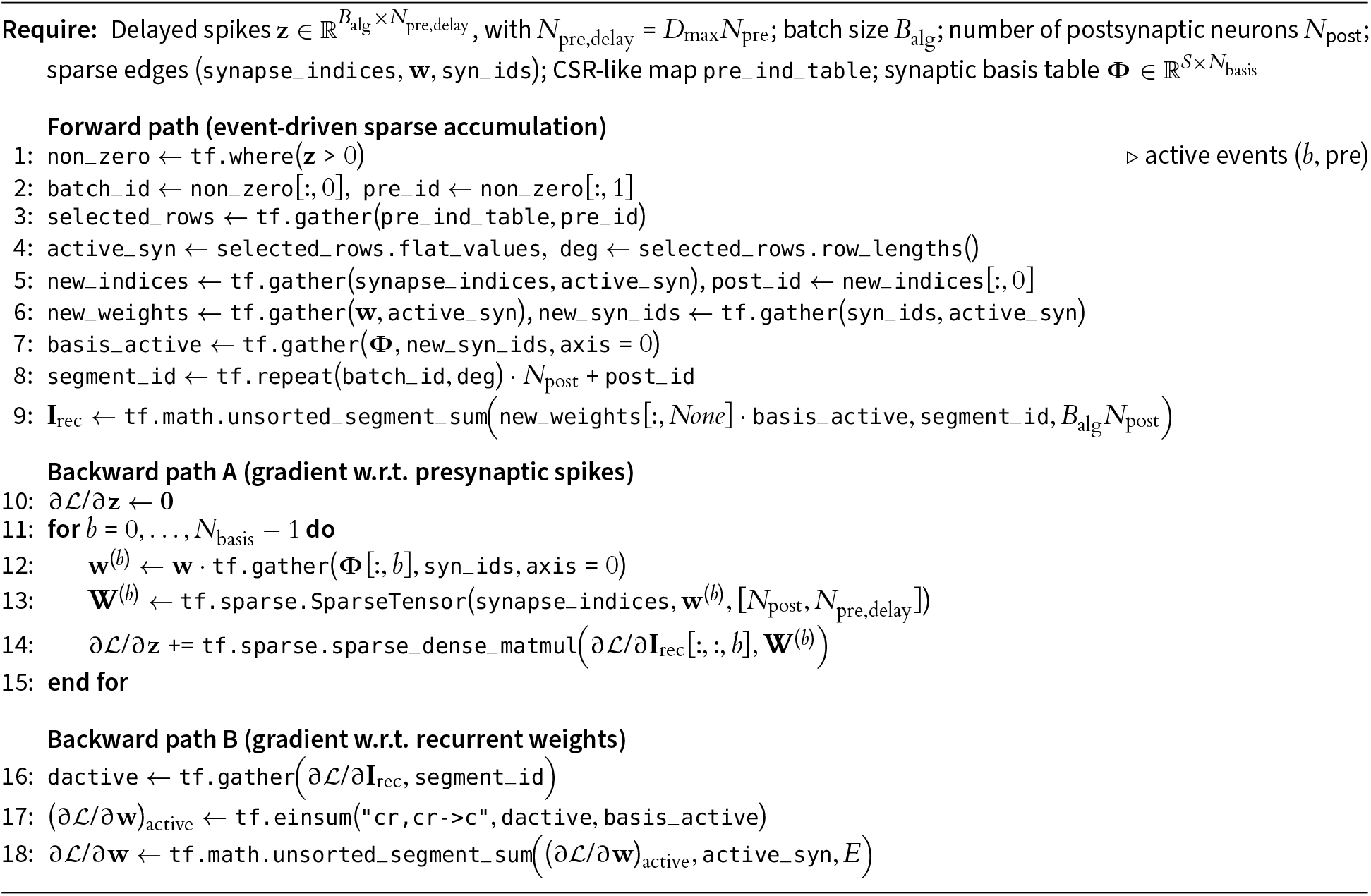

(typical firing rates 3–5 Hz, so only ~ 0.3–0.5% of neurons spike in a 1 ms step). We therefore compute recurrent synaptic drive with an event-driven sparse kernel rather than a dense recurrent matrix multiply.

At each step, only active presynaptic events are expanded through Compressed Sparse Row (CSR) format lookup tables^118^; their contributions are accumulated by segmented reductions over postsynaptic targets and synaptic basis channels (Algorithm 1). Axonal delays are handled through an augmented delay buffer: each presynaptic neuron is expanded into *D*_max_ delay lanes, and synapses with delay *d* index the corresponding lane directly. This preserves delayed transmission while keeping first-order per-step state updates.

The presynaptic lookup table is built on CPU (NumPy/Numba), where irregular indexing/bucketing is efficient, while per-step event expansion and segmented reductions run on GPU. Batches are processed jointly using flattened segment IDs^99^. TensorFlow does not provide a native sparse–sparse matrix multiplication (matmul) path for this update, so we use (i) event-driven expansion + segmented sums in the forward path and (ii) sparse-to-dense products in the backward path to obtain exact Jacobian-vector products under automatic differentiation.

#### Gradient checkpointing

All reported training runs enabled gradient checkpointing on the RSNN forward path (`tf.recompute_grad`). This reduces BPTT activation memory by recomputing selected intermediates during the backward pass instead of storing them, at the cost of additional computation^58^.

#### Mixed precision

We used mixed precision for training (`float16` compute dtype, `float32` master variables) and `float32` for testing/evaluation. The optimizer was wrapped in `LossScaleOptimizer` to reduce `float16` gradient underflow^57^. Most recurrent state tensors and per-step arithmetic run in compute dtype, while trainable weight variables are maintained in `float32` and mirrored into compute shadows after each optimizer update.

#### Graph mode and selective JIT compilation

Training and evaluation loops were implemented in TensorFlow graph mode (@tf.function)^114^. JIT compilation (`jit_compile=True`) was applied selectively to stable tensor-heavy components (e.g., stimulus construction and several loss/regularizer kernels). The recurrent sparse/ragged custom-gradient path was kept without explicit JIT compilation, and global XLA autoclustering was not enabled, because this path was not consistently robust under our sparse custom operators.

#### Global alpha-basis approximation for memory-efficient BPTT

The receptor-specific double-alpha dynamics in Eq. (12) preserve synaptic-kinetic diversity, but their naive receptor-wise implementation is memory-intensive under BPTT because all PSC states must be retained (or recomputed) across time. In the Results (Fig. 2e,f), this appears as the final memory/compression component of the training framework.

A naive receptor-wise implementation stores four PSC states per receptor channel and neuron:

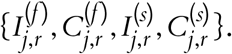

Here *j* indexes the postsynaptic neuron, *r* the receptor family, and the superscripts (*f*) and (*s*) denote the fast and slow alpha components of the double-alpha kernel (Eq. (12)). For each component, 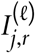 is the PSC amplitude state and 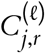 is the auxiliary convolution state that drives it (receiving weighted incoming spikes and decaying with time constant τ_*r,𝓁*_); together they implement the alpha-function shape without explicit spike-history convolution. Across the 121 receptor families described above, a given postsynaptic neuron receives input from at most 11 distinct presynaptic cell types (e.g., an L2/3 excitatory neuron can receive recurrent, LGN, and background drive from each of the 11 types), so it carries up to 11 receptor channels, yielding 44 PSC states per neuron in the naive representation. BPTT memory therefore scales prohibitively as

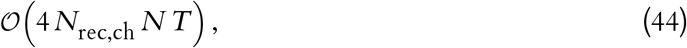

where *N*_rec,ch_ is the number of receptor channels per neuron (up to 11), *N* is neuron count, and *T* is sequence length in steps. To reduce this cost, we approximate each fitted double-alpha kernel with a shared basis of *N*_basis_ single-alpha kernels with global time constants 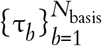. For receptor family *r*,

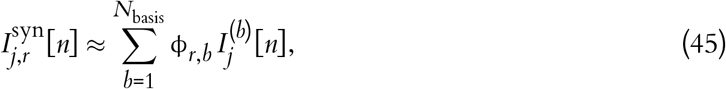

where 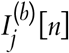 are basis PSC states shared across receptor families and ϕ_*r,b*_ are family-specific coefficients. The basis coefficients and time constants were fit by least squares across all fitted receptor families. The error distribution in Fig. 2f decreases steeply with basis size. We selected *N*_basis_ = 4, for which all fitted receptor families achieved MSE < 10^−3^ over 30 ms of kernel evolution and integrated charge transfer error < 1.7%. The selected basis time constants were

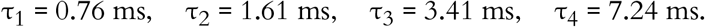

Because basis states are shared across receptor families, each neuron stores only 2*N*_basis_ PSC states (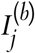 and 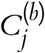), i.e. 8 states for *N*_basis_ = 4, instead of 44 in the receptor-wise implementation. PSC memory under BPTT then scales as

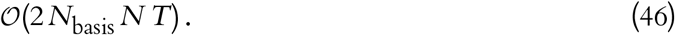

For a 200,000-neuron simulation of 2 s duration at Δ*t* = 1 ms (*T* = 2000) in `float32`, this corresponds to ~11.9 GB for *N*_basis_ = 4, compared with ~66 GB for the naive 11-channel receptor-wise double-alpha representation (5.5 × reduction). This reduction is a key enabler of single-GPU training at circuit scale while preserving receptor-dependent PSC diversity.

### Training pipeline and reporting conventions

#### Gray-screen initialization

At the start of each training epoch, the network state was re-initialized from a spontaneous (gray-screen) state. Specifically, we simulated 500 ms of spontaneous LGN drive starting from the zero state and used the resulting RSNN state as the epoch start state. This reduces state drift across long training runs and improves consistency of recurrent initial conditions.

#### Per-update simulation protocol

In each training step, gradients for the loss (Eq. (27)) across both the spontaneous or evoked chunks were computed by BPTT. Only recurrent and background synaptic weights were optimized; LGN → V1 weights were fixed throughout. Thalamocortical projections establish their coarse wiring early in development and are comparatively stable in the adult cortex^119–121^, motivating their exclusion from optimization. In contrast, intracortical recurrent connections are a primary substrate of experience-dependent plasticity^32^, and background inputs— representing aggregate drive from the rest of the brain—vary in effective strength with cortical and behavioral state^122,123^; both were therefore allowed to adapt during training.

#### Typical training configuration

Unless otherwise specified, reported training runs used:

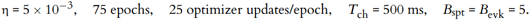

The learning rate, η, was selected by logarithmic grid search over [10^−4^, 10^−1^]. Batch size *B* was chosen to maximize GPU occupancy without exceeding available VRAM.

Training-curve metrics shown in Fig. 2b.

The absolute-loss panel reports epoch means of validation metrics for each loss component ℒ_*q*_, where *q* ∈ {rate, OSI/DSI, sync, volt, w}, and their total. Relative-loss traces are normalized by the first validation epoch *e* = 1:

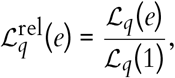

and plotted on a log scale to compare convergence rates across terms with different magnitudes.

### Computational scaling benchmarks

Reference training runtime (reported model). For the reported V1-scale models, one optimizer update (1 simulated second total: 500 ms spontaneous + 500 ms evoked) required ~ 11.8 s wall-clock time on a single NVIDIA RTX PRO 6000 GPU, including stimulus generation, forward simulation, gradient computation, and parameter update. During training, the network operated at an average firing rate of ~ 4 Hz and used ~ 28.5 GB of GPU VRAM. Full optimization required ~ 6.5 h on one NVIDIA RTX PRO 6000 GPU.

Scaling measurements used in Fig. 2d. To characterize scalability, we measured step time and VRAM as a function of network size under:

- training benchmark: *T* = 1000 ms total simulated time per update, batch size *B* = 1,
- testing benchmark: *T* = 4000 ms, batch size *B* = 10,

on one NVIDIA RTX PRO 6000 GPU and 16 AMD EPYC 9754 CPU cores. The testing configuration remained below the real-time reference line for networks < 100, 000 neurons (faster-than-biological-time inference).

Two regimes are visible (Fig. 2d): (i) a fixed-overhead regime for small networks (*N* ≤ 20k), where framework/kernel-launch overhead dominates, and (ii) an approximately linear regime beyond ~ 20, 000 neurons. For small networks, GPU utilization is low; increasing batch size improves effective throughput. To demonstrate this, we additionally benchmarked batch-size scaling on a single NVIDIA L40S GPU (48 GB VRAM) with 8 AMD EPYC 9754 CPU cores (Extended Data Fig. 3), observing sublinear growth in both step time and VRAM with increasing batch size.

#### Linear-regime extrapolation (single-GPU capacity; first-order estimate)

To estimate approximate single-GPU capacity in the linear regime (*N* ≥ 20, 000), we fit

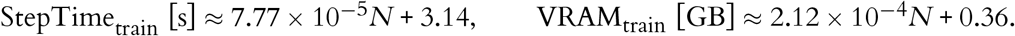

These fits provide first-order extrapolations (Table 1), but absolute numbers are hardware- and software-stack-dependent, and secondary bottlenecks (e.g., host–device transfer, kernel-launch overheads) may become important at very large *N*, where model partitioning across multiple GPUs may be preferable^99^.

**Table 1:**
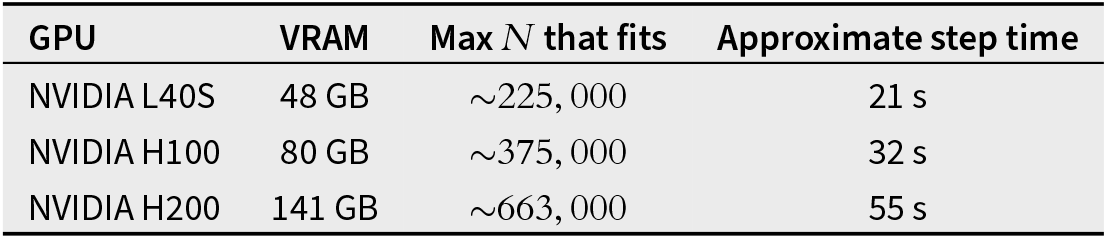
Single-GPU capacity extrapolated from the linear fit in Fig. 2d. Coefficients were estimated from benchmarks of this implementation on an NVIDIA RTX PRO 6000 (96 GB) under the same training protocol (*T* = 1000 ms, *B* = 1). Absolute step times depend on hardware generation, memory bandwidth, drivers, CUDA, and TensorFlow versions.

| GPU | VRAM | Max $N$ that fits | Approximate step time |
| --- | --- | --- | --- |
| NVIDIA L40S | 48 GB | $\sim 225,000$ | 21 s |
| NVIDIA H100 | 80 GB | $\sim 375,000$ | 32 s |
| NVIDIA H200 | 141 GB | $\sim 663,000$ | 55 s |

### Stimulus and data generation methods

Drifting gratings. Training gratings were generated directly in TensorFlow and transformed by an explicit LGN model. For spatial position (*x, y*) and time *t*,

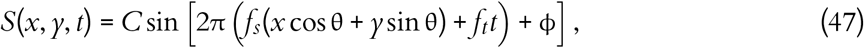

with contrast *C*, orientation θ, phase ϕ, temporal frequency *f*_*t*_, and spatial frequency *f*_*s*_ in cycles per degree. In the default generator, orientation and phase are sampled per trial (or cycled deterministically for regular-angle evaluation), then padded by pre- and post-gray periods.

#### LGN transformation and spike sampling

Stimuli are passed through spatial and temporal LGN filters to obtain firing rates in Hz. LGN spikes are sampled as Bernoulli events with

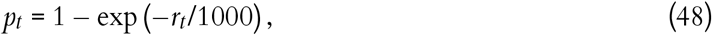

for 1 ms bins. Equivalent stateless generation is used for spontaneous gray-screen LGN activity. For natural scene stimuli (both Brain Observatory and ImageNet), the image contrast applied to the FilterNet spatial filters was doubled to achieve adequate LGN firing rates. Data pipelines are configured as deterministic with per-pipeline folded seeds to ensure reproducibility across distributed workers.

### Evaluation and analysis methods

Unless otherwise specified, all evaluation metrics, statistical analyses, and summary distributions (e.g., aggregated box plots) were computed from neurons in the core region and across an ensemble of 10 independently trained network models initialized with different random seeds. Validation used 2 s gray + 2 s gratings across 8 directions with 10 repeats/direction; natural images and contrast sweeps were held out for out-of-distribution evaluation^20,21,109^.

#### Drifting-grating response metrics

Model responses were evaluated with drifting gratings across eight directions (0° to 315° in 45° increments), with 10 trials per angle. For each neuron, preferred-direction response, OSI, and DSI were estimated from trial-averaged activity over the evoked window; spontaneous rates were estimated from pre-stimulus gray periods.

#### Comparison to experimental distributions

Population-level statistics (spontaneous rate, evoked rate, OSI, DSI, and synchronization/Fano factor profiles) were compared against Neuropixels-derived references using matched cell-type groupings. The same grouping definitions used in the loss functions were applied for evaluation summaries. To quantify the agreement between model and experimental distributions, we computed a similarity score defined as 1 − *D*_KS_, where *D*_KS_ is the Kolmogorov-Smirnov statistic between the two distributions.

#### State and activity diagnostics

During training and evaluation we tracked component losses, mean firing rates, raster summaries, and membrane-voltage traces. Power-spectrum and Fano factor analyses were computed on designated spontaneous and evoked windows for core populations. For Fig. 2c, representative rasters were rendered from checkpoints at epochs 0, 25, 50, 75 using the same validation stimulus pair, with neurons grouped by layer/class and plotted over the analysis core.

#### Orientation and direction selectivity indices

For responses to drifting gratings, the Orientation Selectivity Index (OSI) and Direction Selectivity Index (DSI) were computed using vector summation. Note that these indices are evaluated for individual neurons, which is distinct from the group-level OSI defined earlier for the loss function. Let 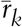 be the trial-averaged firing rate of a given neuron in response to a grating drifting at direction θ_*k*_ (in radians), where the sum runs over the eight tested directions. The DSI is defined as:

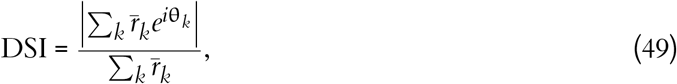

and the OSI is calculated using double-angle vectors:

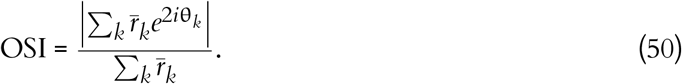

To estimate each neuron’s preferred direction (PD), we used these same trial-averaged drifting-grating responses, averaged over 10 trials per direction. Using the same vector sums as in Eqs. 49 and 50, we define

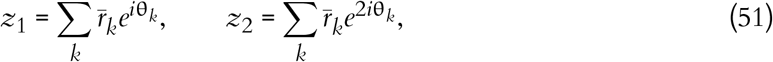

where *z*_1_ and *z*_2_ are the first- and second-order circular moments of the tuning curve. The preferred orientation was obtained from the phase of the second moment as

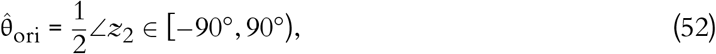

and then disambiguated to a full 360° preferred direction using the sign of the projection of *z*_1_ onto the orientation axis:

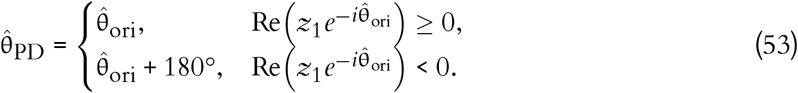

Only neurons with peak-direction mean firing rate ≥ 1 Hz across the eight directions were included in downstream preferred-direction analyses.

#### Image/Stimulus selectivity (response sparsity)

Selectivity to natural images was quantified using response sparsity across the 118 Brain Observatory images^20^. For a given neuron, let *r*_*i*_ be the trial-averaged response to the *i*-th image and *N* = 118 the total number of images. The image selectivity *S* is defined as:

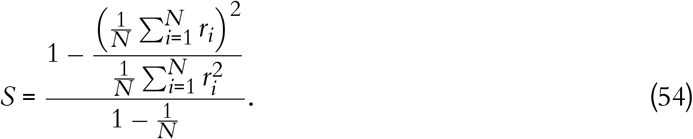

A value of 0 indicates equal responses to all images, while a value of 1 indicates a response to only a single image.

An analogous stimulus selectivity was computed using responses to drifting gratings. For a given neuron, let *r*_*j*_ be the trial-averaged firing rate in response to the *j*-th direction of motion and *N* = 8 the number of directions tested. The stimulus selectivity is computed using the same formula, yielding a value of 0 when the neuron responds equally to all directions and 1 when it responds to only a single direction.

#### Contrast response simulations

To evaluate contrast-dependent modulation, trained networks were presented with drifting gratings at varying contrast levels (*C* ∈ {0.05, 0.1, 0.2, 0.4, 0.6, 0.8}) while keeping spatial and temporal frequency fixed at 0.04 cycles per degree (cpd) and 2 Hz, respectively. Firing rates, OSI, and DSI were computed at each contrast level per neuron and then aggregated by cell type. These simulations were not included in training (which used only high-contrast gratings at *C* = 0.8) and thus serve as an out-of-distribution generalization test.

#### Network stability and participation ratio

To monitor network stability during optimization and detect pathological regimes—such as runaway excitation, extreme suppression, or transient synchronized events—we tracked the mean population firing rate and the maximum participation ratio at the end of each epoch (Extended Data Fig. 2). Both metrics were computed from 5 independent 500 ms trials for both spontaneous and evoked conditions.

The participation ratio is defined as the fraction of active neurons within a discrete time window. To avoid obscuring brief bursts of activity, we analyzed the maximum participation ratio across non-overlapping time bins of width Δ*t* ∈ {10, 50, 100} ms. For bin *b*, the fraction of active neurons is:

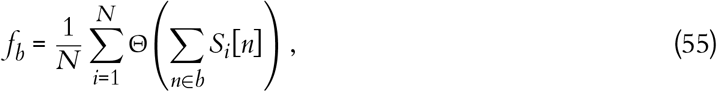

where *N* is the total number of neurons, *S*_*i*_[*n*] ∈ {0, 1} is the spike indicator of neuron *i* at simulation step *n* (1 ms resolution), as defined in Eq. 8, and Θ(*x*) is the Heaviside step function (Θ(*x*) = 1 for *x* > 0, and 0 otherwise). We report the mean and standard deviation of the maximum *f*_*b*_ over all bins, computed across the 5 trials.

#### Most exciting inputs (MEIs)

To probe the features driving individual model neurons, we generated most exciting inputs (MEIs) following the approach and CNN architecture of Walker et al.^70^. For each neuron, a convolutional neural network was first trained to predict the neuron’s trial-averaged responses to natural images. The CNN was then used as a differentiable surrogate, and gradient ascent in the stimulus space was performed to discover synthetic images that maximally excite each neuron.

ImageNet stimuli consisted of 5000 images presented once and 100 images repeated 10 times, with each image shown for 250 ms followed by a 250 ms gray screen inter-stimulus interval, matching the protocol of Walker et al.^70^. Response reliability was quantified by an oracle-correlation metric computed from the repeated ImageNet presentations.

For a given neuron *i*, let *r*_*i,g,k*_ denote its response (spike count) to the *k*-th presentation (*k* ∈ {1, …, *K*}) of the *g*-th unique repeated image (*g* ∈ {1, …, *G*}), where *K* = 10. We adopted a leave-one-out approach to compute the mean response to the same image, excluding the current trial. For each trial *k*, the leave-one-out average response is given by:

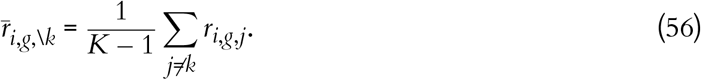

The single-trial responses and their corresponding leave-one-out averages are then vectorized across all repeated images and trials, yielding two vectors **r**_*i*_ and 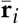, each of length *G* × *K*. The oracle correlation ρ_*i*_ for neuron *i* is defined as the Pearson correlation coefficient between these two vectors:

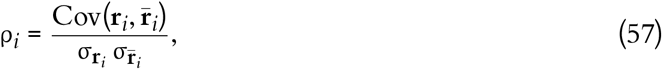

where Cov(·, ·) denotes the sample covariance and σ denotes the sample standard deviation.

For MEI generation, we selected a total of 2000 neurons among those with the highest 30% oracle correlation within each cell type, pooled across all 10 trained model networks.

To characterize the spatial structure of MEIs, we computed the two-dimensional spatial power spectrum for each MEI and summarized each neuron’s spectral profile by the ratio of high-frequency power (0.08–0.16 cycles per degree) to low-frequency power (0.02–0.04 cycles per degree).

### Synaptic weight analysis

To explore the emergence of functional microcircuits following optimization, we systematically analyzed the relationship between trained synaptic weights and the functional tuning properties of pre- and post-synaptic neuron pairs. Note that the presence or absence of connections is not modified by our training procedure; only synaptic weights are adjusted. The analyses were conducted across the 19 × 19 cell-type connectivity matrix, as well as in aggregated form (layer segregated excitatory types and layer aggregated inhibitory types).

#### Response correlation analysis

For each connected neuron pair in the network, we calculated the Pearson correlation coefficient of their trial-averaged responses to the 118 Brain Observatory natural images^20^. Synaptic weights were then grouped by these response correlations into three functional domains: anti-correlated ([−1, −0.5]), uncorrelated (or very weakly correlated) ([−0.25, 0.25]), and highly correlated ([0.5, 1.0]). To quantify the strength of like-to-like connectivity, we computed the normalized weight differences relative to the “uncorrelated” baseline:

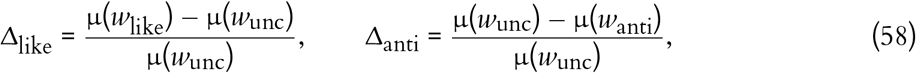

where µ(*w*_like_), µ(*w*_anti_), and µ(*w*_unc_) denote the mean synaptic weight in the like-correlated, anti-correlated, and uncorrelated domains, respectively. By this definition, positive values for both Δ_like_ and Δ_anti_ indicate a like-to-like connectivity structure (stronger connections for similarly responding neurons and weaker connections for oppositely responding neurons, relative to the uncorrelated baseline), whereas negative values indicate the opposite—that is, an anti-like-to-like structure. Note that the above is true for both excitatory and inhibitory connections. The latter have negative weights. Thus, e.g., stronger inhibition for highly correlated vs. uncorrelated neurons would result in dividing a negative µ(*w*_like_) − µ(*w*_unc_) by a negative µ(*w*_unc_), giving a positive result for Δ_like_ and indicating like-to-like inhibition.

To avoid conflating laminar targeting preferences with functional tuning relationships, the response correlation analysis for inhibitory cell types (PV, SST, VIP) shown in Figs. 5, 6 was restricted to within-layer connections—that is, inhibitory neurons were paired only with excitatory neurons from the corresponding cortical layer. An exception was made for L1 Inh neurons, which were paired with excitatory neurons from all layers, as L1 contains no excitatory cells. Cell-type pairs for which no neuron pairs with sufficient response correlation (< − 0.5 or > 0.5) were excluded from the analysis, and the corresponding Δ metrics were not calculated.

#### Preferred-direction difference analysis

For each synapse connecting presynaptic neuron *i* to post-synaptic neuron *j*, we computed the circular preferred-direction difference as

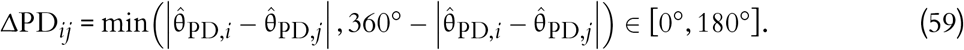

Synaptic weights were binned by ΔPD_*ij*_, and the relationship was quantified using a cosine-series fit of the form:

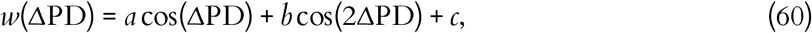

where *a* captures direction-specific like-to-like structure, *b* captures orientation-specific like-to-like structure, and *c* is the baseline offset. The normalized effect sizes *a*/*c* and *b*/*c* are reported, with positive values indicating like-to-like and negative values indicating anti-like-to-like relations (again, this applies to both excitatory and inhibitory connections). The cosine-series parameters were estimated by ordinary least squares, and significance of individual coefficients was assessed with two-sided *t*-tests (see Statistics).

#### Comparison to electron microscopy data

To directly compare model predictions to anatomy, we used the V1DD electron microscopy dataset^31^, which links functional tuning differences (including responses to drifting gratings) to the area of the post-synaptic density (PSD) as a proxy for synaptic strength (or the sum of PSD for multisynaptic connections). We restricted our analysis to neurons responsive to 1 Hz, full-field drifting gratings. Neurons were classified as responsive if their mean response to the preferred direction was >2.5 standard deviations above the activity during a spontaneous period. We selected neurons with reconstructed axons and identified all connections to other responsive neurons. Because the V1DD dataset provides a relatively small and fixed set of reconstructed connections (with functional tuning data available only for excitatory neurons), we employed a Monte Carlo resampling approach to match sample sizes between model and experiment. For each pathway, we repeatedly redrew the same number of connections as available in the V1DD data from our model and recomputed the tuning-dependent statistic and its associated *p*-values. This procedure was repeated 100 times to yield a distribution of model *p*-values under matched sample sizes, which was compared against the single *p*-value obtained from the experimental measurements.

### Outgoing weight cohort analysis

To investigate functional heterogeneity within projection pathways, neurons from each of the 19 cell types were grouped into three cohorts—low (bottom tertile), mid (middle tertile), and high (top tertile)—based on their total absolute outgoing synaptic weight summed across all postsynaptic targets in the entire network (both core and periphery). Cohort assignment was performed independently within each cell type (e.g., L4_Exc, L5_ET, L6_VIP), ensuring that each cell type contributed approximately one-third of its neurons to each cohort. To avoid boundary artifacts, the selection pool for the cohorts was strictly limited to source neurons residing within the 200 µm analysis core 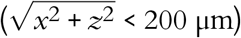.

Following grouping, the firing rates and OSI were compared between 10 instances of the trained and untrained networks. The target preferences of each cohort were characterized by computing the proportional distribution of outgoing synaptic weight across postsynaptic cell types (Exc, PV, SST, VIP).

We focused on characterizing the relation between functional properties and outgoing weights rather than incoming weights, because we did not observe a strong, training-shaped relationship between total incoming synaptic weights and functional metrics. An analogous cohort analysis based on total incoming synaptic weights was also performed and is reported in Extended Data Figure 9.

### Simulated perturbations

To establish causal links between specific network sub-populations and large-scale network dynamics, we performed targeted silencing simulations. Silencing was implemented by injecting a constant hyperpolarizing current ( − 1000 pA) into the targeted neurons for the entire duration of the visual stimulus presentation.

#### Cohort-specific silencing

To dissect the distinct roles of high-impact versus low-impact projection streams, we selectively silenced either the high-outgoing-weight or low-outgoing-weight cohorts of excitatory or inhibitory populations within the core network. Cohort definitions followed the same tertile-based grouping described above. For each suppression condition, the DGs with 8 directions × 10 trials (0.04 cpd, 2 Hz) were presented.

#### Inhibitory cell-type-specific cohort suppression

We also evaluated the distinct, network-wide regulatory impacts of specific inhibitory interneuron types. We selectively silenced either the high-outgoing-weight or low-outgoing-weight cohorts of one specific inhibitory type (PV, SST, or VIP) at a time.

#### Perturbation analysis

The functional consequences of these perturbations were evaluated by computing the percentage change in firing rate and OSI relative to the unperturbed baseline. Neurons directly targeted for suppression were excluded from the post-perturbation metric calculations. This exclusion ensured that the resulting heatmaps represented indirect, trans-synaptic, network-wide effects rather than the trivial suppression of the targeted neurons. The unperturbed baseline was computed from the same non-targeted population to ensure a matched comparison.

#### Synaptic weight distribution constraints

In the constrained run, synaptic weights were initialized by sampling from the experimentally derived distribution as described above, and training was performed with the full multi-objective loss including the weight regularization term (ℒ_*w*_) based on the Earth Mover’s distance. In the unconstrained run, we initialized weights uniformly (all excitatory connections at the same value and all inhibitory connections at the same value), and we removed the weight regularization term (ℒ _*w*_) from the loss function. This comparison therefore removes the synaptic-weight prior during both initialization and training, allowing synaptic weights to evolve without distribution constraints (Extended Data Fig. 14).

Oracle score calculation in MICrONS data.

We calculated the response reliability of layer 5 (L5) neurons from the MICrONS electron microscopy dataset following their method^22^. Namely, co-registered neurons were selected by excluding the top 25% of residuals and the bottom 25% of separation scores. Oracle scores were calculated as the mean leave-one-out Pearson correlation across ten repeated stimulus presentations: for each neuron, let **r**_1_, **r**_2_, …, **r**_10_ denote the response vectors across stimulus conditions for the ten repeats. The oracle score is then 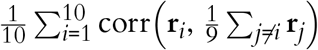, where corr(·, ·) denotes Pearson correlation.

### Statistics

For comparisons of oracle scores across L5 subtypes (Extended Data Fig. 5a), the Kruskal–Wallis test was used, followed by pairwise comparisons with Bonferroni correction for three pairs. For the cosine-series fits of synaptic weight versus preferred-direction difference (Figs. 5, 6; Extended Data Fig. 7), the model *w* = *a* cos θ + *b* cos 2θ + *c* was fit by ordinary least squares. Significance of the coefficients *a* (direction tuning) and *b* (orientation tuning) was assessed with two-sided *t*-tests on each coefficient (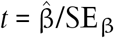, df = *n* −3), where the standard errors were obtained from 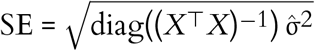, with *X* the design matrix and 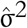 the residual mean square. For the weight-versus-response-correlation analyses (Extended Data Fig. 8), significance of the difference metrics Δ_like_ and Δ_anti_ was assessed by comparing synaptic weights in the like-correlated or anti-correlated group against the uncorrelated group using a two-sided Welch’s *t*-test.

### Software and hardware

The model was constructed using BMTK^37^ with the NEST simulator 3.6^124,125^, employing the SONATA data format^35^. MICrONS data were accessed via the CAVEclient interface^22^. The SONATA network files produced by this construction pipeline serve as input to the TensorFlow-based^114^ differentiable simulator.

All differentiable simulations and training were implemented in TensorFlow 2.15^114^. The TensorFlow implementation operates on the SONATA-derived connectivity and cell-parameter files generated by the BMTK/NEST construction pipeline, enabling direct reuse of the biologically grounded network specification in the gradient-based training workflow.

Benchmarks reported in Fig. 2d were run on a single NVIDIA RTX PRO 6000 GPU (96 GB VRAM) with 16 AMD EPYC 9754 CPU cores. Benchmarks reported in Extended Data Fig. 3 were run on a single NVIDIA L40S GPU (48 GB VRAM) with 8 AMD EPYC 9754 CPU cores. Final production training runs were performed on a high-performance computing cluster using a single NVIDIA A100 GPU (40 GB VRAM) and Intel Xeon Gold 6330N processors.

Simulations for model characterization (Figs. 3–6) were done with BMTK FilterNet and PointNet (with NEST simulator 3.6^125^).

#### Use of large language models

GPT (OpenAI), Gemini (Google), and Claude (Anthropic) were used to assist with editing manuscript text for clarity and generating code templates. All scientific content, data analyses, and conclusions are the sole responsibility of the authors.

---

**Extended Data Figure 1:**
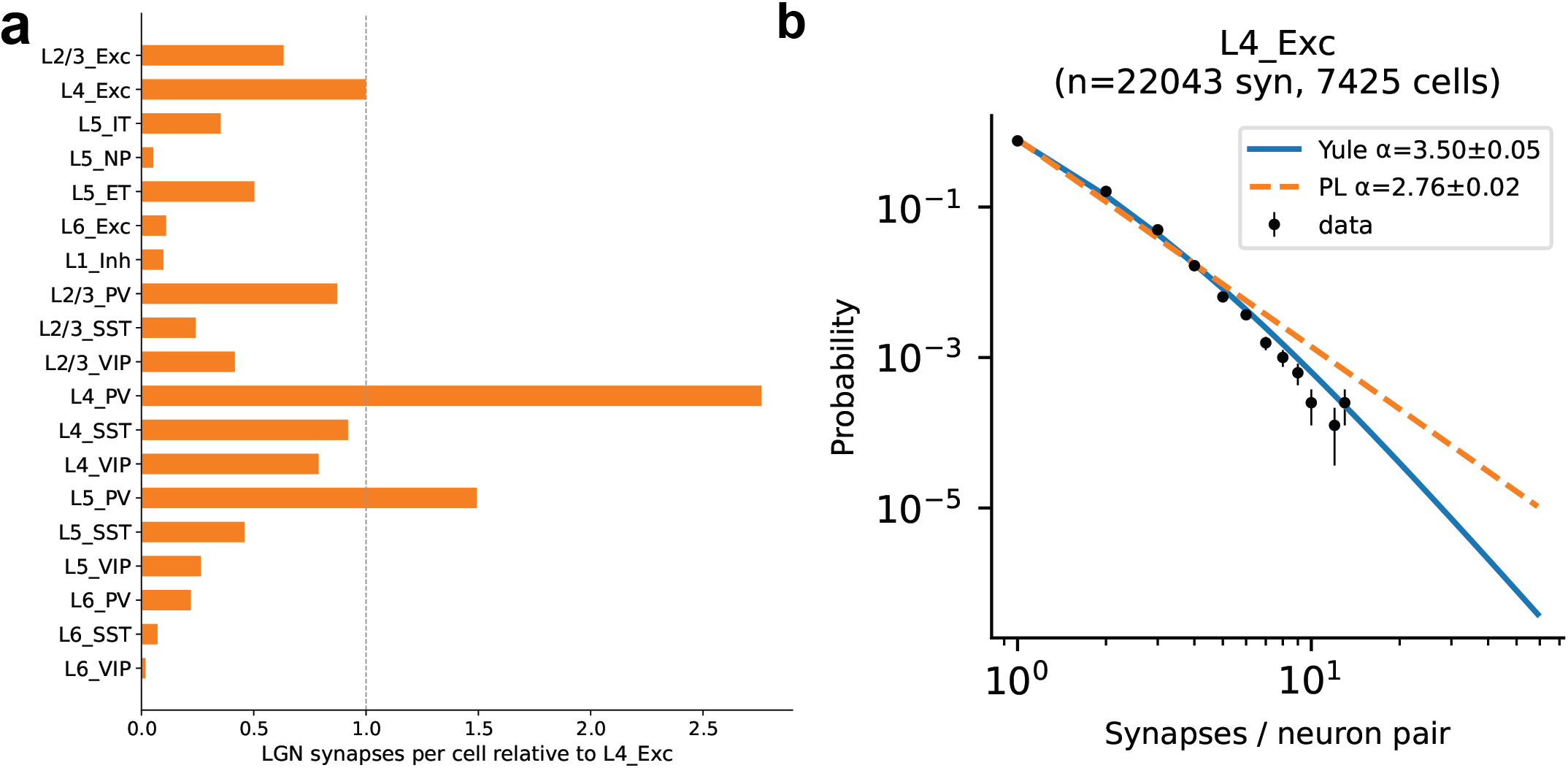
**a**, LGN synapse count per cell for each cell type relative to L4 Exc population. **b**, Per-connection synapse count distribution follows a Yule–Simon distribution. An example Yule–Simon distribution fit (blue solid line) to a per-connection synapse count distribution between LGN axons and L4 pyramidal neurons. Yule–Simon distribution fits better than the power-law distribution (orange dashed line).

**Extended Data Figure 2:**
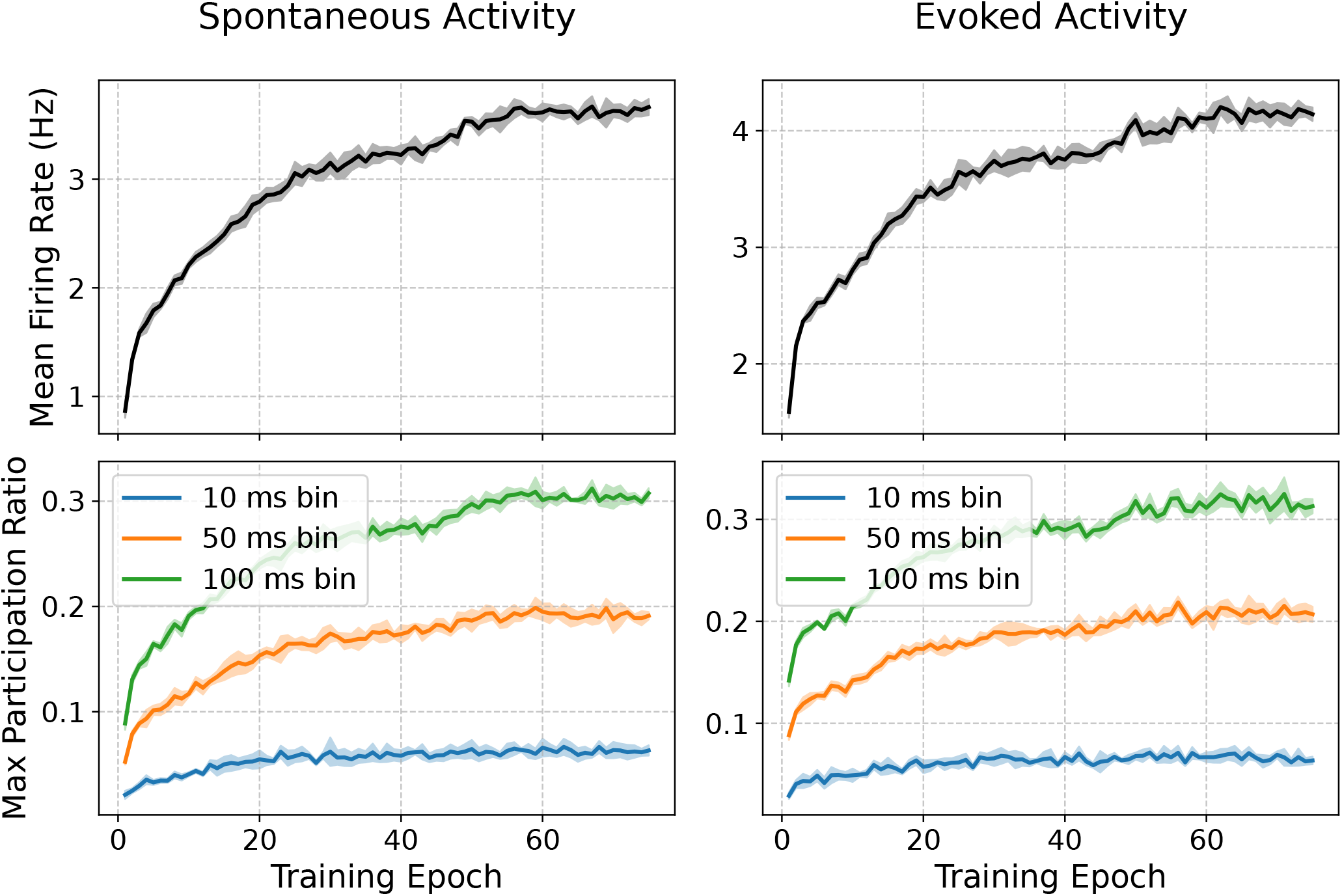
Firing rate and participation ratio during training. Mean firing rate and maximum participation ratio (fraction of neurons that fired in a bin) across 500 ms simulations are shown for spontaneous (left) and evoked activity (right). The result is from a single representative network. The shading indicates standard deviation across 5 stimulus trials.

**Extended Data Figure 3:**
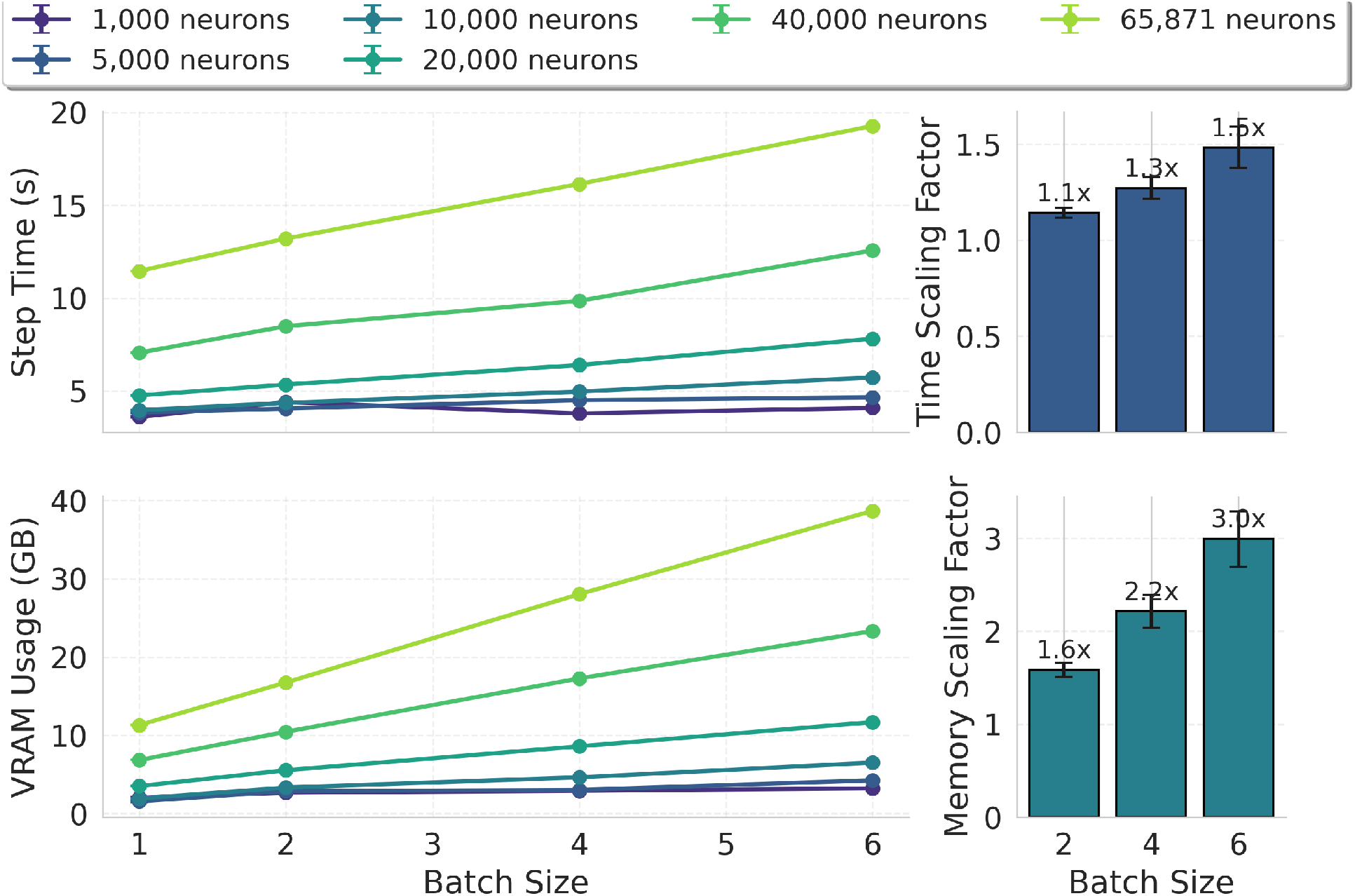
Batch-size scaling of training time and memory. Benchmarks were run with the TensorFlow simulator for a 1000 ms (500 ms of gray screen and 500 ms of drifting gratings) training step. Left panels show absolute step time and GPU memory versus batch size for multiple network sizes (mean *±* SEM over 3,000 runs). Right panels report scaling factors normalized to batch size *B* = 1 (mean *±* SEM over the 6 network configurations shown above). Increasing batch size from 1 to 6 produced sublinear step-time growth (about 1.5*×* slower) while VRAM also increased sublinearly, defining the efficient batch size processing in our implementation.

**Extended Data Figure 4:**
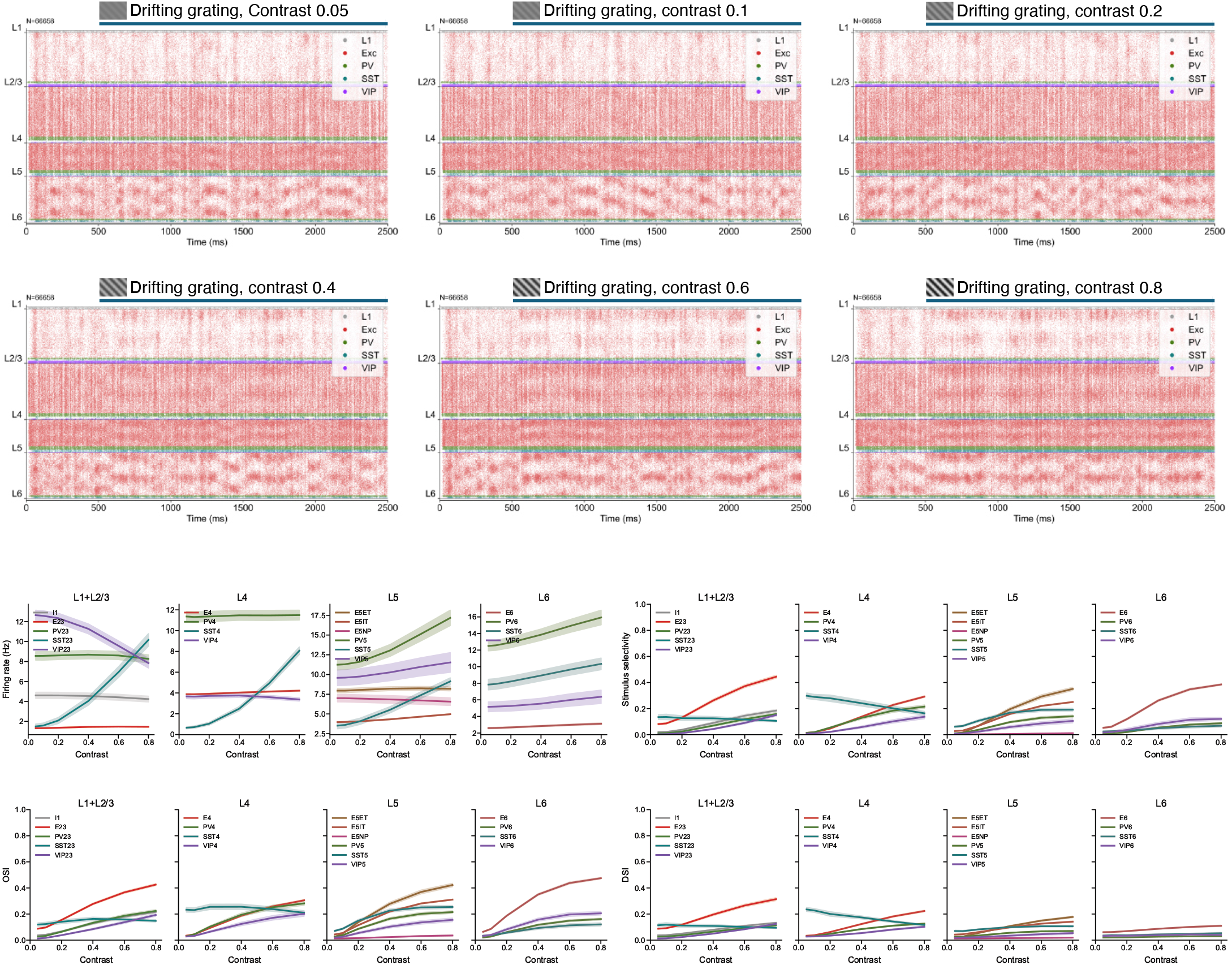
Contrast response. Raster plots of a representative network at different contrast levels are shown for 45° drifting gratings (top). Line plots show firing rates, stimulus selectivity, orientation selectivity index (OSI), and direction selectivity index (DSI) (bottom) for all cell types. Lines show the mean value across *N* = 10 networks, where each network contributes the population-averaged response per cell type. Shaded regions indicate 95% confidence intervals across networks (t-distribution).

**Extended Data Figure 5:**
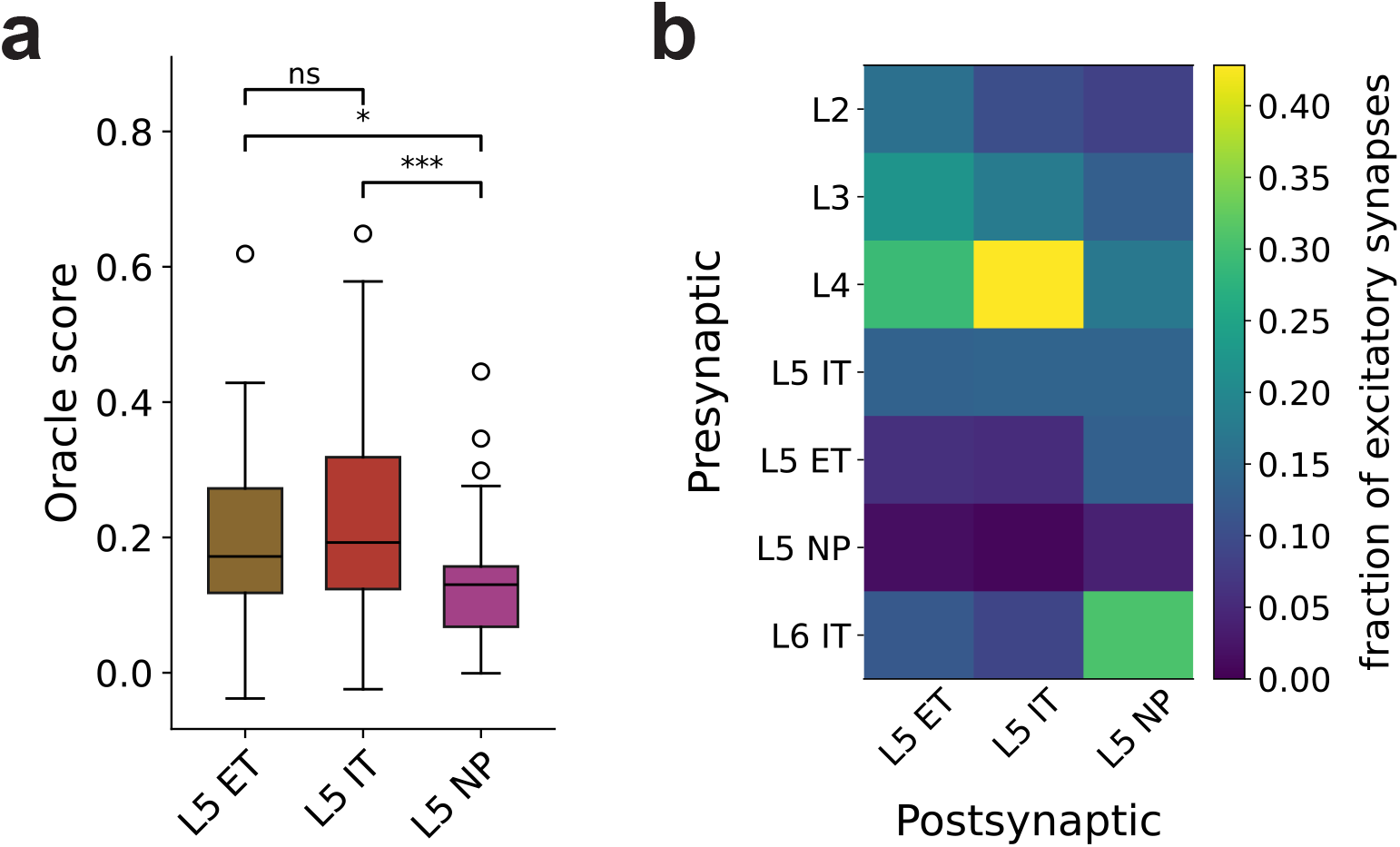
Oracle scores and input connectivity of layer 5 neurons in MICrONS. **a**, Oracle scores. Statistical significance was assessed using the Kruskal–Wallis test followed by pairwise comparisons with Bonferroni correction (*n* = 3 pairs). Stars indicate significance levels (* *p* < 0.05, *** *p* < 0.001). **b**, Fraction of input synapses from excitatory cell types (L5 ET, *n* = 77; L5 IT, *n* = 106; L5 NP, *n* = 19).

**Extended Data Figure 6:**
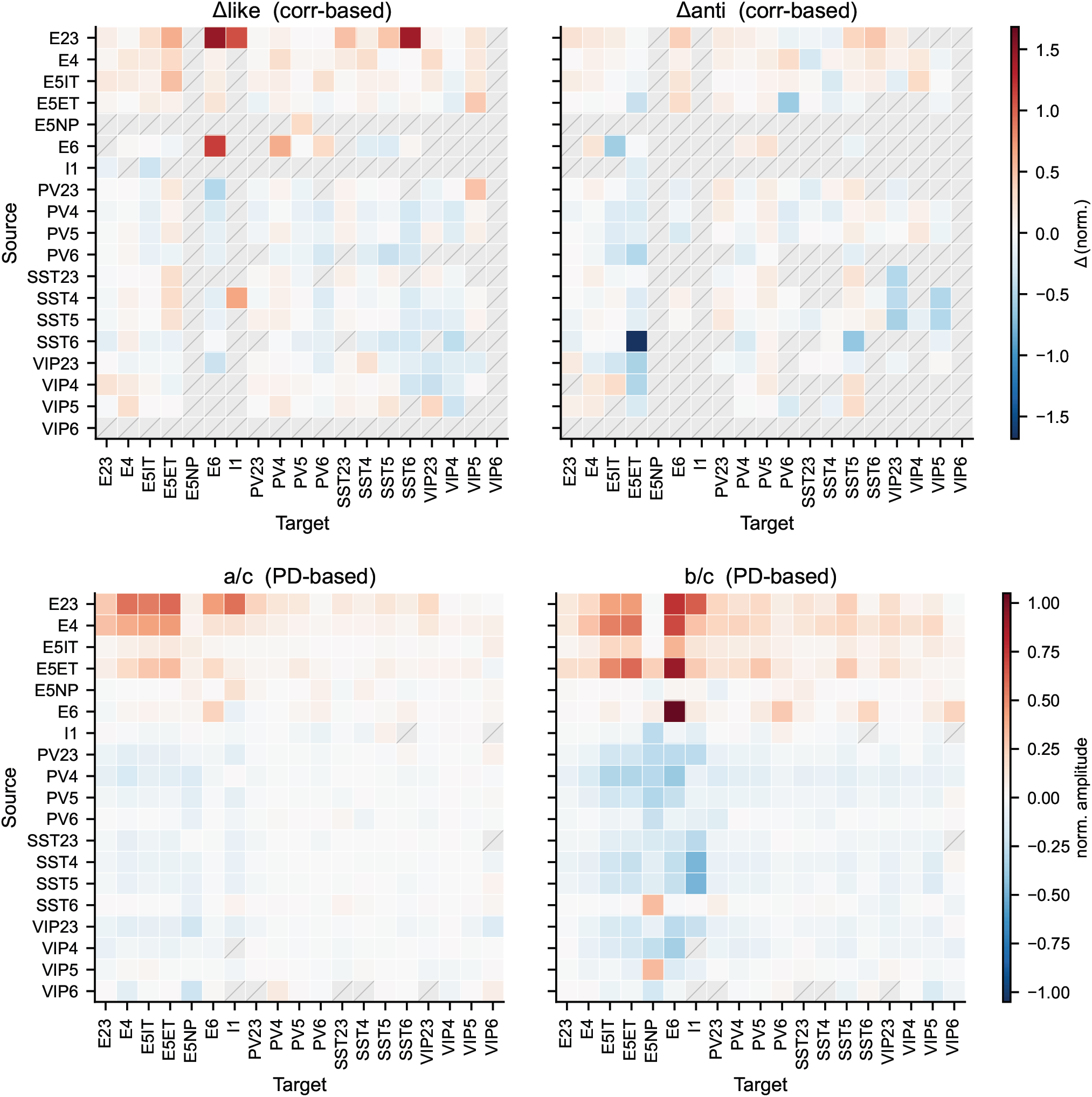
Like-to-like metrics. Full heatmaps of the like-to-like metrics are shown for all cell-type pairs. The correlation-based metrics (Δ_like_, Δ_anti_) quantify how synaptic weight varies with the Pearson correlation of responses to natural images between connected neuron pairs. The preferred-direction (PD) based metrics (*a*/*c, b*/*c*) quantify how synaptic weight varies with the difference in preferred direction of motion estimated from drifting grating responses (see Methods). The results are the mean of *N* = 10 networks.

**Extended Data Figure 7:**
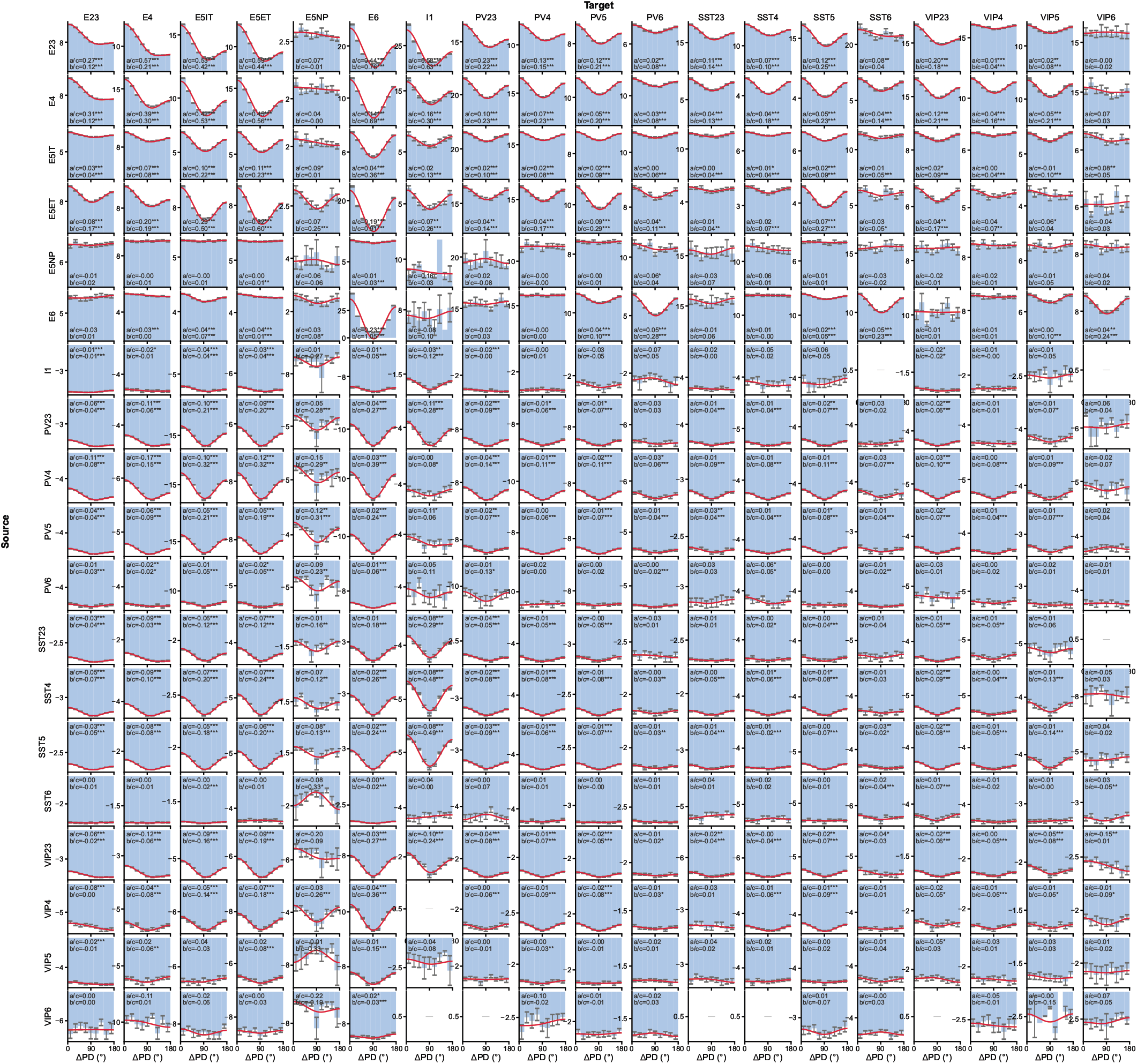
Like-to-like characterization with DGs. Full histograms of synaptic weight as a function of difference in preferred-direction for DG responses are shown for all cell-type pairs. The *y*-axis is the average weight (pA). Significance of cosine-fit coefficients was assessed with two-sided *t*-tests (see Methods). Stars indicate significance levels (* *p* < 0.05, ** *p* < 0.01, *** *p* < 0.001). The results are based on aggregated data from *N* = 10 networks.

**Extended Data Figure 8:**
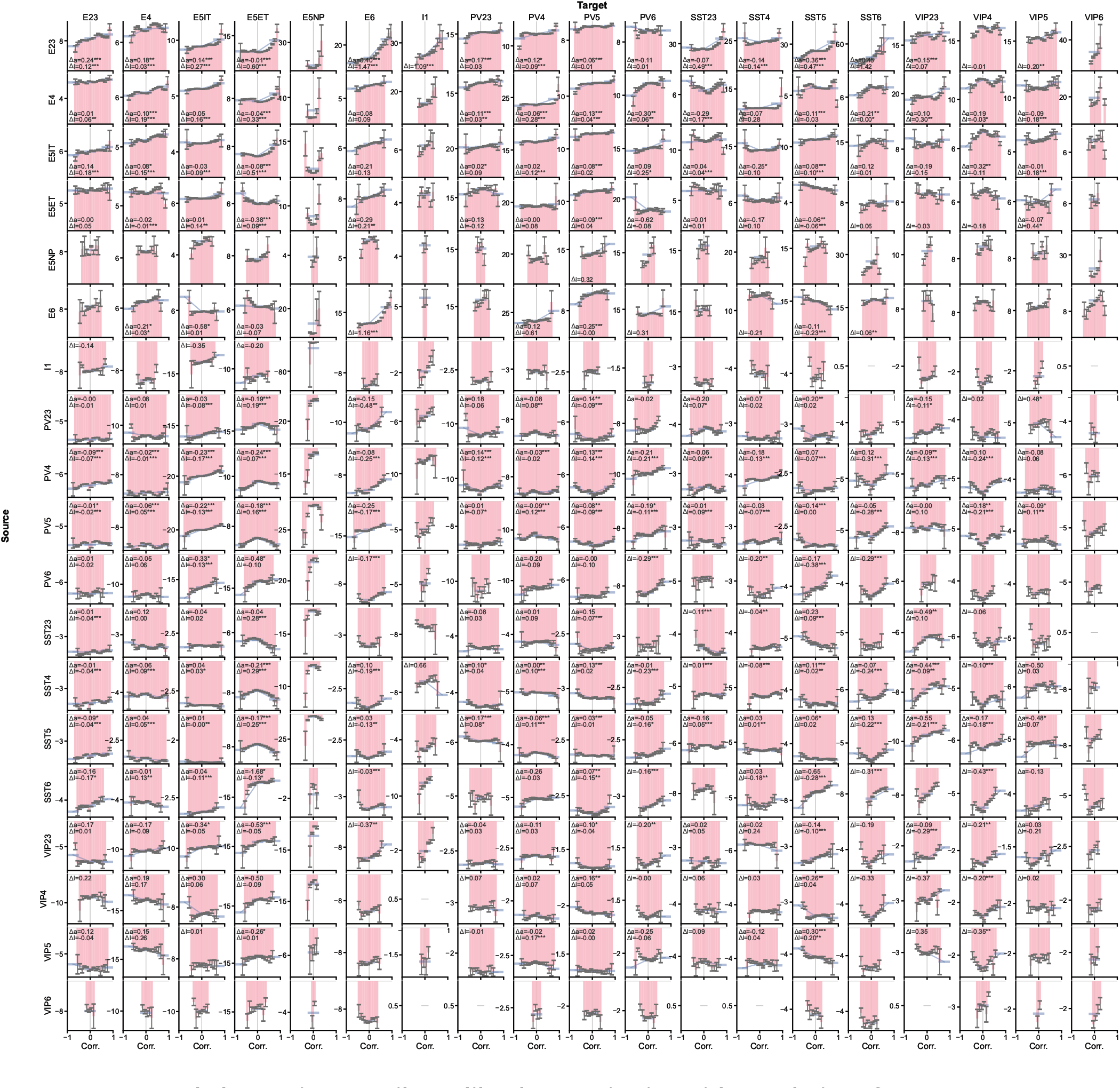
Like-to-like characterization with correlation of responses to natural images. Full histograms of synaptic weight as a function of correlation of responses to Brain Observatory natural images are shown for all cell-type pairs. The *y*-axis is the average weight (pA). Significance of the difference metrics was assessed with a two-sided *t*-test (see Methods). Stars indicate significance levels (* *p* < 0.05, ** *p* < 0.01, *** *p* < 0.001). The results are based on aggregated data from *N* = 10 networks.

**Extended Data Figure 9:**
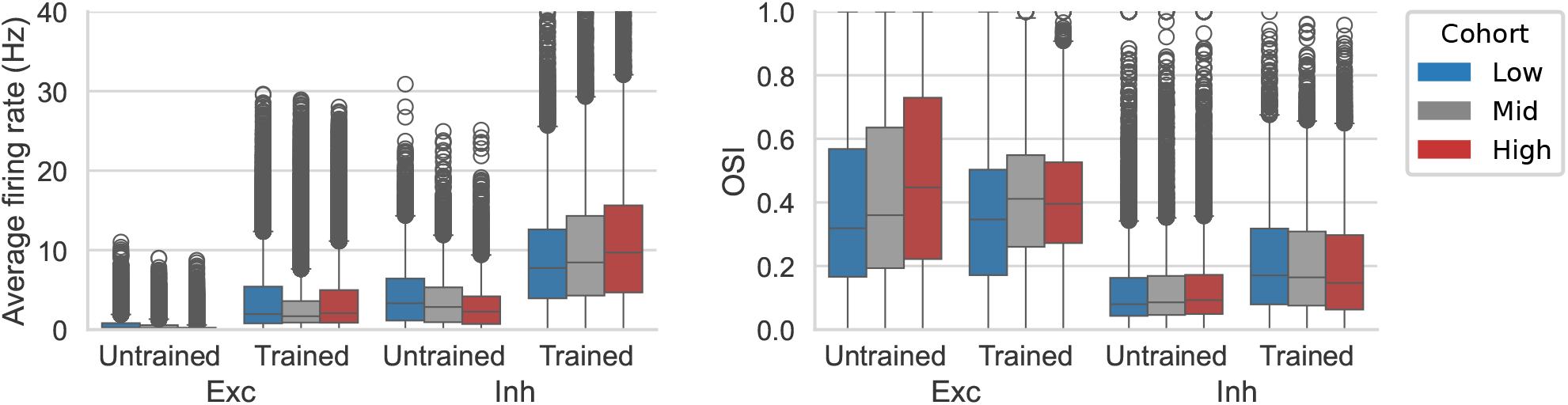
Cohorts by incoming synaptic weights. Average firing rates and orientation selectivity index (OSI) for cohorts separated by total incoming synaptic weight. The results are based on aggregated data from *N* = 10 networks.

**Extended Data Figure 10:**
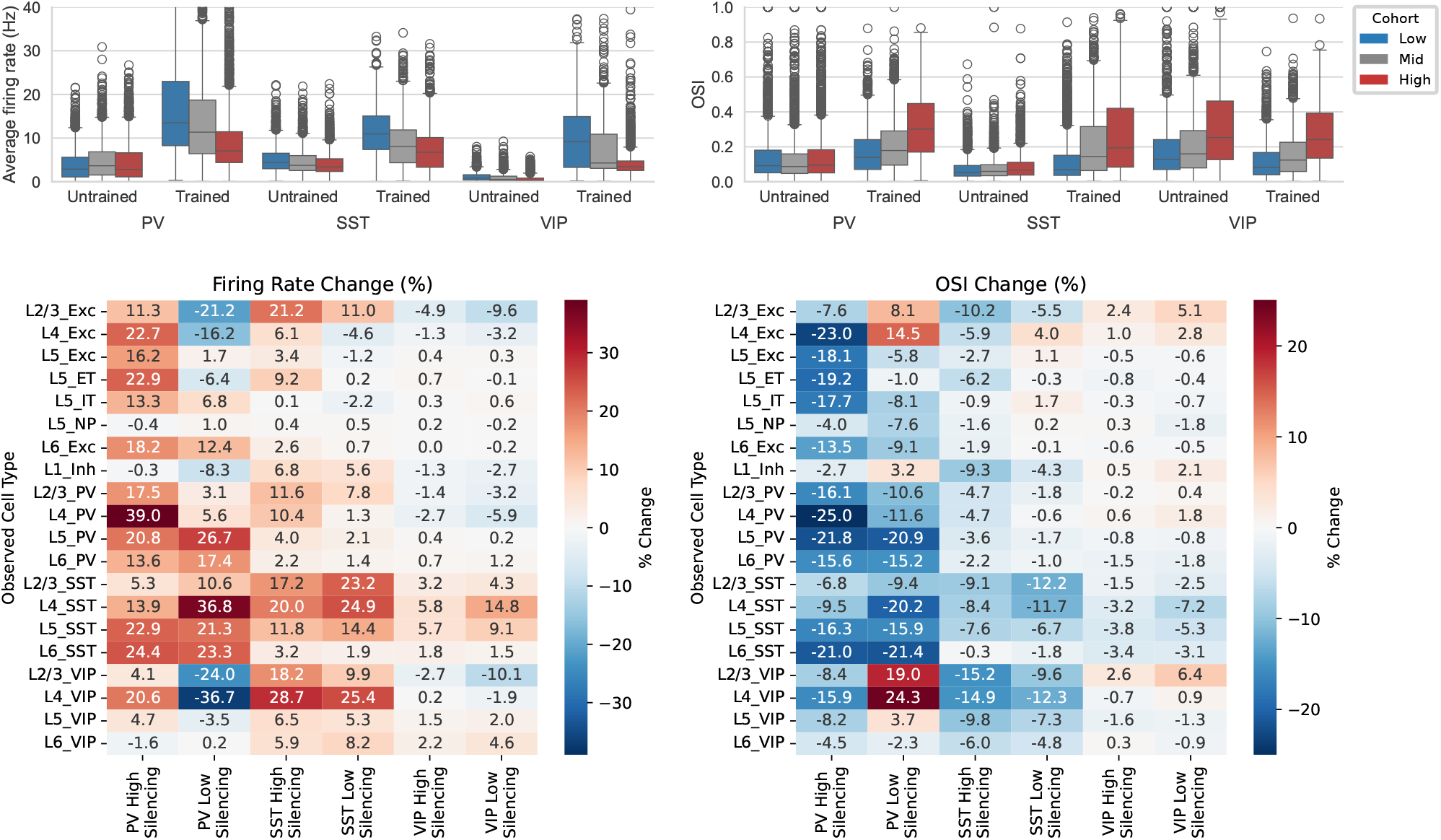
Cell-type cohort suppression analysis. Top: functional features (left: firing rates, right: OSI) of the cohorts stratified by outgoing weights for inhibitory subtypes. Bottom: The effect of suppression of the subtype cohorts on the firing rates and OSI. The results are based on aggregated data from *N* = 10 networks.

**Extended Data Figure 11:**
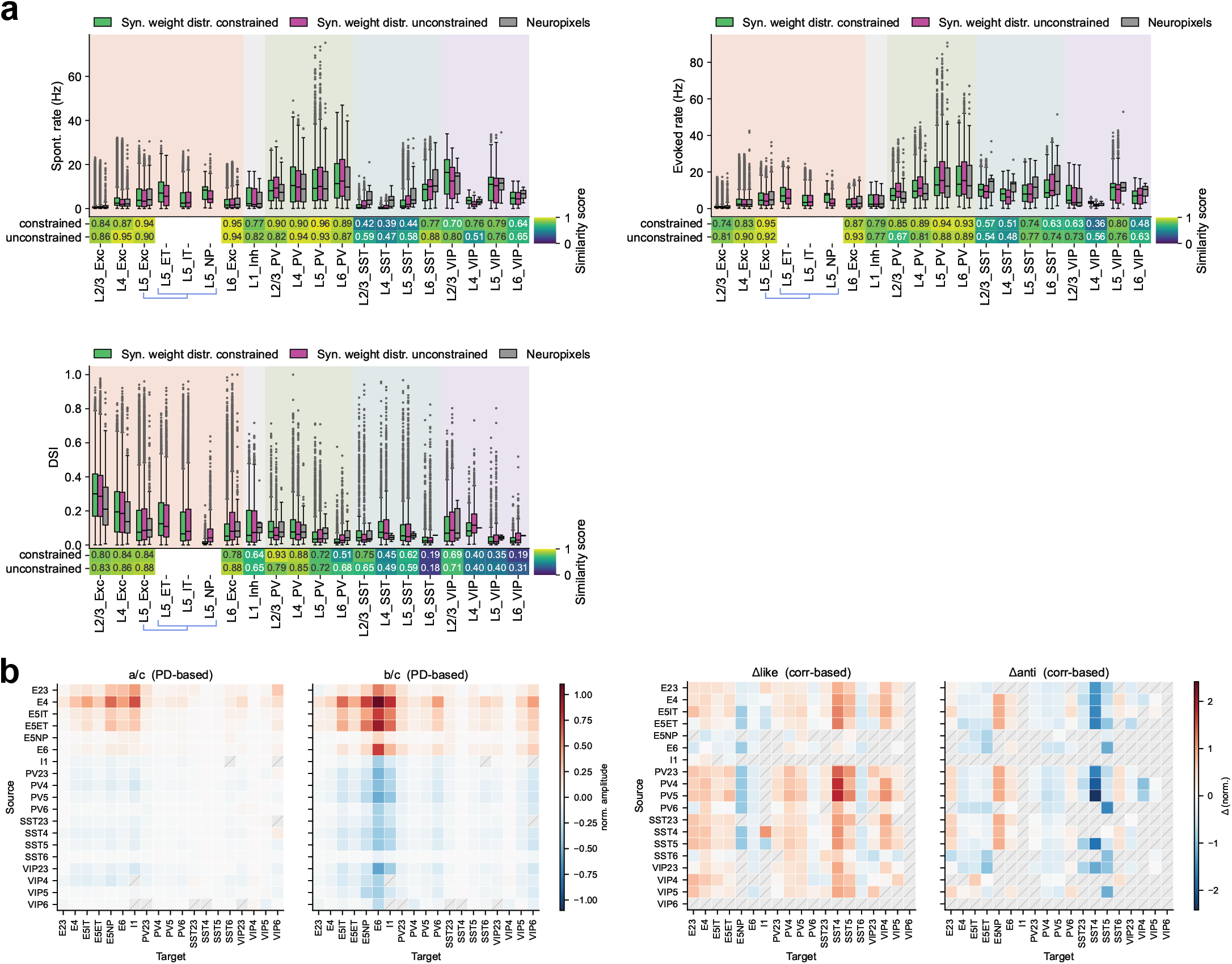
Additional metrics for networks trained without synaptic weight distribution constraints. **a**, Additional metrics (spontaneous firing rates, evoked firing rates, and direction selectivity index) for unconstrained networks. **b**, Full heatmaps of like-to-like metrics for unconstrained networks.

**Extended Data Figure 12:**
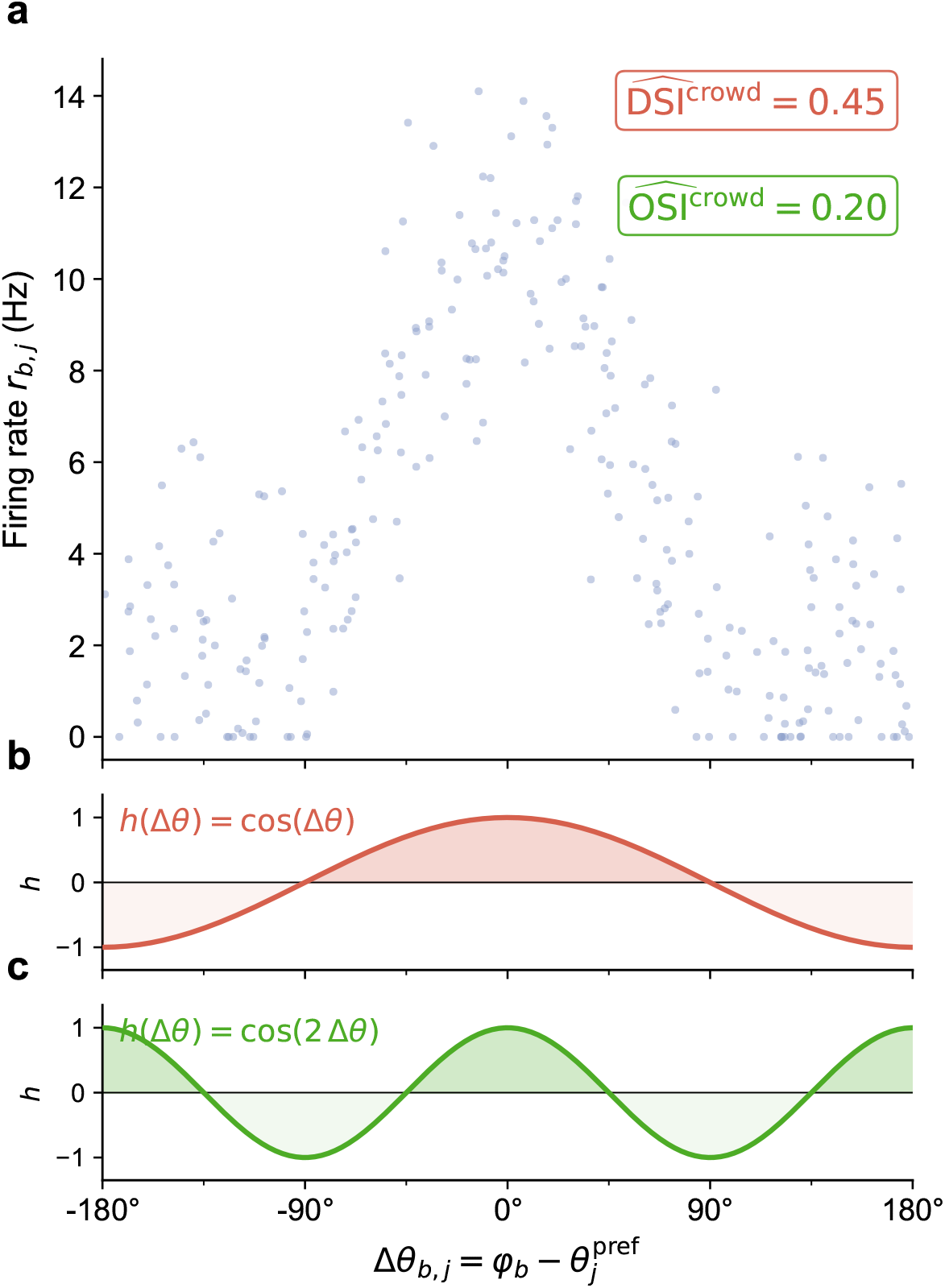
Crowd-surrogate OSI/DSI loss. **a**, Synthetic illustration of the crowd-surrogate OSI/DSI loss. Each dot represents the single-trial firing rate *r*_*b,j*_ of one model neuron, plotted against its angular offset from the presented grating direction, 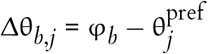. Because preferred directions are uniformly distributed within each cell type by construction, the neuron cloud traces out the population-average tuning profile in a single trial, replacing the classical average over repeated stimulus directions. Both crowd-surrogate estimates are indicated in the upper right. **b, c**, Projection kernels used to compute DSI (**b**, *h*(Δθ) = cos(Δθ)) and OSI (**c**, *h*(Δθ) = cos(2 Δθ)). Each surrogate is the dot product of normalized firing rates with the corresponding kernel, divided by the mean normalized rate *D*_*b,c*_ (see Methods).

**Extended Data Figure 13:**
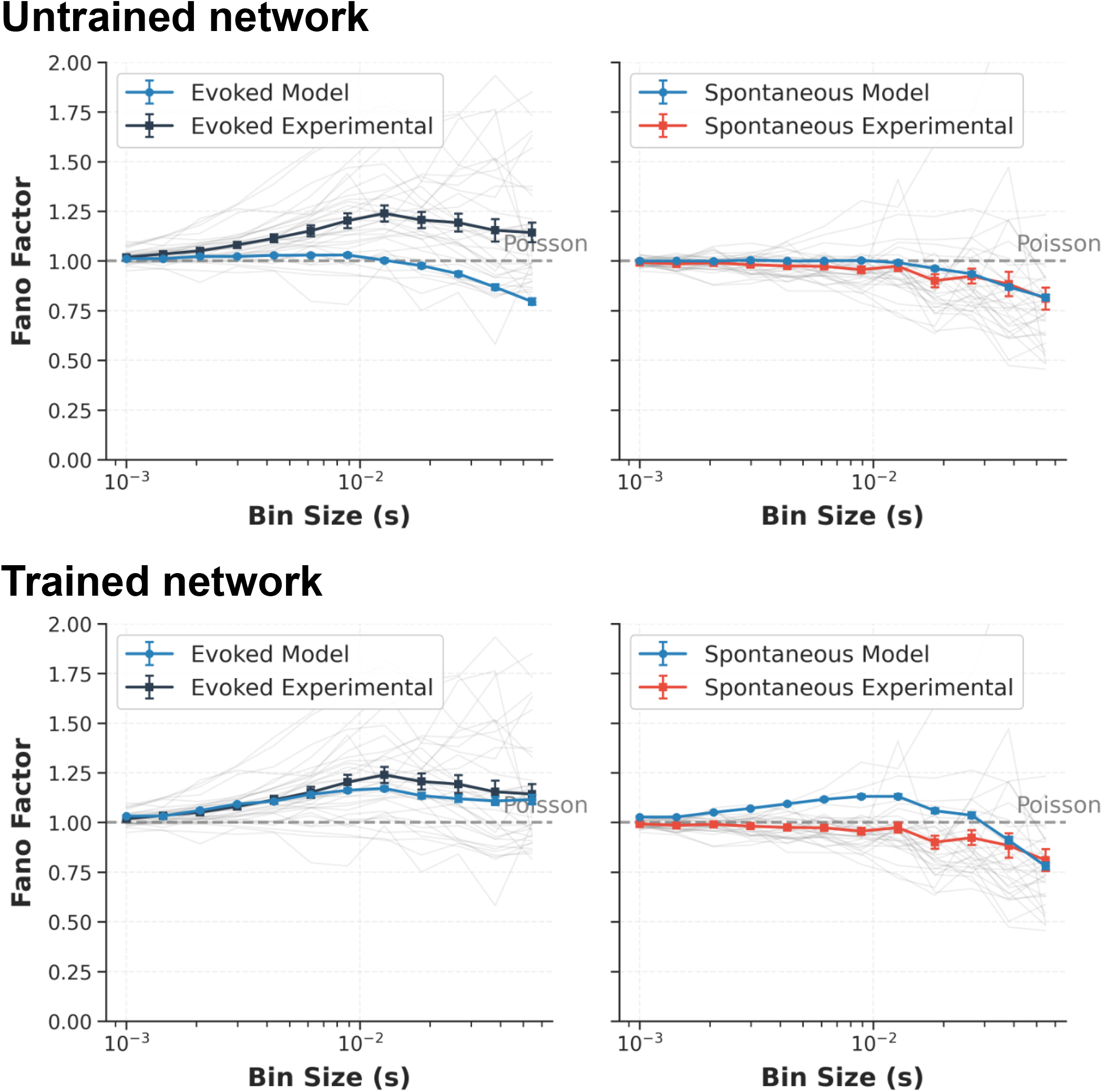
Multi-scale Fano factors. Fano factors computed across multiple bin sizes for evoked (left) and spontaneous (right) activity. The model results are shown for a single representative trained network (blue line; mean *±* SEM from multiple pools of neurons). The experimental results are derived from the Brain Observatory Neuropixels dataset^21^. Black and red lines indicate the mean and SEM of 32 mice; thin gray lines indicate individual mice.

**Extended Data Figure 14:**
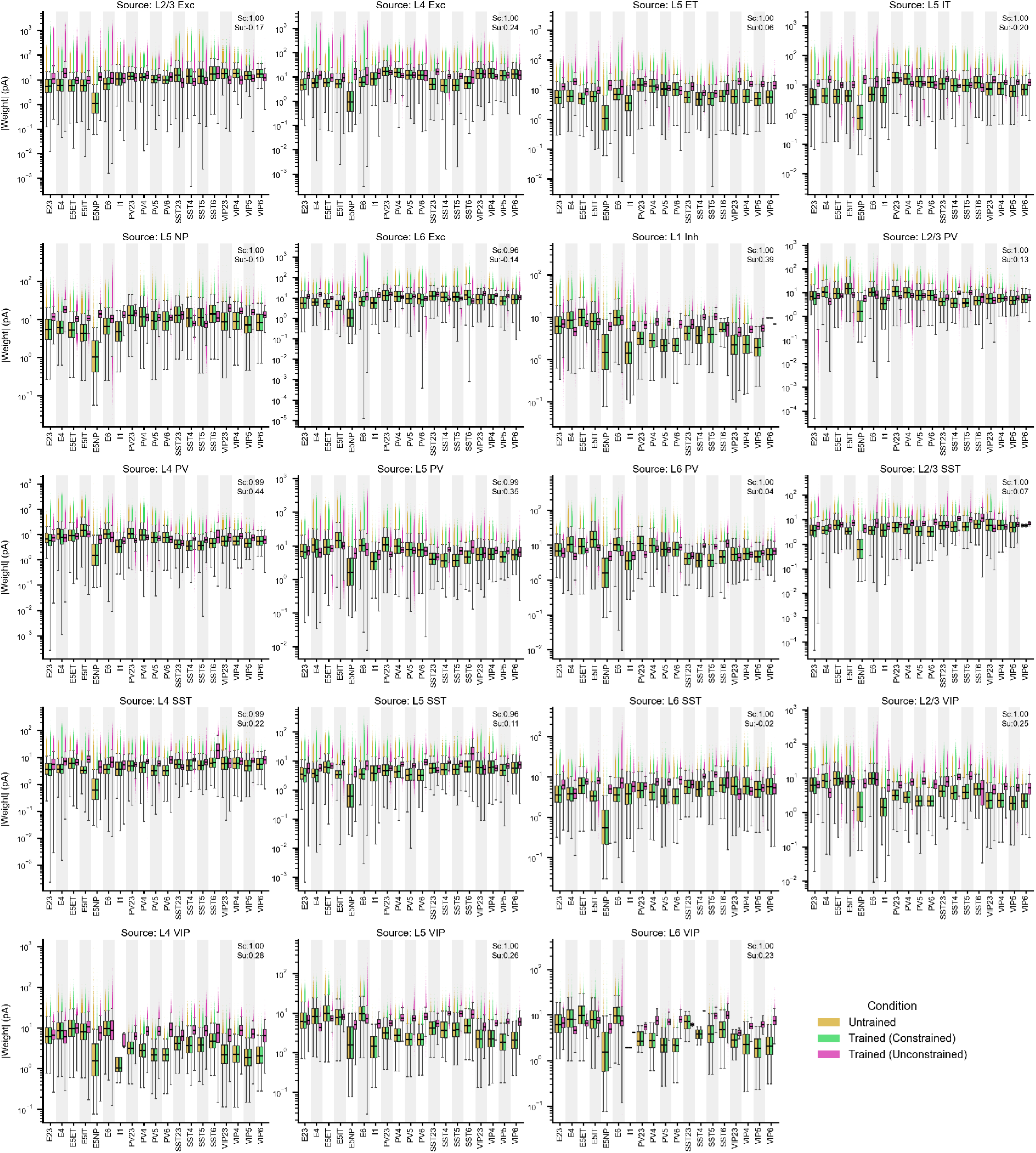
Recurrent synaptic weight distributions across training conditions. Each panel shows absolute synaptic weight distributions (note log scale) for connections from a given source cell type onto each of the 19 target cell types (x-axis), aggregated across *N* = 10 networks. Three conditions are shown: Untrained (yellow), Trained—Constrained (green, trained with synaptic weight distributions constrained to match experimentally measured values), and Trained—Unconstrained (pink, trained without this constraint). Because the weight distributions in the Untrained condition are set up according to the experimental data^19^, they serve as regularization targets during training—when training employs such constraints (that is, the Trained—Constrained case). *S*_*c*_ and *S*_*u*_ indicate the similarity score (1 − *D*_KS_, defined in Methods) of weights between the Untrained and each trained condition, respectively.

## References

1. Gerstner, W., Sprekeler, H., & Deco, G. “Theory and Simulation in Neuroscience”. Science 338, 60–65 (2012).

2. D’Angelo, E., Solinas, S., Garrido, J., Casellato, C., Pedrocchi, A., Mapelli, J., et al. “Realistic modeling of neurons and networks: towards brain simulation”. Functional Neurology 28, 153–166 (2013).

3. Einevoll, G. T., Destexhe, A., Diesmann, M., Grün, S., Jirsa, V., de Kamps, M., et al. “The Scientific Case for Brain Simulations”. Neuron 102, 735–744 (2019).

4. D’Angelo, E. & Jirsa, V. “The quest for multiscale brain modeling”. Trends in Neurosciences 45, 777–790 (2022).

5. Haufler, D., Ito, S., Koch, C., & Arkhipov, A. “Simulations of cortical networks using spatially extended conductance-based neuronal models”. The Journal of Physiology 601, 3123–3139 (2023).

6. Zanichelli, N., Schons, M., Freeman, I., Shiu, P., & Arkhipov, A. “State of Brain Emulation Report 2025”. arXiv preprint arXiv:2510.15745 (2025).

7. Arkhipov, A., da Costa, N., de Vries, S., Bakken, T., Bennett, C., Bernard, A., et al. “Integrating multimodal data to understand cortical circuit architecture and function”. Nature Neuroscience 28, 717–730 (2025).

8. Prinz, A. A., Bucher, D., & Marder, E. “Similar network activity from disparate circuit parameters”. Nature Neuroscience 7, 1345–1352 (2004).

9. Goaillard, J.-M., Taylor, A. L., Schulz, D. J., & Marder, E. “Functional consequences of animal-to-animal variation in circuit parameters”. Nature Neuroscience 12, 1424–1430 (2009).

10. Eriksson, O., Bhalla, U. S., Blackwell, K. T., Crook, S. M., Keller, D., Kramer, A., et al. “Combining hypothesis- and data-driven neuroscience modeling in FAIR workflows”. eLife 11, e69013 (2022).

11. Gouwens, N. W., Berg, J., Feng, D., Sorensen, S. A., Zeng, H., Hawrylycz, M. J., et al. “Systematic generation of biophysically detailed models for diverse cortical neuron types”. Nature Communications 9, 710 (2018).

12. Tasic, B., Yao, Z., Graybuck, L. T., Smith, K. A., Nguyen, T. N., Bertagnolli, D., et al. “Shared and distinct transcriptomic cell types across neocortical areas”. Nature 563, 72–78 (2018).

13. Teeter, C., Iyer, R., Menon, V., Gouwens, N., Feng, D., Berg, J., et al. “Generalized leaky integrate-and-fire models classify multiple neuron types”. Nature Communications 9, 709 (2018).

14. Gouwens, N. W., Sorensen, S. A., Berg, J., Lee, C., Jarsky, T., Ting, J., et al. “Classification of electrophysiological and morphological neuron types in the mouse visual cortex”. Nature Neuroscience 22, 1182–1195 (2019).

15. Gouwens, N. W., Sorensen, S. A., Baftizadeh, F., Budzillo, A., Lee, B. R., Jarsky, T., et al. “Integrated Morphoelectric and Transcriptomic Classification of Cortical GABAergic Cells”. Cell 183, 935–953.e19 (2020).

16. Scala, F., Kobak, D., Bernabucci, M., Bernaerts, Y., Cadwell, C. R., Castro, J. R., et al. “Phenotypic variation of transcriptomic cell types in mouse motor cortex”. Nature 598, 144–150 (2021).

17. Yao, Z., van Velthoven, C. T. J., Kunst, M., Zhang, M., McMillen, D., Lee, C., et al. “A high-resolution transcriptomic and spatial atlas of cell types in the whole mouse brain”. Nature 624, 317–332 (2023).

18. Seeman, S. C., Campagnola, L., Davoudian, P. A., Hoggarth, A., Hage, T. A., Bosma-Moody, A., et al. “Sparse recurrent excitatory connectivity in the microcircuit of the adult mouse and human cortex”. eLife 7, e37349 (2018).

19. Campagnola, L., Seeman, S. C., Chartrand, T., Kim, L., Hoggarth, A., Gamlin, C., et al. “Local connectivity and synaptic dynamics in mouse and human neocortex”. Science 375, eabj5861 (2022).

20. De Vries, S. E. J., Lecoq, J. A., Buice, M. A., Groblewski, P. A., Ocker, G. K., Oliver, M., et al. “A largescale standardized physiological survey reveals functional organization of the mouse visual cortex”. Nature Neuroscience 23, 138–151 (2020).

21. Siegle, J. H., Jia, X., Durand, S., Gale, S., Bennett, C., Graddis, N., et al. “Survey of spiking in the mouse visual system reveals functional hierarchy”. Nature 592, 86–92 (2021).

22. The MICrONS Consortium. “Functional connectomics spanning multiple areas of mouse visual cortex”. Nature 640, 435–447 (2025).

23. Billeh, Y. N., Cai, B., Gratiy, S. L., Dai, K., Iyer, R., Gouwens, N. W., et al. “Systematic Integration of Structural and Functional Data into Multi-scale Models of Mouse Primary Visual Cortex”. Neuron 106, 388–403.e18 (2020).

24. Rimehaug, A. E., Stasik, A. J., Hagen, E., Billeh, Y. N., Siegle, J. H., Dai, K., et al. “Uncovering circuit mechanisms of current sinks and sources with biophysical simulations of primary visual cortex”. eLife 12, e87169 (2023).

25. Galván Fraile, J., Scherr, F., Ramasco, J. J., Arkhipov, A., Maass, W., & Mirasso, C. R. “Modeling circuit mechanisms of opposing cortical responses to visual flow perturbations”. PLOS Computational Biology 20, e1011921 (2024).

26. Rimehaug, A. E., Dale, A. M., Arkhipov, A., & Einevoll, G. T. “Uncovering population contributions to the extracellular potential in the mouse visual system using Laminar Population Analysis”. PLOS Computational Biology 20, e1011830 (2024).

27. Bodor, A. L., Schneider-Mizell, C. M., Zhang, C., Elabbady, L., Mallen, A., Bergeson, A., et al. “The synaptic architecture of layer 5 thick tufted excitatory neurons in mouse visual cortex”. Nature Neuroscience 28, 1704–1715 (2025).

28. Chen, G., Scherr, F., & Maass, W. “A data-based largescale model for primary visual cortex enables brain-like robust and versatile visual processing”. Science Advances 8, eabq7592 (2022).

29. Wang, C., Zhang, T., Chen, X., He, S., Li, S., & Wu, S. “BrainPy, a flexible, integrative, efficient, and extensible framework for general-purpose brain dynamics programming”. eLife 12, e86365 (2023).

30. Deistler, M., Kadhim, K. L., Pals, M., Beck, J., Huang, Z., Gloeckler, M., et al. “JAXLEY: differentiable simulation enables large-scale training of detailed biophysical models of neural dynamics”. Nature Methods 22, 2649–2657 (2025).

31. Allen Institute. V1 Deep Dive dataset. Data available at https://github.com/AllenInstitute/v1dd_physiology. 2024.

32. Ko, H., Hofer, S. B., Pichler, B., Buchanan, K. A., Sjöström, P. J., & Mrsic-Flogel, T. D. “Functional specificity of local synaptic connections in neocortical networks”. Nature 473, 87–91 (2011).

33. Cossell, L., Iacaruso, M. F., Muir, D. R., Houlton, R., Sader, E. N., Ko, H., et al. “Functional organization of excitatory synaptic strength in primary visual cortex”. Nature 518, 399–403 (2015).

34. Ding, Z., Fahey, P. G., Papadopoulos, S., Wang, E. Y., Celii, B., Papadopoulos, C., et al. “Functional connectomics reveals general wiring rule in mouse visual cortex”. Nature 640, 459–469 (2025).

35. Dai, K., Hernando, J., Billeh, Y. N., Gratiy, S. L., Planas, J., Davison, A. P., et al. “The SONATA data format for efficient description of large-scale network models”. PLoS Computational Biology 16, e1007696 (2020).

36. Rossi, L. F., Harris, K. D., & Carandini, M. “Spatial connectivity matches direction selectivity in visual cortex”. Nature 588, 648–652 (2020).

37. Dai, K., Gratiy, S. L., Billeh, Y. N., Xu, R., Cai, B., Cain, N., et al. “Brain Modeling ToolKit: An open source software suite for multiscale modeling of brain circuits”. PLoS Computational Biology 16, e1008386 (2020).

38. Durand, S., Iyer, R., Mizuseki, K., de Vries, S., Mihalas, S., & Reid, R. C. “A comparison of visual response properties in the lateral geniculate nucleus and primary visual cortex of awake and anesthetized mice”. Journal of Neuroscience 36, 12144–12156 (2016).

39. Cuntz, H., Bird, A. D., Mittag, M., Beining, M., Schneider, M., Mediavilla, L., et al. “A general principle of dendritic constancy: A neuron’s size-and shapeinvariant excitability”. Neuron 109, 3647–3662 (2021).

40. Song, S., Sjöström, P. J., Reigl, M., Nelson, S., & Chklovskii, D. B. “Highly nonrandom features of synaptic connectivity in local cortical circuits”. PLoS Biology 3, e68 (2005).

41. Buzsáki, G. & Mizuseki, K. “The log-dynamic brain: how skewed distributions affect network operations”. Nature Reviews Neuroscience 15, 264–278 (2014).

42. Markram, H., Muller, E., Ramaswamy, S., Reimann, M. W., Abdellah, M., Sanchez, C. A., et al. “Reconstruction and Simulation of Neocortical Microcircuitry”. Cell 163, 456–492 (2015).

43. Reimann, M. W., Gevaert, M., Shi, Y., Lu, H., Markram, H., & Muller, E. “A null model of the mouse whole-neocortex micro-connectome”. Nature communications 10, 3903 (2019).

44. Dura-Bernal, S., Herrera, B., Lupascu, C., Marsh, B. M., Gandolfi, D., Marasco, A., et al. “Large-Scale Mechanistic Models of Brain Circuits with Biophysically and Morphologically Detailed Neurons”. Journal of Neuroscience 44 (2024).

45. Nicola, W. & Clopath, C. “Supervised learning in spiking neural networks with FORCE training”. Nature communications 8, 2208 (2017).

46. Huh, D. & Sejnowski, T. J. Gradient Descent for Spiking Neural Networks. in Advances in Neural Information Processing Systems (eds Bengio, S., Wallach, H., Larochelle, H., Grauman, K., Cesa-Bianchi, N., & Garnett, R.) 31 (Curran Associates, Inc., 2018).

47. Bellec, G., Salaj, D., Subramoney, A., Legenstein, R., & Maass, W. Long short-term memory and Learning-to-learn in networks of spiking neurons. in Advances in Neural Information Processing Systems 31 (2018).

48. Bellec, G., Scherr, F., Subramoney, A., Hajek, E., Salaj, D., Legenstein, R., et al. “A solution to the learning dilemma for recurrent networks of spiking neurons”. Nature Communications 11, 3625 (2020).

49. Strata, P. & Harvey, R. “Dale’s principle”. Brain Research Bulletin 50, 349–350 (1999).

50. Neftci, E. O., Mostafa, H., & Zenke, F. “Surrogate Gradient Learning in Spiking Neural Networks: Bringing the Power of Gradient-Based Optimization to Spiking Neural Networks”. IEEE Signal Processing Magazine 36, 51–63 (2019).

51. Zenke, F. & Vogels, T. P. “The Remarkable Robustness of Surrogate Gradient Learning for Instilling Complex Function in Spiking Neural Networks”. Neural Computation 33, 899–925 (2021).

52. Kingma, D. P. & Ba, J. “Adam: A Method for Stochastic Optimization”. arXiv preprint arXiv:1412.6980 (2014).

53. Nemirovskij, A. S. & Yudin, D. B. Problem Complexity and Method Efficiency in Optimization. A Wiley-Interscience publication; translated by E. R. Dawson, 388. ISBN: 9780471103455 (John Wiley & Sons, Chichester and New York, 1983).

54. Surace, S. C., Pfister, J.-P., Gerstner, W., & Brea, J. “On the choice of metric in gradient-based theories of brain function”. PLOS Computational Biology 16, e1007640 (2020).

55. Pogodin, R., Cornford, J., Ghosh, A., Gidel, G., Lajoie, G., & Richards, B. “Synaptic Weight Distributions Depend on the Geometry of Plasticity”. arXiv preprint arXiv:2305.19394 (2023).

56. Cornford, J., Pogodin, R., Ghosh, A., Sheng, K., Bicknell, B. A., Codol, O., et al. “Brain-like learning with exponentiated gradients”. bioRxiv (2024).

57. Micikevicius, P., Narang, S., Alben, J., Diamos, G., Elsen, E., Garcia, D., et al. “Mixed precision training”. arXiv preprint arXiv:1710.03740 (2017).

58. Chen, T., Xu, B., Zhang, C., & Guestrin, C. “Training deep nets with sublinear memory cost”. arXiv preprint arXiv:1604.06174 (2016).

59. Lappalainen, J. K., Tschopp, F. D., Prakhya, S., McGill, M., Nern, A., Shinomiya, K., et al. “Connectomeconstrained networks predict neural activity across the fly visual system”. Nature 634, 1132–1140 (2024).

60. Niell, C. M. & Stryker, M. P. “Highly Selective Receptive Fields in Mouse Visual Cortex”. Journal of Neuro-science 28, 7520–7536 (2008).

61. Fu, Y., Tucciarone, J. M., Espinosa, J. S., Sheng, N., Darcy, D. P., Nicoll, R. A., et al. “A Cortical Circuit for Gain Control by Behavioral State”. Cell 156, 1139–1152 (2014).

62. Niell, C. M. & Scanziani, M. “How Cortical Circuits Implement Cortical Computations: Mouse Visual Cortex as a Model”. Annual Review of Neuroscience 44, 517–546 (2021).

63. Ayaz, A., Saleem, A. B., Schölvinck, M. L., & Carandini, M. “Locomotion controls spatial integration in mouse visual cortex”. Current Biology 23, 890–894 (2013).

64. Jordan, R. & Keller, G. B. “Opposing Influence of Top-down and Bottom-up Input on Excitatory Layer 2/3 Neurons in Mouse Primary Visual Cortex”. Neuron 108, 1194–1206.e5 (2020).

65. Keller, A. J., Roth, M. M., & Scanziani, M. “Feedback generates a second receptive field in neurons of the visual cortex”. Nature 582, 545–549 (2020).

66. Stringer, C., Pachitariu, M., Steinmetz, N., Reddy, C. B., Carandini, M., & Harris, K. D. “Spontaneous behaviors drive multidimensional, brainwide activity”. Science 364, eaav7893 (2019).

67. Kim, C. M. & Chow, C. C. “Training Spiking Neural Networks in the Strong Coupling Regime”. Neural computation 33, 1199–1233 (2021).

68. Herranz-Celotti, L. & Rouat, J. “Stabilizing Spiking Neuron Training”. arXiv preprint arXiv:2202.00282 (2024).

69. Millman, D. J., Ocker, G. K., Caldejon, S., Kato, I., Larkin, J. D., Lee, E. K., et al. “VIP interneurons in mouse primary visual cortex selectively enhance responses to weak but specific stimuli”. eLife 9, e55130 (2020).

70. Walker, E. Y., Sinz, F. H., Cobos, E., Muhammad, T., Froudarakis, E., Fahey, P. G., et al. “Inception loops discover what excites neurons most using deep predictive models”. Nature Neuroscience 22, 2060–2065 (2019).

71. Wertz, A., Trenholm, S., Yonehara, K., Hillier, D., Raics, Z., Leinweber, M., et al. “Single-cell–initiated monosynaptic tracing reveals layer-specific cortical network modules”. Science 349, 70–74 (2015).

72. Kuan, A. T., Bondanelli, G., Driscoll, L. N., Han, J., Kim, M., Hildebrand, D. G. C., et al. “Synaptic wiring motifs in posterior parietal cortex support decisionmaking”. Nature 627, 367–373 (2024).

73. Znamenskiy, P., Kim, M.-H., Muir, D. R., Iacaruso, M. F., Hofer, S. B., & Mrsic-Flogel, T. D. “Functional specificity of recurrent inhibition in visual cortex”. Neuron 112, 991–1000.e8 (2024).

74. Ogando, M. B., Abdeladim, L., Sit, K. K., Shin, H., Sridharan, S., Gopakumar, K., et al. “Feature-specific inhibitory connectivity augments the accuracy of cortical representations”. bioRxiv, 2025.08.02.668307 (2025).

75. Lee, W.-C. A., Bonin, V., Reed, M., Graham, B. J., Hood, G., Glattfelder, K., et al. “Anatomy and function of an excitatory network in the visual cortex”. Nature 532, 370–374 (2016).

76. Loewenstein, Y., Kuras, A., & Rumpel, S. “Multiplicative dynamics underlie the emergence of the lognormal distribution of spine sizes in the neocortex in vivo”. The Journal of Neuroscience: The Official Journal of the Society for Neuroscience 31, 9481–9488 (2011).

77. Gal, E., London, M., Globerson, A., Ramaswamy, S., Reimann, M. W., Muller, E., et al. “Rich cell-typespecific network topology in neocortical microcircuitry”. Nature Neuroscience 20, 1004–1013 (2017).

78. Bonifazi, P., Goldin, M., Picardo, M. A., Jorquera, I., Cattani, A., Bianconi, G., et al. “GABAergic hub neurons orchestrate synchrony in developing hippocampal networks”. Science (New York, N.Y.) 326, 1419–1424 (2009).

79. Luccioli, S., Angulo-Garcia, D., Cossart, R., Malvache, A., Módol, L., Sousa, V. H., et al. “Modeling driver cells in developing neuronal networks”. PLOS Computational Biology 14, e1006551 (2018).

80. Sanzeni, A., Akitake, B., Goldbach, H. C., Leedy, C. E., Brunel, N., & Histed, M. H. “Inhibition stabilization is a widespread property of cortical networks”. eLife 9, e54875 (2020).

81. Mahrach, A., Chen, G., Li, N., van Vreeswijk, C., & Hansel, D. “Mechanisms underlying the response of mouse cortical networks to optogenetic manipulation”. eLife 9, e49967 (2020).

82. Fink, A. J. P., Muscinelli, S. P., Wang, S., Hogan, M. I., English, D. F., Axel, R., et al. “Experience-dependent reorganization of inhibitory neuron synaptic connectivity”. bioRxiv (2025).

83. White, J. G., Southgate, E., Thomson, J. N., & Brenner, S. “The structure of the nervous system of the nematode Caenorhabditis elegans”. Philosophical Transactions of the Royal Society of London. B, Biological Sciences 314, 1–340 (1986).

84. Cook, S. J., Jarrell, T. A., Brittin, C. A., Wang, Y., Bloniarz, A. E., Yakovlev, M. A., et al. “Whole-animal connectomes of both Caenorhabditis elegans sexes”. Nature 571, 63–71 (2019).

85. Winding, M., Pedigo, B. D., Barnes, C. L., Patsolic, H. G., Park, Y., Kazimiers, T., et al. “The connectome of an insect brain”. Science 379, eadd9330 (2023).

86. Dorkenwald, S., Matsliah, A., Sterling, A. R., Schlegel, P., Yu, S.-c., McKellar, C. E., et al. “Neuronal wiring diagram of an adult brain”. Nature 634, 124–138 (2024).

87. Jun, J. J., Steinmetz, N. A., Siegle, J. H., Denman, D. J., Bauza, M., Barbarits, B., et al. “Fully integrated silicon probes for high-density recording of neural activity”. Nature 551, 232–236 (2017).

88. Potjans, T. C. & Diesmann, M. “The Cell-Type Specific Cortical Microcircuit: Relating Structure and Activity in a Full-Scale Spiking Network Model”. Cerebral Cortex 24, 785–806 (2014).

89. Tsodyks, M. V., Skaggs, W. E., Sejnowski, T. J., & McNaughton, B. L. “Paradoxical effects of external modulation of inhibitory interneurons”. The Journal of Neuroscience 17, 4382–4388 (1997).

90. Pfeffer, C. K., Xue, M., He, M., Huang, Z. J., & Scanziani, M. “Inhibition of inhibition in visual cortex: the logic of connections between molecularly distinct interneurons”. Nature Neuroscience 16, 1068–1076 (2013).

91. Pi, H.-J., Hangya, B., Kvitsiani, D., Sanders, J. I., Huang, Z. J., & Kepecs, A. “Cortical interneurons that specialize in disinhibitory control”. Nature 503, 521–524 (2013).

92. Marder, E. & Taylor, A. L. “Multiple models to capture the variability in biological neurons and networks”. Nature Neuroscience 14, 133–138 (2011).

93. Kriegeskorte, N. & Douglas, P. K. “Cognitive computational neuroscience”. Nature Neuroscience 21, 1148–1160 (2018).

94. Randi, F., Sharma, A. K., Dvali, S., & Leifer, A. M. “Neural signal propagation atlas of Caenorhabditis elegans”. Nature 623, 406–414 (2023).

95. Dura-Bernal, S., Neymotin, S. A., Suter, B. A., Dacre, J., Moreira, J. V. S., Urdapilleta, E., et al. “Multiscale model of primary motor cortex circuits predicts in vivo cell-type-specific, behavioral state-dependent dynamics”. Cell Reports 42 (2023).

96. Shiu, P. K., Sterne, G. R., Spiller, N., Franconville, R., Sandoval, A., Zhou, J., et al. “A Drosophila computational brain model reveals sensorimotor processing”. Nature 634, 210–219 (2024).

97. Plesser, H. E., Davison, A. P., Diesmann, M., Fukai, T., Gemmeke, T., Gleeson, P., et al. “Building on models—a perspective for computational neuroscience”. Cerebral Cortex 35, bhaf295 (2025).

98. Igarashi, J., Yamaura, H., & Yamazaki, T. “Large-Scale Simulation of a Layered Cortical Sheet of Spiking Network Model Using a Tile Partitioning Method”. Frontiers in Neuroinformatics 13, 71 (2019).

99. Lu, W., Du, X., Wang, J., Zeng, L., Ye, L., Xiang, S., et al. “Simulation and assimilation of the digital human brain”. Nature Computational Science 4, 890–898 (2024).

100. Kuriyama, R., Akira, K., Green, L., Herrera, B., Dai, K., Iura, M., et al. Microscopic-Level Mouse Whole Cortex Simulation Composed of 9 Million Biophysical Neurons and 26 Billion Synapses on the Supercomputer Fugaku. in (Association for Computing Machinery, 2025), 2158–2171.

## Methods References

101. Lee, S., Hjerling-Leffler, J., Zagha, E., Fishell, G., & Rudy, B. “The Largest Group of Superficial Neocortical GABAergic Interneurons Expresses Ionotropic Serotonin Receptors”. The Journal of Neuroscience 30, 16796–16808 (2010).

102. Yao, Z., van Velthoven, C. T. J., Nguyen, T. N., Goldy, J., Sedeno-Cortes, A. E., Baftizadeh, F., et al. “A taxonomy of transcriptomic cell types across the isocortex and hippocampal formation”. Cell 184, 3222–3241.e26 (2021).

103. Lien, A. D. & Scanziani, M. “Cortical direction selectivity emerges at convergence of thalamic synapses”. Nature 558, 80–86 (2018).

104. Arkhipov, A., Gouwens, N. W., Billeh, Y. N., Gratiy, S., Iyer, R., Wei, Z., et al. “Visual physiology of the layer 4 cortical circuit in silico”. PLoS computational biology 14, e1006535 (2018).

105. Simon, H. A. “On a class of skew distribution functions”. Biometrika 42, 425–440 (1955).

106. Pachón, A., Polito, F., & Sacerdote, L. “Random Graphs Associated to some Discrete and Continuous Time Preferential Attachment Models”. Journal of Statistical Physics 162, 1608–1638 (2016).

107. Guo, Y., Chen, Y., Zhang, L., Wang, Y., Liu, X., Tong, X., et al. Reducing information loss for spiking neural networks. in European Conference on Computer Vision (2022), 36–52.

108. Rotter, S. & Diesmann, M. “Exact digital simulation of time-invariant linear systems with applications to neuronal modeling”. Biological cybernetics 81, 381–402 (1999).

109. Galván Fraile, J. Neocortical dynamics and computational mechanisms: Integrating sensory and higher-order inputs. Available at https://dspace.uib.es/xmlui/handle/11201/173343. PhD thesis (Universitat de les Illes Balears, 2025).

110. Abbott, L. F. & Nelson, S. B. “Synaptic plasticity: taming the beast”. Nature Neuroscience 3, 1178–1183 (2000).

111. Fernandez-Hart, T., Knight, J. C., & Kalganova, T. “Posit and floating-point based Izhikevich neuron: A Comparison of arithmetic”. Neurocomputing 597,

112. Werbos, P. J. “Backpropagation through time: what it does and how to do it”. Proceedings of the IEEE 78, 1550–1560 (1990).

113. Lillicrap, T. P. & Santoro, A. “Backpropagation through time and the brain”. Current opinion in neuro-biology 55, 82–89 (2019).

114. Abadi, M., Agarwal, A., Barham, P., Brevdo, E., Chen, Z., Citro, C., et al. TensorFlow: Large-Scale Machine Learning on Heterogeneous Systems. Software available from: https://www.tensorflow.org/. 2015.

115. Eshraghian, J. K., Ward, M., Neftci, E. O., Wang, X., Lenz, G., Dwivedi, G., et al. “Training spiking neural networks using lessons from deep learning”. Proceedings of the IEEE (2023).

116. Dabney, W., Rowland, M., Bellemare, M., & Munos, R. Distributional reinforcement learning with quantile regression. in Proceedings of the AAAI conference on artificial intelligence 32 (2018).

117. Scherr, F. & Maass, W. “Analysis of the computational strategy of a detailed laminar cortical microcircuit model for solving the image-change-detection task”. bioRxiv, 2021–11 (2021).

118. Buluç, A., Fineman, J. T., Frigo, M., Gilbert, J. R., & Leiserson, C. E. Parallel sparse matrix-vector and matrix-transpose-vector multiplication using compressed sparse blocks. in Proceedings of the twenty-first annual symposium on Parallelism in algorithms and architectures (2009), 233–244.

119. Katz, L. C. & Shatz, C. J. “Synaptic activity and the construction of cortical circuits”. Science 274, 1133–1138 (1996).

120. Hensch, T. K. “Critical period plasticity in local cortical circuits”. Nature Reviews Neuroscience 6, 877–888 (2005).

121. Espinosa, J. S. & Stryker, M. P. “Development and plasticity of the primary visual cortex”. Neuron 75, 230–249 (2012).

122. Niell, C. M. & Stryker, M. P. “Modulation of Visual Responses by Behavioral State in Mouse Visual Cortex”. Neuron 65, 472–479 (2010).

123. Engel, T. A., Steinmetz, N. A., Gieselmann, M. A., Thiele, A., Moore, T., & Boahen, K. “Selective modulation of cortical state during spatial attention”. Science 354, 1140–1144 (2016).

124. Gewaltig, M.-O. & Diesmann, M. “NEST (NEural Simulation Tool)”. Scholarpedia 2, 1430 (2007).

125. Villamar, J., Vogelsang, J., Linssen, C., Kunkel, S., Kurth, A., Schöfmann, C. M., et al. NEST 3.6. 2023. 127903 (2024).

